# A major disease-related point mutation in spastin alters dramatically the dynamics and allostery of the motor

**DOI:** 10.1101/2024.10.17.618886

**Authors:** Shehani Kahawatte, Amanda C. Macke, Carter St. Clair, Ruxandra I. Dima

## Abstract

Spastin is a microtubule-severing AAA+ ATPase that is highly expressed in neuronal cells and plays a crucial role in axonal growth, branching, and regeneration. This machine oligomerizes into hexamers in the presence of ATP and the microtubule carboxy-terminal tails (CTTs). Conformational changes in spastin hexamers, powered by ATP hydrolysis, apply forces on the microtubule, ultimately leading to the severing of the filament. Mutations disrupt the normal function of spastin, impairing its ability to sever microtubules effectively and leading to abnormal microtubule dynamics in neurons characteristic for the set of neurodegenerative disorders called hereditary spastic paraplegias (HSP). Experimental studies have identified the HSP-related R591S (*Drosophila melanogaster* numbering) mutation as playing a crucial role in spastin. Given its significant role in HSP, we employed a combination of molecular dynamics simulations with machine learning and graph network based approaches to identify and quantify the perturbations caused by the R591S HSP mutation on spastin’s dynamics and allostery with functional implications. We found that the functional hexamer, upon the HSP-related mutation, loses the ability to execute the primary motion associated with the severing action. The study of allosteric changes upon the mutation showed that the regions that are most perturbed are those involved in the formation of the inter-protomer contacts. The mutation induces rigidity in the allosteric networks of the motor making it more likely to experience loss of function as any applied perturbations could not be easily dissipated by passing through a variety of alternative paths as in the wild-type (WT) spastin.

## 1. INTRODUCTION

Microtubules (MT) are dynamic, cylindrical polymers that form an integral part of the eukaryotic cytoskeleton. Composed of *α* and *β* tubulin subunits, these versatile structures play critical roles in maintaining cellular architecture, facilitating intracellular transport, and ensuring proper chromosome segregation during cell division.^1,2^ A key regulatory mechanism of MT dynamics is MT severing, a process that modulates the length of MTs. This function is mediated by specialized MT-associated proteins (MAPs) known as the MT-severing enzymes. These enzymes, which belong to the AAA+ (ATPases Associated with diverse cellular Activities) superfamily, harness the energy from ATP hydrolysis to generate mechanical forces that induce breaks within the MT lattice.^3–5^ MT severing enzymes are pivotal in various cellular processes, including spindle formation during mitosis, and ciliogenesis, underscoring their importance in cellular function and organization.^1,6–9^

Spastin is a MT-severing machine that plays a crucial role in axonal growth, branching, and regeneration.^1,10^ The AAA+ domain of spastin, which is primarily responsible for MT severing, is divided into two subdomains: the nucleotide-binding domain (NBD) and the helical bundle domain (HBD). The NBD contains several highly conserved functional regions within AAA+ proteins: the Walker-A (WA), Walker-B (WB), and Arg-finger motifs, all of which are essential for ATP binding and hydrolysis, as well as the pore loop regions, specifically PL1 and PL2, which play a critical role in substrate binding.^1,11,12^

Studies revealed that spastin exists as a monomer at cellular concentrations but oligomerizes into hexamers in the presence of ATP and the MT carboxy-terminal tails (CTTs), which project from the MT surface.^1,11,13^ Cryo-electron microscopy (cryo-EM) has identified two distinct conformations of these hexamers, namely spiral (PDB ID: 6P07) and ring (PDB ID:6PEN), both resolved in complex with a glutamate-rich peptide substrate.^12,14^ In the spiral conformation, from *Drosophila melanogaster*, the protomers are organized in a right-handed spiral, with a gap of approximately 35 °A between the terminal protomers, while all the nucleotide-binding pockets of the protomers are occupied by ATP.^11,12^ Upon ATP hydrolysis, protomer F loses its ATP and becomes highly flexible, causing the gap between the terminal protomers to close and transition into the ring conformation.^14^ Conformational changes, powered by ATP hydrolysis, apply forces on the MT lattice, ultimately leading to its severing. Our previous work indicated that the terminal protomers A and F, which undergo significant conformational changes, play an active role in driving the mechanical action, while the inner protomers provide stability to the hexamer and assist in gripping the substrate, ensuring efficient severing.^2–5,15,16^

In the hexameric structures of spastin, the interactions between protomers B through E follow a canonical convex-to-concave pattern, where the convex interface of protomer i interacts with both the NBD and HBD of protomer i-1, and the concave interface of protomer i interacts only with the NBD of protomer i+1.^8,11,12^ Structural studies showed that PL1 and PL2 form a tight double-helical staircase around the substrate for binding and creating inter-protomer contacts within the spastin hexamer.^12^ Although both PL1 and PL2 are considered essential for severing activity, PL1 has been identified as crucial for recognizing and binding the substrate, whereas PL2 facilitates communication of this action to the rest of the protein.^11,15–17^ For example, in our most recent study on spastin’s allostery we found that PL2 is a crucial region for transmitting allosteric signals within the NBD.^16^ The other important regions that facilitate the oligomerization of spastin include PL3 in the NBD and the CT-Helix (CT-Hlx) in the HBD. PL3 is recognized as crucial for transmitting allosteric signals both within the NBD and from the NBD to the HBD, while the CT-Hlx is a long helix that wraps over the HBD from the convex to the concave interface of a spastin monomer.^12,16,17^ Effective allosteric communication among these regions ensures that spastin retains its oligomeric structure, which is crucial for its cellular functions.

Hereditary spastic paraplegias (HSP) are a group of neurodegenerative disorders characterized by progressive weakness and spasticity in the lower limbs, primarily caused by mutations in spastin.^11,12,18,19^ These mutations disrupt the normal function of spastin, impairing its ability to sever MTs effectively and leading to abnormal MT dynamics in neurons. The resulting accumulation of damaged MTs can cause axonal degeneration and neuronal dysfunction, contributing to the clinical symptoms of HSP.^18–21^ To date, researchers have identified over 200 spastin mutations linked to HSP, with approximately 30% classified as missense mutations.^18^ Notably, most of these missense mutations cluster within spastin’s AAA+ ATPase domain, and have diverse effects on spastin’s activity, including disruption of oligomerization, impairment of nucleotide binding, and destabilization of the AAA+ domain structure.^12^ Despite the availability of genetic information, analyses relevant to understanding the molecular mechanisms by which these mutations cause alterations in spastin’s activity are limited. Therefore, detailed studies focusing on the specific effects of spastin mutations associated with HSP are crucial for uncovering the disorder’s underlying causes and for developing effective targeted therapies.

In this work, we conducted a detailed analysis of the role of the R591S (*D. melanogaster* numbering as in the solved cryo-EM structure of the spastin spiral^12^) HSP mutation, located near the PL2 region, on spastin’s dynamics and allostery with functional implications. Experimental studies have identified R591 as very important in spastin, being positioned at the center of an allosteric network that couples substrate-binding pore loops to ATP hydrolysis and oligomerization.^12^ Moreover, this residue is perfectly conserved in the meiotic subclade, underscoring its critical role in spastin.^11,16,18^ Our recent study also identified R591 as a hub, demonstrating its structural importance in the protein, and as a central residue in several protomers, highlighting its significance in the allosteric communication pathways.^16^ Given its significant role in spastin function, we aim to identify the perturbations caused by the R591S HSP mutation on spastin’s functionality.

Following literature findings,^22^ to decipher the allosteric sites/regions in the R591S mutant we employed a multi-pronged approach, which we previously used to dissect the allostery of the WT spastin.^16^ Our combination of molecular dynamics simulations with machine learning and graph network based approaches revealed substantial changes in both the dynamics of the tertiary and quaternary forms of the protein as well as in their allostery, with direct implications for the proper function of this molecular motor. This is the first time the effects of an HSP-related mutation on the functional regions of spastin have been characterized at the molecular level.

## 2. Materials and Methods

### 2.1 System Preparation & Simulation Details

#### 2.1.1 Starting Structures

The initial structure of the spastin hexamer was derived from the original cryo-EM structure of spastin in its spiral conformation^12^ from the Protein Data Bank^23^ (PDB) ID: 6P07), with the missing residues modeled using the Modeller program.^15^ Utilizing this structure, we prepared the mutant spastin by mutating residue R591 to S591 in all the protomers within the spiral hexamer using the PyMOL software (version 2.5).^24^ Following our previous work, we simulated both systems in the presence of ATP and a minimal substrate (E15), referred to as the COMPLEX state, as well as in the absence of them, referred to as the APO state.^15^ To decipher the tertiary contribution from the quaternary ensemble, we performed the mutation on a monomer, following our previous work.^16,17^ For both the WT and the R591S mutated monomeric systems, two additional states, the substrate (SUB) and nucleotide (NUC) states were constructed apart from the COMPLEX and APO states. These states allowed us to study the effect of the mutation on binding each partner.^17^

#### 2.1.2 Simulation Details

The simulations for the COMPLEX and APO states of the WT spastin hexamer were obtained from our previous work,^15,16^ whereas new simulations were performed for the R591S mutant systems. These simulations utilized all-atomistic molecular dynamic (MD) simulations with the GROMACS^25^ molecular modeling software version 2022 and the GROMOS96 54a7 force field.^26^ Based on this force field, the automated topology builder server was used to construct force field parameters for ATP.^15^ Each system was positioned at the center of a dodecahedral box, with dimensions approximately 150x150x150 Å^3^. The solvation was carried out using the explicit solvent model of single point charge (SPC), and neutralized with NaCl ions. We also applied three-dimensional periodic boundary conditions (PBC) to the systems. Additional information regarding the R591S mutant hexamer setups is listed in Table S1. For energy minimization and to eliminate steric overlaps, we used the Verlet cutoff technique and the steepest descent algorithm for 50,000 steps, with the convergence criteria of the maximal force value being less than 23.9006 kcal/mol/Å (1000 kJ/mol/nm).^27^ Next, the systems were equilibrated first with the NVT ensemble to raise the temperature to 300K, using a velocity-rescaling thermostat and the leapfrog integrator algorithm for 500 ps.^28^ We then used the NPT ensemble to maintain the system’s pressure at 1.0 bar for 500 ps using the Parrinello-Rahman pressure coupling scheme, again with the leapfrog integrator algorithm.^29^ To constrain the bond lengths involving hydrogen atoms, we used the LINCS approach which allowed us to use a longer integration step of 2 fs^30^ and to determine the electrostatic interactions we utilized the Particle Mesh Ewald (PME) algorithm.^15,31^ For each system, at least five production MD simulations were performed for a minimum of 200 ns (see Table S1).

The same procedures were used to perform the monomeric simulations for both the WT and R591S mutant states. For the WT monomeric states, we started from the simulations from our previous work^17^ and extended each simulation up to 200 ns. For the R591S mutant monomeric states we ran new simulations, with each state having at least three MD trajectories for a minimum of 200 ns. Further details regarding the monomer simulations can be found in Tables S2 and S3. It is important to note that in the monomer setups, over the extended time-scale, the minimal substrate was found to dissociate in the SUB state for the WT as well as in the SUB and COMPLEX states for the R591S systems. Therefore, for the mentioned setups, truncated simulations were used, as discussed in the Results section.

### 2.2 Data Analysis

#### 2.2.1 Root Mean Square Deviation (RMSD)

To evaluate the convergence of our simulations and the overall stability of each system, we calculated the root mean-square deviation (RMSD) of the protein backbone for each MD simulation, with respect to the initial structure, using GROMACS. Tables S4 and S5 report the global average RMSD values used to compare the stability of each state in the WT and R591S mutant systems. The individual RMSD plots for the described monomer simulations are depicted in Figure S1. Figures S2a and Figure S2c display the RMSD plots for the R591S mutant hexameric simulations, while the RMSD plots for the WT hexameric simulations are available in our previous work (Figures S1c2 and S1c4 of Reference^15^).

#### 2.2.2 Dynamic Cross Correlation Matrix (DCCM) Convergence

To further verify the convergence of our MD simulations, we computed the Dynamic Cross Correlation Matrix (DCCM) for each system using the MDTraj python library.^32^ This matrix illustrates the time-dependent correlations between residue pairs based on Equation S1. To evaluate the convergence of the DCCM matrix for each state, we calculated the mean square distance R(t) between the correlation values of residue pairs at successive time intervals, according to Equation S2.^15,33^ As the DCCM matrix converges, R(t) → 0. Figure S2 illustrates the DCCM convergence for each state of the R591S mutant hexamer. Note that, when evaluating R(t) for each state, we concatenated the data obtained from each trajectory of that specific state up to time t.

#### 2.2.3 RMSD based Double Clustering Analysis

A two-step RMSD-based clustering analysis was performed on the WT and the R591S mutant hexameric systems to evaluate the structural changes in the spastin spiral conformation caused by the mutation.^34^ First, we concatenated all the MD simulations per state (APO and COMPLEX) from the WT and the mutant systems separately and then we performed the initial clustering on these concatenated trajectories using GROMACS (gmx cluster) with a RMSD cutoff of 0.14 nm.^34–36^ The resulting clustering centers were then clustered to identify similar groups. To determine an appropriate RMSD cutoff for the second clustering step, we tested various cutoffs between 0.15 nm and 0.5 nm. We chose a cutoff that ensured at least 70% of the central structures from the first clustering step were included within the top two clusters in the second clustering step for each state of both the WT and the mutant.^34^ With this criteria, a cutoff of 0.45 nm was chosen (details about the cutoff evaluation are provided in Table S6). Since the top 2 clusters cover at least 70% of the conformations, we considered these clusters as representative of the simulations. The central structures of the top two clusters obtained from the second clustering step were extracted for further analysis (cluster1:Figure 1a and cluster2:Figure S3a). For each system (WT and R591S mutant), the central structure of cluster1 for the APO state and the COMPLEX state were aligned, and the RMSD values of functional regions were calculated using the Visual Molecular Dynamics (VMD).^37^ This process was repeated for cluster 2. We used the calculated RMSD values, depicted in Figure 1 and Figures S3-S5, to evaluate global structural changes due to the mutation upon ligand removal. For clarification, the RMSD values between the WT structures are denoted as RMSD_WT(COMP)→WT(APO)_, and the RMSD values between the mutant structures are denoted as RMSD_R591S(COMP)→R591S(APO)_.

**Figure 1:**
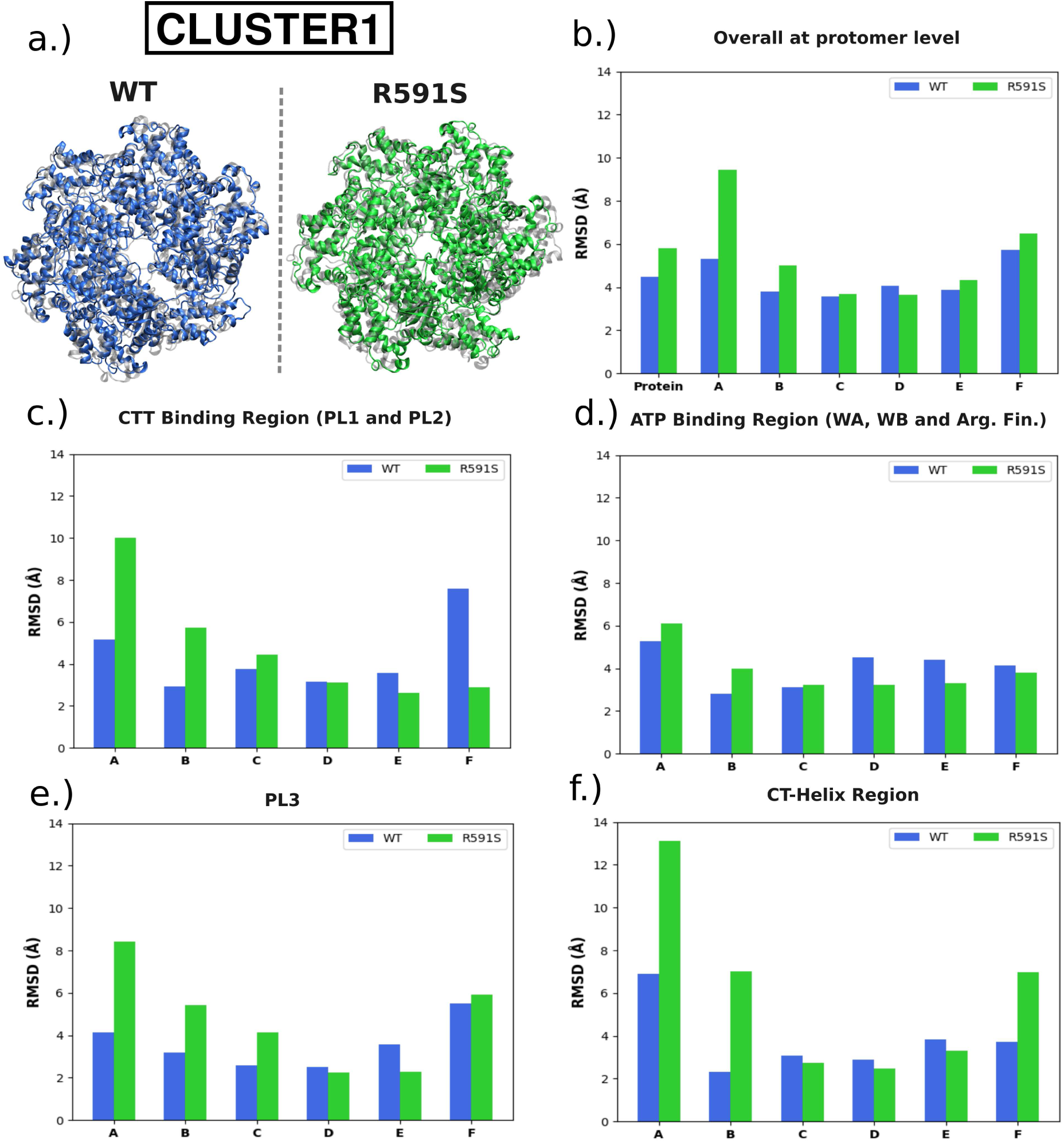
The RMSD values of the central structure of the APO state, aligned with the central structure of the COMPLEX state for cluster 1. (values were obtained by aligning the whole protein) (a) Central structures of cluster1 obtained from the second clustering step for WT and the mutant systems.(Grey indicates the COMPLEX structure, while blue and green indicate the APO structure of the WT and mutant systems, respectively) (b) Comparison of the entire protein and each protomer (c) Comparison of the CTT binding region (including both PL1 and PL2) (d) Comparison of the ATP binding region (including WA, WB and ARG fin) (e) Comparison of the PL3 region (f) Comparison of the CT-Hlx region. The residue numbers of these functional regions can be found in Table S7.

#### 2.2.4 Essential Dynamics Analysis

To determine how the R591S mutation affects the overall structure and dynamics of the motor, a detailed principal component analysis (PCA) was performed for both systems. We concatenated the trajectories and used GROMACS tools (gmx covar and gmx aneig) based on the C-*α* atoms to calculate the covariance matrix, and then extract the first two component motions^38–40^ representing the largest covered variance (see Tables S8-S10) in the MD simulations.^40–42^ We analyzed the global motions of the motor as well as the motions of the PL1 (A550 to L567) and the PL2 (S589 to F606) regions from each protomer.^12,15^ We note that, for each of the PLs, we consider five residues before and after the reported positions from the literature spanning the loop identified in the experimental structure, which in the case of PL2 includes the mutated position (R591).

To compare the essential spaces, we used the root-mean-square inner product (RMSIP) from Equation S3. We used this metric to compare the global motions between the WT and R591S mutant in each state, found in Table S11. We also visualized both their individual PC1 and PC2 subspaces, with the resulting motions represented in porcupine plots (Figure 2 and Figure S6 represent the PC1 and PC2 global motions and Figures S10-S11 represent the PL motions). We also projected the R591S mutated system onto the WT system PC space,^40,43^ which allowed us to evaluate how the behavior of the spastin severing motor changes due to the single point mutation. We calculated the free energy landscape (using gmx sham) for each of these motions to evaluate the conformational space in lieu of the known appropriate collective variable.^15,22,44–46^ Here, we focused on changes in the number of minima, rather than their specific locations, and extracted the representative conformations that reflect the changes in the landscape (see Figures S7-S9). We also provide a comparison of each minima to the starting structure using the backbone RMSD in Tables S12-S14.

**Figure 2:**
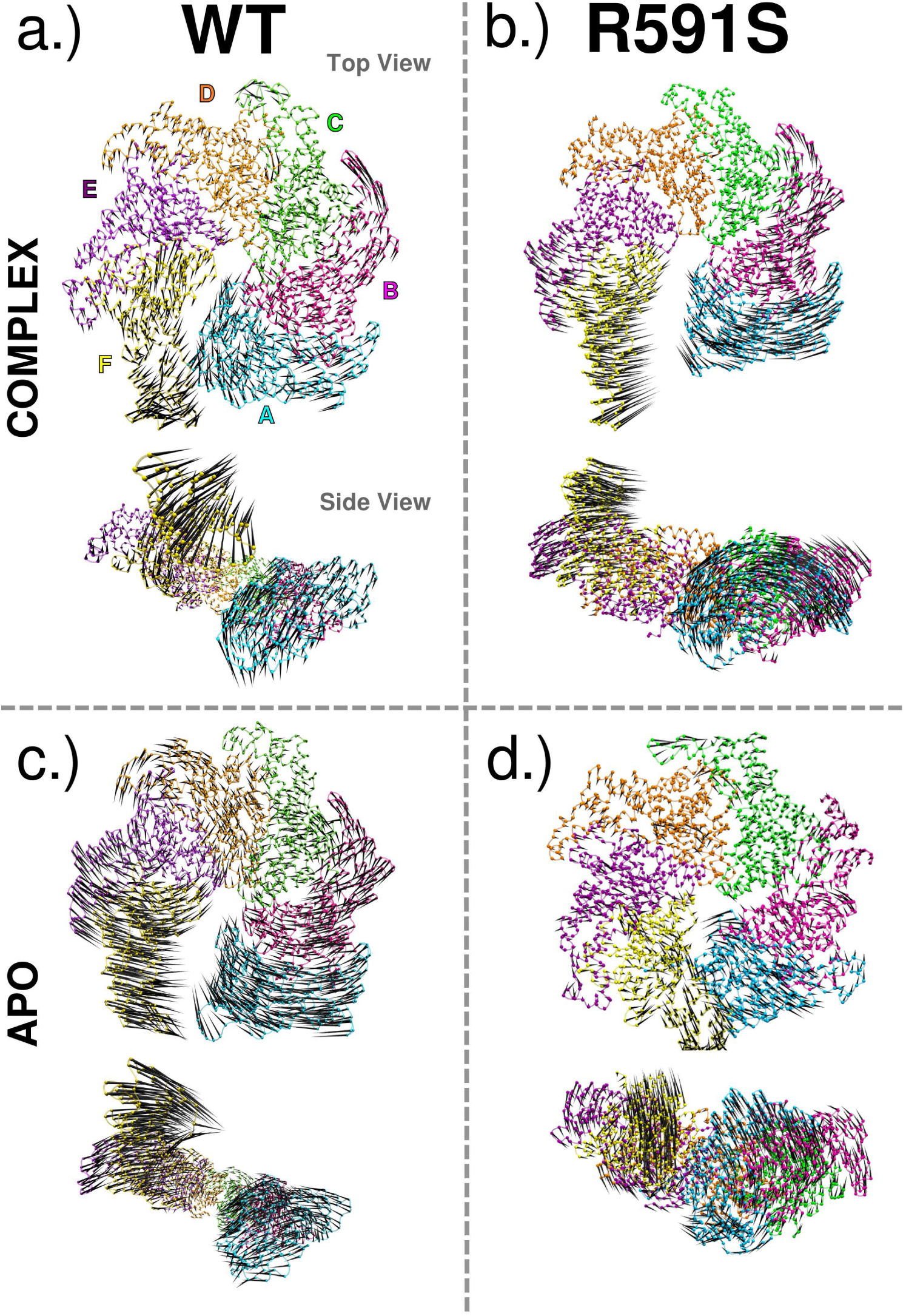
The porcupine plots illustrate the global motions corresponding to PC1 (a) global motions of WT COMPLEX state (b) global motions of R591S COMPLEX state (c) global motions of WT APO state (d) global motions of R591S APO state.The protomer labels, as indicated in (a), are used consistently across the entire figure. A side view of the protein is also included to clearly display the motions of the terminal protomers, A and F. The variance covered by PC1 for each of these systems is in Table S8.

#### 2.2.5 Salt Bridge Analysis

Salt bridges have been shown to play a crucial role in stabilizing the structure and dynamics of spastin.^15^ To understand the effect of the R591S mutation on the salt bridge networks in spastin, we conducted a salt bridge analysis using the VMD software.^37^ We analyzed intra protomer salt bridges for each protomer and inter protomer salt bridges for each pair of protomers, except between the terminal protomers A and F. Salt bridges are characterized by a maximum distance of 4 Å between positively charged (ARG or LYS) and negatively charged (ASP or GLU) residues. We selected only those salt bridges that were present for at least 1 ns in the simulation trajectories, following our previous work.^15^ We then calculated the average persistence time for each selected salt bridge and determined the average persistence time across the chains if the salt bridge was found to be present for more than 10 ns in at least three protomers. Tables S16–S18 list the average persistence times of the inter and intra protomer salt bridges that are present in at least three protomers for longer than 10 ns for the various states of the WT and the R591S mutant systems.

#### 2.2.6 Allostery with Machine Learning

It is crucial to understand the allostery of a protein as it provides insights into how proteins regulate their functions in response to various events. We employed classification machine learning techniques in combination with the trajectories from the MD simulations^47^ to (i) identify the allosteric regions affected by the mutation in the COMPLEX and the APO states separately; (ii) identify the allosteric regions affected by the mutation during the transition from COMPLEX state to APO state (i.e due to the removal of both the ligands), following our previous work.^16^ Note that this analysis was conducted only for the hexamer, to identify alterations in the allosteric regions for the quaternary structure of spastin. This method involved a two-step approach: 1) extract relevant descriptors and 2) classify the states.

##### Descriptor Extraction

For both the APO and COMPLEX states of the mutant system, we sampled the MD simulation trajectories every 200 ps and calculated various biochemically relevant descriptors that could potentially identify the allosteric regions.^48^ These descriptors were extracted at the level of secondary structures, with 33 secondary structures per protomer, following the same criteria for secondary structure naming used in our previous work for WT spastin^16^ (The list of the secondary structure names can be found in Table S2 of Reference^16^). The extracted biochemically relevant descriptors listed in Table 1 were chosen based on their significance in identifying the most important allosteric regions upon ligand removal in WT spastin, as found in our previous work.^16^

**Table 1:** Description of the extracted descriptors.

| Descriptor ID | Description |
| --- | --- |
| RSA | The ratio of solvent accessibility of each residue X to the maximum possible accessibility in a Gly-X-Gly tripeptide. |
| CONTACTS | The fraction of contacts at the interface between monomers with respect to the first frame of each trajectory. (A contact is defined as a Cα located within neighboring monomers found within 8 Å of any Cα within the secondary structure being analyzed.) |
| VDW | Van der Waals energy associated with the interaction between a secondary structure and the remainder of the protein (kJ/mol). |
| COULOMB | Coulombic energy associated with the interaction between a secondary structure and the remainder of the protein (kJ/mol). |

In this analysis, a ’feature’ is defined as the descriptor values for each frame from each trajectory corresponding to a specific secondary structure. For the RSA and energetic descriptors, we had 198 features with respect to all secondary structures for the six protomers of the hexamer. However, for the CONTACTS descriptor, we retained only the features that had at least six initial contacts with neighboring protomers. To calculate the descriptors values for each secondary structure, we used GROMACS, VMD, and/or Python following our previous work.^16,25,37^

##### Classification of the states

To achieve the first goal of this analysis, we merged the datasets of the WT and mutant systems for each descriptor and performed classification to distinguish between the two systems. This classification was conducted separately for the COMPLEX and APO states to identify changes in allosteric responses caused by the mutation in the presence and absence of ligands. We selected XGBoost as the classification algorithm, as it was the most effective method for classifying spastin states in our previous work on WT spastin.^16^ The XGBoost model was built using the PyCaret library in Python, utilizing default parameters and cross-validation settings provided by PyCaret.^49^ The aim of using classification was to identify the key features that allowed the model to distinguish between the WT and mutant systems, specifically focusing on the allosteric regions affected by the mutation. To pinpoint these key features, we applied SHAP (SHapley Additive exPlanations) analysis.^16,50^ We listed the top 10 features that contributed the most to the model and repeated this process for all descriptors to identify the regions with significant changes in physical properties due to the mutation. SHAP analysis not only highlighted these important features but also allowed us to visualize the descriptor ranges defining the system change through the beeswarm plots. Figures S12 and S13 shows the beeswarm plots for the four descriptors corresponding to COMPLEX and APO states, respectively. In these plots, each point is colored based on how its descriptor value compares to the average (magenta: increase, blue: decrease). The right side of each plot shows the observations with positive SHAP values that helped the XGBoost model to classify the mutant system.

A similar approach was applied to achieve our second goal, which was to identify the allosteric responses caused by the mutation during the transition from the COMPLEX state to the APO state, corresponding to ligand removal. In this case, for each descriptor, we merged the datasets of the COMPLEX and APO states of the same system and used classification to differentiate between the two states. Note that when building this model, we removed features located in the nucleotide or substrate binding regions, as well as any regions within 3 Å of the respective binding sites, to avoid biasing the model with ligand binding sites. Figure S14 presents the beeswarm plots of the 4 descriptors for the transition from COMPLEX to APO in the R591S system. The right side of each plot indicates the values that helped the model to classify the APO state. In addition, we compared the top features obtained for the mutant system with those identified for the COMPLEX to APO transition in the WT system from our previous work (see Figure 7c and Figures S39c – S41c of Reference^16^).

#### 2.2.7 Allostery with Graph Network Theory

Graph network theory is an established method for tracking allostery through protein systems by evaluating their connectivity and dynamics through the use of undirected graph networks built with the NetworkX Python library.^51^ Each residue is represented by a node in the network, which is then connected via edges based on a contact map. The edges are then assigned a weight based on the type of graph network of interest such that small weights represent two nodes that are closely associated while a large weight represent two nodes that are relatively less associated. Following our previously established protocols, we first performed network analysis on the monomeric system to establish the tertiary behavior from the longer time-scale simulations, including the additional NUC and SUB states used to decipher the roles of the ATP and the CTT.^16^ Using the WT monomer as a baseline, we examined how the mutation affects the ligand binding and the overall tertiary behavior. The same analysis was then extended to the hexamer to study the quaternary behavior and how the mutation impacts the quaternary allosteric communication.

##### Dynamic Graph Networks

To capture the dynamic behavior of the MD simulations, we constructed a graph network where nodes represent residues in contact (defined by the C-alpha atoms of non-covalently bonded residues within 10 Å according to VMD measure), and edges weighted according to *w_ij_* = -log(|*c_ij_*|), where *c_ij_* are the pairwise correlations calculated from the average DCCM across the trajectories for the given systems.^22,52–54^ The appropriate *c_ij_* cutoff for each state was determined by ensuring that the largest cluster in the network contained at least 90% of the nodes for the system. For each monomeric system, the cutoff values were found to be at 0.6 or 0.5 and for the hexameric system we used the same cutoff of 0.6 as we previously found for both the COMPLEX and APO states of the WT system (shown in Table S21).^16^ Following our previous study, we used the betweenness centrality parameter (*C_B_*), calculated using Equation S4, to identify the most central nodes (top 10%) in the network.^16,53,55–58^ The most central residues of the WT and R591S mutant monomeric systems are reported in Tables S22 and S23, respectively and their mappings onto the corresponding structures are depicted in Figure S15. Additionally, the most central residues of the R591S mutant hexamer in both the COMPLEX and APO states are provided in Tables S26 and S27, and represented in Figure S19.

##### Path Analysis for Allosteric Propagation

In addition to tracking changes to the overall network, we describe changes to the propagation (flux) of allosteric signals between essential regions of interest using a detailed path analysis. Paths are computed as the sum of the weighted edges used to propagate from source to sink. The shortest path (Dijkstra’s path) describes the optimal path a signal takes through the network.^59^ Using the Yen algorithm, we also obtained the next 20,000 suboptimal paths -defined as the next shortest paths -between each pair of residues for a total of 80,000 paths per collection.^60^ From these paths we determined the relative degeneracy of each node visited by the paths between regions of interest.^22,52^ The degree of degeneracy reflects the rigidity of communication between nodes: high degeneracy indicates a requirement for specific positions, while low degeneracy suggests flexibility within the system, allowing for alternative pathways. First we track the same path captured in our previous study to identify how the allosteric center (S589/R591S) communicated with the CT-Hlx (W749/Y753) in the HBD.^16^ By comparing the same path between the WT and the R591S mutant, we can describe how the change of the allosteric center impacts inter-domain communication. The optimal paths for the WT and R591S monomeric systems are in Table S24 and for the hexameric systems are in Tables S28-S31. In order to understand how the mutation impacted the communication between the binding sites, we evaluated an additional set of paths from the ATP binding pocket (P525/T530) to the CTT binding PLs (PL2-H596/R601). This analysis allowed us to describe intra-domain allosteric communication. The optimal paths comparing the WT and R591S monomeric systems are presented in Table S25, while those for the hexameric systems are provided in Tables S32-S35. In addition, we calculated the average degeneracy per secondary structure, as done in our previous study,^16^ to compare the WT with the mutant and to identify rigid regions.

## 3. RESULTS & DISCUSSION

### 3.1 Global structural stability in spastin is affected by the R591S HSP mutation

We utilized the backbone RMSD of the protein with respect to the corresponding initial structure for the WT and the R591S mutant to evaluate the overall stability of the two systems. In the monomeric COMPLEX state, we observed that the substrate dissociated from the mutated spastin after ∼ 90 ns, whereas it remained bound to the WT spastin for the full 200 ns trajectory. In the mutant, the substrate detached first from PL2 and then from PL1, indicating that the mutation at the allosteric center R591 near PL2 compromised its ability to bind the substrate. Consequently, when the connection to PL2 was lost, PL1 alone could not maintain substrate binding, leading to detachment from the monomeric COMPLEX state of spastin. In contrast, we found that the R591S mutation has no sizable effect on the NUC state of the spastin monomer. In the SUB state of the monomer the mutant had a lower average RMSD than the WT (see Figure S1 and Table S4). Importantly, we again observed the detachment of the substrate from the monomer in both the WT and mutant. Results from our previous work suggested that the binding of the substrate to spastin is primed by ATP binding.^16^ Thus, the absence of the nucleotide in the SUB state is likely behind the substrate detachment. We conclude that the mutation has only a modest effect on the SUB state of the monomeric spastin. The R591S mutant of the monomeric APO state showed a significantly lower average RMSD compared to the other three states (see Figure S1 and Table S4). In contrast, among the WT states, APO had the highest global RMSD, making it the least stable state. Overall, considering all the states, we found that the R591S mutation led to a reduced flexibility in the spastin monomer.

Next, we evaluated the stability of the hexameric structure of spastin, which represents a major functional state during the severing cycle of the machine. Comparing the RMSDs for the R591S mutant hexamers, shown in Figure S2, with those of the WT spastin hexamers from our previous work,^15^ we observed that the COMPLEX state of R591S mutant exhibits a greater variation in RMSD values across the set of trajectories than the corresponding state of the WT (see Reference^15^ Figure S1c4). Specifically, the higher RMSD values correspond to a substantial opening of the hexamer at the interface between the terminal protomers A and F. In contrast, the APO state had a significantly lower average RMSD in the R591S mutant compared to the WT, recalling the finding in the monomer (Figure S2 and Table S5). These results suggest that the mutation reduces substantially the flexibility of both the individual protomers and of their hexameric assembly in the absence of ligands.

### 3.2 Structural changes in the spastin hexamer resulting from the removal of the ligands are enhanced by the R591S HSP mutation

To gain quantitative insights into specific structural changes of the spastin hexamer caused by the R591S mutation, we employed the RMSD-based two-step clustering analysis detailed in the Methods section. To compare the WT with the mutant, we obtained representative structures and calculated the RMSD values separately for the WT and the mutant (shown in Figures 1 and S3–S5).

In the WT system, when comparing the structural deviations of the whole protein between the COMPLEX and APO states, we observed an RMSD value of 4.5 Å for cluster 1 and 7.6 Å for cluster 2 (RMSD_WT(COMP)→WT(APO)_). This is indicative of overall structural changes due to the absence of ligands in the WT system. Importantly, upon the R591S mutation, the RMSD value of the overall protein increased to 5.8 Å (RMSD_R591S(COMP)→R591S(APO)_) in cluster 1 and to 12.1 Å in cluster 2, which indicates that the mutation has caused global structural changes in the protein, beyond the changes due to the absence of ligands observed in the WT system (see Figure 1b for cluster 1 and S3b for cluster 2).

At the protomer level, comparing the RMSD values obtained by first aligning the whole protein reveals that, in the WT system, the terminal protomers A and F have larger RMSD values than the inner protomers in the absence of ligands in both clusters (see Figure1b and S3b). This trend was also observed in the mutant, with even higher RMSD values than the WT system. Specifically, in protomer A, the RMSD value has increased to 9.4 Å in cluster 1 and to 17.9 Å in cluster 2 of the mutant, compared to 5.3 Å in cluster1 and 8.1 Å in cluster 2 of the WT. However, as shown in Figure S4a, when the protomers were aligned separately, the RMSD values of the terminal protomers in the mutant decreased compared to the values in the WT. This indicates that, upon the removal of the ligands, the internal structures of the terminal protomers have changed less in the mutant compared to the WT, recapitulating the above results. Thus, the large RMSD values obtained for the mutant system, when the whole protein was aligned first, are due to changes in the orientation of the terminal protomers relative to the rest of the system.

Next, we aimed to identify the functional regions that undergo the most significant structural changes in various protomers upon the removal of the ligands and whether or not they are affected by the mutation. First, we examined the CTT binding region (PL1 and PL2). In cluster 1, when comparing the CTT binding regions of the WT system after aligning the whole protein, we found higher RMSD values for the CTT binding regions of the terminal protomers compared to the CTT binding regions of the inner protomers. In particular, protomer F had the highest RMSD (RMSD_WT(COMP)→WT(APO)_) with a value of 7.58 Å, and protomer B had the lowest value of 2.94 Å (see Figure 1c). Following the mutation, we noticed an entirely new pattern. The RMSD value (RMSD_R591S(COMP)→R591S(APO)_) of the CTT binding region of protomer A increased substantially to 10 Å in cluster 1 and to 14.61Å in cluster 2, and the RMSD of the CTT binding region of protomer B, which had the lowest value in the WT, also increased in both the clusters. Surprisingly, we observed that the RMSD of the CTT binding region of protomer F, which was the highest in the WT, dropped to the lowest values in the hexamer as it decreased to 2.87 Å in cluster 1 and to 7.81 Å in cluster 2 upon the mutation (see Figure 1c and S3c). The same pattern was observed when the CTT binding regions were aligned separately for protomers A and F, but with lower RMSD values compared to when aligning the whole protein (Figure S4b). These results suggest that the R591S mutation has a significantly greater impact on the orientation of the CTT binding regions of the terminal protomers with respect to the spiral hexamer, but in different ways, with their flexibility increasing dramatically in A and decreasing dramatically in F, while having a minor impact on the internal structure of the CTT binding regions compared to their structure in the WT upon the removal of the ligands.

We also investigated the structural alterations in PL1 and PL2 independently, with the results being depicted in Figure S5. The results for PL1 (Figures S5a and S5b) suggest that, upon the unbinding of both ligands, the mutation prevents the loss of the internal structure of the PL1 regions from both terminal protomers A and F, while it increases significantly the orientation flexibility of PL1 from protomer A and preserves the orientation of PL1 in protomer F with respect to the hexamer. For the PL2 region (see Figures S5c and S5d), the pattern of change recalls the behavior in PL1, but the differences in values are smaller compared to those in PL1. Importantly, for the ATP binding regions, we found that they are mostly impervious to the mutation (See Figures 1d and S3d).

Next, we focused on the oligomerization regions of the hexamer: PL3 and the CT-Hlx. For the WT system, we observed that the PL3 regions of the terminal protomers exhibit higher RMSD values than those of the inner protomers in both clusters (Figures 1e and S3e). Particularly, PL3 of protomer F had the highest RMSD recalling the above results for the CTT binding region. Upon the mutation, the RMSD for PL3 of protomer A reaches 8.41 Å. However, when the PL3 regions of the terminal protomers are aligned separately, we observe a decrease in the RMSD values for both terminal protomers in the mutant system (Figure S4e). This suggests that there are significant changes in the orientation of PL3, especially for protomer A, with respect to the rest of the hexamer. Additionally, the neighboring protomers to the terminal protomers, i.e., protomers B and E, also exhibit significant structural changes in PL3.

When comparing the CT-Hlx we observed that, for the WT system, in cluster 1 protomer A exhibited the highest RMSD value and protomer B exhibited the lowest RMSD value (Figure 1f). In cluster 2 the highest RMSD value was observed for protomer F and the lowest was observed for protomer D (Figure S3f). These findings imply that the absence of ligands impacts more the CT-Hlx region of the terminal protomers, while having lesser effects on the inner protomers. Upon the mutation, similar to the results obtained for PL3, the CT-Hlx of protomer A exhibits a much higher RMSD value, indicating more deviations due to the mutation. Additionally, the CT-Hlx of protomers F and B also show higher RMSD values.

In summary, our analysis shows that the terminal protomers are more significantly im-pacted by ligand removal than the inner protomers in the WT system. Following the mutation, the orientation of the terminal protomers with respect to the rest of the hexamer is greatly affected, while there are almost no internal structural changes in each protomer. When considering the functional regions, we observed that, due to the mutation, structural changes occur noticeably in the CTT binding regions and the oligomerization regions of the terminal protomers, with negligible effects on the ATP binding regions.

### 3.3 The R591S HSP mutation induces dramatic changes to the essential dynamics of the spastin motor

We extracted further information from the conformational dynamics probed in the simulations by carrying out a principal component analysis (PCA) for each setup. We kept the first two components in the WT as they cover 75% of the variance in the COMPLEX state and 83% variance in the APO state (Table S8). For the R591S mutant, the first two PCs cover 60% of the variance in the COMPLEX and 48% in the APO states. These initial observations demonstrated a difference in the essential subspace suggesting that the presence of the mutation dramatically alters the motions of the hexameric assembly -particularly in the APO state. We found moderate overlap between the WT and R591S mutant in both the COMPLEX (0.634) and in the APO (0.644) states using the RMSIP (Table S11). To further quantify the spaces we computed the free energy landscapes (FEL) and compared the motions described by the subspaces. We first characterized the WT space for the COMPLEX and for the APO states for reference.

In the WT COMPLEX state, the motions captured in PC1, shown in the porcupine plots in Figure 2a, are characterized by opposing axial movements in the terminal protomers A and F, suggesting an opening of the gate between these protomers. PC2 motions shown in Figure S6a also indicates an opening of the gate, corresponding to the motion of the HBD of protomer F moving outward relative to protomer A. This suggests the main dynamic motions of the WT COMPLEX state correspond to the opening of the gate between the terminal protomers. In the free energy landscape, we identified three primary minima separated by relatively high-barriers (Figure S7a), corresponding to the described motions. In the APO state, we also found three minima with slightly lower barriers between them compared to the COMPLEX case. The representative structures for each minima (Figure S8a) represent similar global conformations to the COMPLEX state due to the absence of the ligands, with the exception of minimum i, which corresponds to a much more substantial opening of the gate between the terminal protomers. The motions captured in PC1, shown in the porcupine plots in Figure 2c, are characterized by rotations in the terminal protomers A and F, resulting in gate opening, but in a manner different from the COMPLEX state. The second PC characterizes unique behaviors, corresponding to axial motions of the HBD in protomer F and a loss of correlated global motions in protomer A (Figure S6c).

In the COMPLEX state, we found that the presence of the mutation caused two additional minima to appear in the landscape (Figure S7b). The barriers between the minima are much lower compared to the WT barriers suggestive of an increased conformational flexibility in the presence of the mutation. The motions observed in PC1 (Figure 2b) show a rotation in protomer A and a horizontal translation of protomer F. These motions are striking as they demonstrate how the mutation affects the global behavior of the motor. The WT motions imply an important action where, in order to exert the downward motion of severing,^2–4,16^ the terminal protomers have to move axially in opposite directions. In the mutant this behavior is no longer the most significant action in the essential dynamics suggesting that the hexamer may no longer undergo the opening of the gate between the terminal protomers that allows for the severing action.^3–5^ In the APO state of the mutant, we found a unique landscape with three main basins of attraction separated by relatively low barriers, with two basins each composed of two minima separated by shallow barriers not observed in the WT. Surprisingly, these minima are found to be less conformationally diverse in comparison to the WT as none correspond to the open hexamer conformations. The corresponding PC motions are more disordered and not even remotely similar to the dynamics captured in the WT (Figure2d). In PC1, the HBD of protomer F is executing upward motions, while in PC2 the HBD of F hardly moves at all. In addition, there are observable differences in the rotation and orientation of protomer A. This suggests a particularly dramatic change in the dynamics of the hexamer due to the mutation in the absence of the ligands.

However, it is important to note that these projections in the two PC spaces cannot provide a complete view of the differences in conformational space and the corresponding motions between the mutant and the WT because they are not using the same space. To address this, we project the coordinates of the mutant system onto the WT space (Figure S9). As mentioned, in the COMPLEX state, the WT occupies three main regions of the landscape that correspond to specific orientations of terminal protomer F that lead to the opening of the asymmetric spiral structure. When the mutant is projected onto the WT space, it populates a distinct landscape that is characterized by more shallow barriers between identified minima as well as two small regions (iii & iv) between minima ii and v that were not observed in the WT. In the PC1 and PC2 space, the mutant does not access the minima corresponding to an axial motion. This confirms that the presence of the mutation is preventing the above described required motions for severing. Similarly, the projection of the mutant onto the WT space in the APO state led to a unique landscape characterized by only two minima that correspond to two very restrained conformations that are not able to access the open hexamers observed in the WT.

Severing activity requires that the PLs of the motor bind to the MT substrate to translocate the CTT through the pore.^11,12,15^ To extrapolate on how the changes of the global motions due to the mutation may impact the severing capabilities, we extracted the PC1 and PC2 motions for PL1 and PL2 (Tables S9 and S10 and Figures S10 and S11). In the WT COMPLEX state, PC1 (Figure S10a) of PL1 is dominated by axial motions in the terminal protomers, mimicking the global behavior of the protomers. In the mutant (Figure S10b), PL1 exhibits motions corresponding to the closing of the pore, such as the horizontal motion of PL1 of protomer F towards PL1 of A. These motions also suggest a distortion of the pore that is then emphasized with the PL2 motions (Figures S10b and S10d), which match the twisting of protomers A and B discussed above (Figure S10b). This twisting motion points towards a potentially altered disassociation pathway of the dimer. In the WT APO state (Figure S11), the PC1 for the PL1 loops of the terminal protomers shows that protomer F moves laterally towards protomer A, while protomer B moves upward. Meanwhile, PL2 undergoes motions that imply an opening of the pore where protomers F and A move outward and away from each other. In the APO mutant state (Figure S11b), the PC motions are dramatically stunted in comparison to the behavior in the WT and indicate a deactivation and potentially a loss of a correlated motion of the PL1 loops in the presence of the mutation -particularly in the absence of the ligands.

### 3.4 Salt Bridge Analysis highlights the distinct effects of the R591S mutation on different ligand bound states of spastin

Salt bridges are critical for the structure and function of spastin, as they support the conformational changes necessary for ATPase activity and stabilize the protein-substrate interactions, ensuring efficient binding and severing of MTs.^12^ To understand the effect of the R591S HSP mutation on these interactions, we analyzed the salt bridges involving residues in each of the pore-loop regions (PL1 and PL2) and the overall protein. Tables S15-S18 list the salt bridge pairs that were observed for more than 10 ns and found in at least 3 protomers in our simulations for both the APO and COMPLEX states of the WT and the R591S mutant hexamers.

Focusing on the inter-protomer salt bridges formed between positions in the PL regions, in the WT both COMPLEX and APO state, there is only long lived salt bridge was D559-K555 (Table S15). We found that the formation of this salt bridge between PL1 residues of the neighboring protomers was not affected by the R591S mutation. The analysis of inter-protomer salt bridges formed between residues outside the PL1 and PL2 regions indicates that, in both the COMPLEX and APO states of the WT and mutant systems, the long-lived salt bridges are formed at the oligomerization interface of the protomers (Figure 3). Specifically, the salt bridge between E497 in the NBD region of protomer i+1 and R704 in the HBD region of protomer i had the longest persistence time, being observed for more than 100 ns in the COMPLEX state of both the WT and the mutant, and for more than 65 ns in the APO state of both the systems (Table S16). Other common inter-protomer salt bridges observed in the COMPLEX state of the WT and mutant systems were D471-K459 and E472-K459, both between positions from Helix 1, a region important for oligomerization on the concave interface of a protomer, and D697-K644, which forms between the HBD and the NBD of neighboring protomers. Additionally, we found more salt bridges in the mutant COMPLEX state, namely E597-K603, a salt bridge between regions close to PL2, and D618-K479, a salt bridge between the NBD regions of the neighboring protomers (Table S16). However, these additional salt bridges in the mutant system have shorter persistence times compared to the salt bridges present also in the WT system.

**Figure 3:**
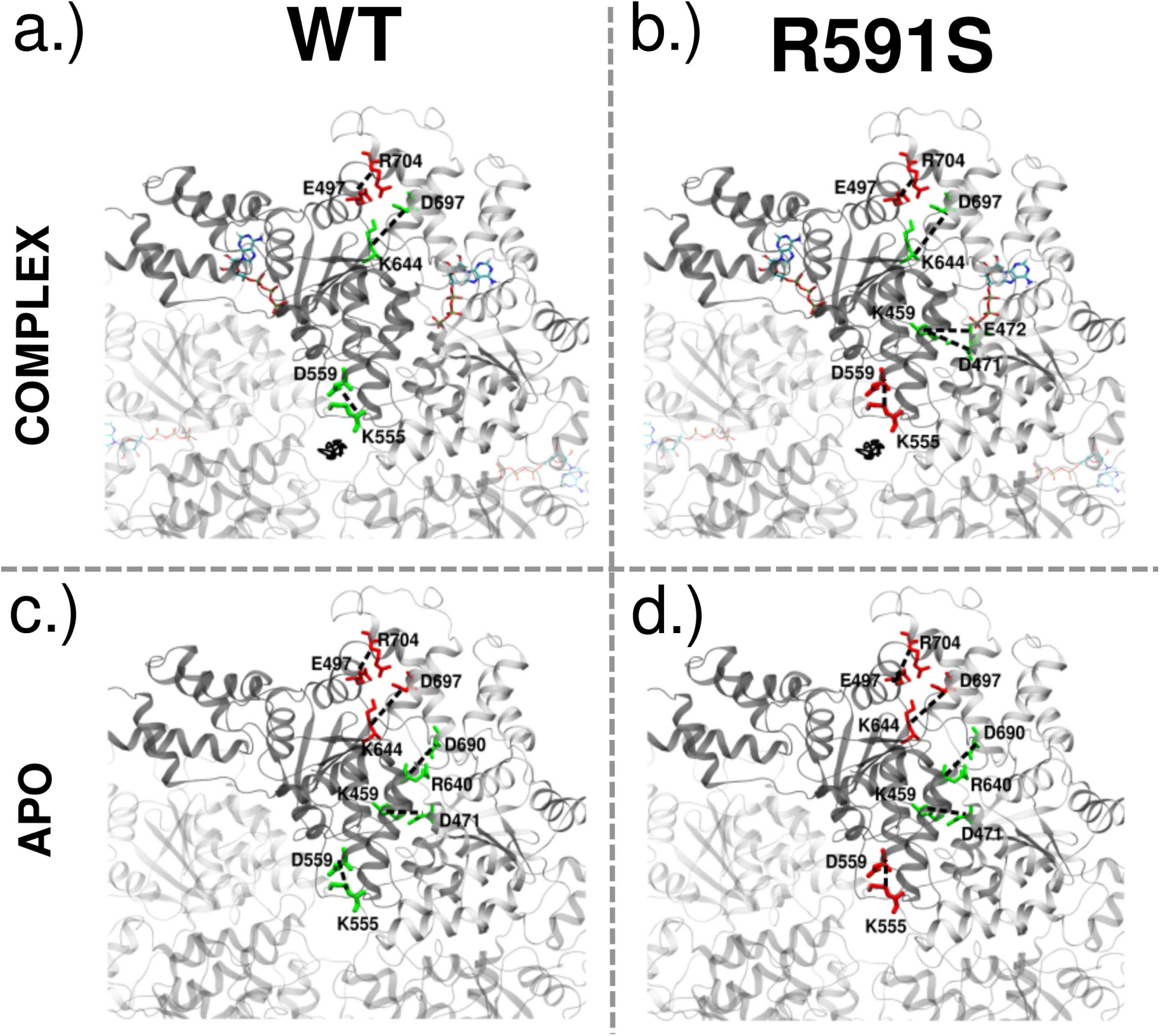
Inter-protomer salt bridges between the neighboring protomers that were observed for more than 25 ns in at least 3 protomer pairs. The salt bridges indicated in green had persistence time (t) between 25 ns and 50 ns and the salt bridges indicated in red had *t >* 50 ns (a) The COMPLEX WT system (b) The COMPLEX R591S mutated system (c) The APO WT system (d) The APO R591S mutated system. A detailed list of the salt bridges shown here is in Tables S15 and S16.

Compared to the COMPLEX state, in the APO state we found more common inter-protomer salt bridges between the WT and mutant (See Table S16). Besides the common salt bridges observed in the COMPLEX state, we also identified D618-K696 (NBD to HBD) with similar lifetimes in both systems, and D690-R640 (HBD to NBD) with a higher persistence time in the WT system. Interestingly, in the APO state of the mutant, there were no additional salt bridges, unlike the behavior of the COMPLEX state.

Focusing on the intra-protomer salt bridges, we found that they occur mainly among positions located in the HBD region and at the convex interface of the NBD region, as shown in Figure 4. Comparing the intra-protomer salt bridges in the HBD region of the COMPLEX state, we found a higher number of salt bridges in the mutant than in the WT system.

**Figure 4:**
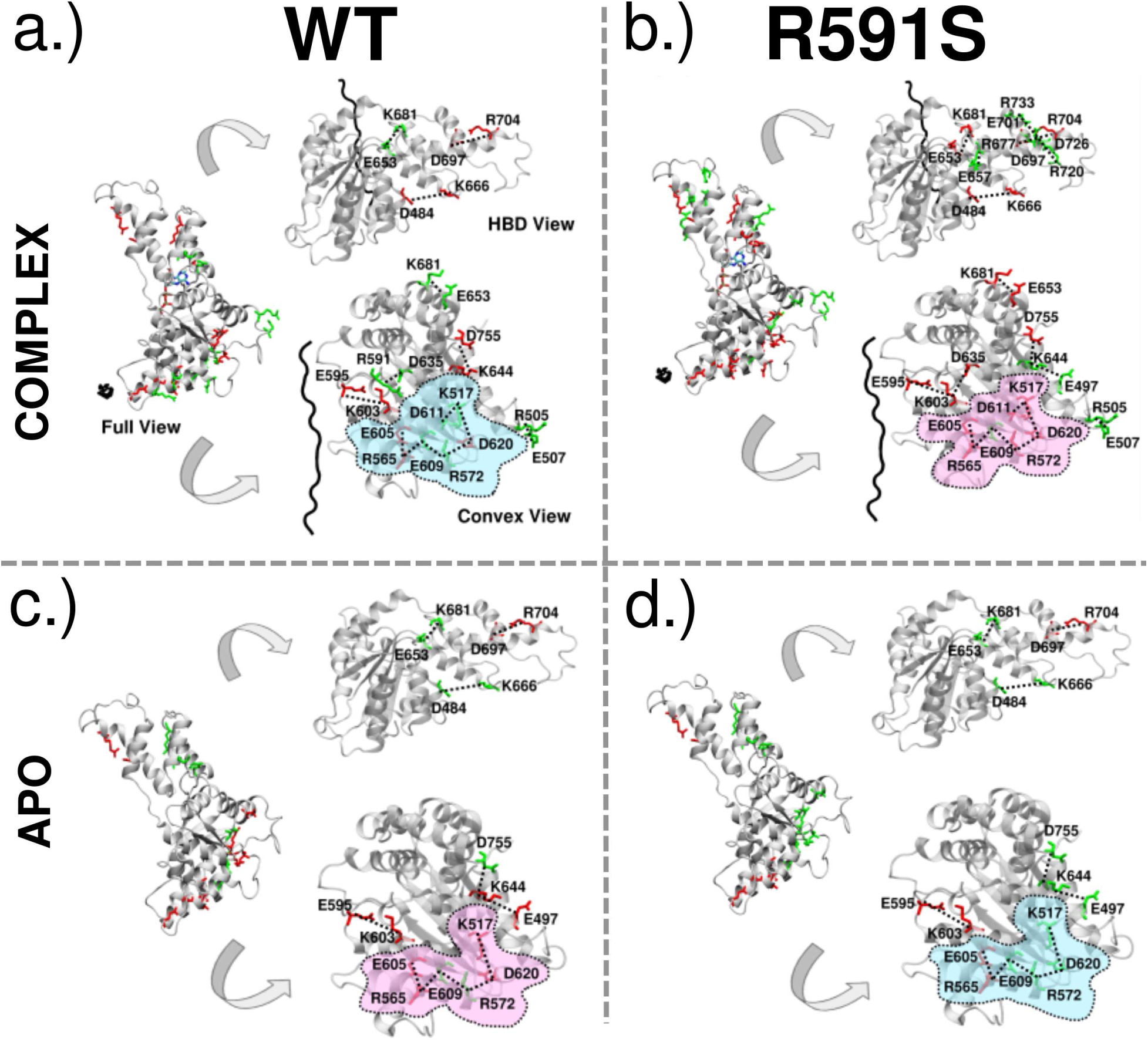
Intra-protomer salt bridges that were observed for more than 25ns in at least 3 protomers. The salt bridges indicated in green had persistence time (t) between 25ns and 50ns and the salt bridges indicated in red had *t >* 50ns (a) The COMPLEX WT system (b) The COMPLEX R591S mutated system (c) The APO WT system (d) The APO R591S mutated system. The highlighted area indicates a salt bridge network that was found to be common in all states, but with different persistence times. Blue indicates that most of the salt bridges in the network have shorter persistence times and pink indicates that most of the salt bridges in the network have longer persistence times. A detailed list of salt bridges shown here is in Table S17 and S18.

Notably, there are three salt bridges that are common between the WT and mutant systems: D484-K666, E653-K681 and D697-R704. Comparing the lifetimes of these salt bridges, all three last longer in the mutant than in the WT, unlike the above described behavior of the common inter-protomer salt bridges that had similar duration in both systems. The additional salt bridges found in the HBD region of the mutant have shorter lifetimes than the salt bridges shared with the WT, recalling the behavior in the inter-protomer COMPLEX case. In the APO state we observe only the three salt bridges common in the COMPLEX state being present in both WT and mutant systems. Interestingly, the persistence times of these salt bridges have also not been affected by the R591S mutation. This highlights the importance of these three salt bridges in the HBD region, as they are formed in both the fully functional COMPLEX state of spastin and in the APO state. It also suggests that the R591S mutation led to changes only in the HBD region in the COMPLEX state, without affecting the HBD region in the APO state, which points to the importance of the interplay between the mutation and the binding of co-factors (ATP and CTT) for the formation of long-lived salt bridges in the HBD.

Comparing the intra-protomer salt bridges in the NBD region, we identified a salt bridge network that is common to all four systems, as shown in Figure 4. This network couples residues in PL1, PL2, and in the convex interface of the protomer. In the COMPLEX state, the network has 7 members: E605 from PL2, R565 from PL1, and residues E609, R572, D620, K517, and D611 from the convex interface. The distinction between this salt bridge network in the WT and mutant systems lies solely in their lifetimes. Comparing the lifetimes, the results show that most salt bridges in the mutant system persist longer than those in the WT. In the APO state, we found the same salt bridge network but with only 6 members, excluding residue D611 observed in the COMPLEX state. However, in contrast to the findings in the COMPLEX state, we observe that in the APO state, the WT system exhibits longer lifetimes for these salt bridges compared to the mutant system.

Overall, our analysis of salt bridges highlights the impact of the R591S HSP mutation on the inter-and intra-protomer connectivity of spastin. Specifically, in the COMPLEX state we observe an increase in the number of inter-protomer salt bridges formed in the mutant, which suggests a potential structural change in response to the mutation. Conversely, in the APO state, we observed more common inter-protomer salt bridges with the WT system, indicating that the mutation has not significantly affected the connectivity between protomers of spastin when no ligands are bound. Furthermore, the intra-protomer salt bridge network in the APO state reveals that, although the network is not affected by the mutation, it has become less stable. In contrast, the network in the COMPLEX state is longer lived in the mutant.

### 3.5 The HSP mutation causes distinct allosteric responses on different ligand bound states of spastin

We extracted several biochemically relevant descriptors and applied a machine learning approach to identify the allosteric regions affected by the R591S mutation. Using the XGBoost classification model, we first analyzed separately the COMPLEX and the APO states. For each state we classified them as either the WT or the R591S mutant. This analysis helped us target the allosteric response of the COMPLEX and the APO states, respectively, to the mutation. Secondly, we classified the states of the WT as either COMPLEX or APO, and the states of the mutant as either COMPLEX or APO. This analysis helped us target the way the allosteric response of the transition between the COMPLEX and the APO states is affected by the mutation. For both types of changes we applied SHAP analysis to identify the most important allosteric regions that changed their physical properties due to the R591S mutation. Figures S12-S14 show the top 10 features for different descriptors that were identified as allosterically responding to the mutation.

We start by presenting the allosteric response of the COMPLEX state to the mutation. The loop L13 from the HBD was found to be important across all four descriptors in classifying the WT and the mutant, which indicates that L13 has a very strong allosteric response to the mutation. Another region, L16, also from the HDB and positioned along the concave interface, was important in the CONTACTS and in both energetic descriptors. Furthermore, in the COULOMB descriptor, we found that L16 was important in all chains, except for chain D, and showed increased descriptor values in all those chains due to the mutation. This suggests that the ectrostatic interactions involving L16, from the concave interface of a protomer, are destabilized due to the mutation. Furthermore, this instability may cause difficulties in forming stable native interactions between neighboring protomers, potentially impacting the overall assembly, and even the function of the hexameric complex. L3 is another important site, this time in the NBD, located along the convex interface, which was significant in three descriptors (RSA and the two energetic descriptors). Literature suggests that L3 is critical for spastin oligomerization.^12,16^ Thus, our results indicate that the mutation has allosterically-impacted two regions important for spastin oligomerization. Other regions found to exhibit an allosteric response to the mutation in at least two descriptors include H1, a helix in the NBD located at the concave interface; H5, the helix following PL1, which is one of the pore loops important for substrate binding; H10 and H11, located in the HBD and known to be important for spastin oligomerization; and L9, a region at the convex interface critical for making contacts between the NBD regions of the protomers (see Figure 5a). In summary, we found that, in the COMPLEX state, the presence of the mutation induces an allosteric response in many of the regions involved in the oligomerization of spastin.

**Figure 5:**
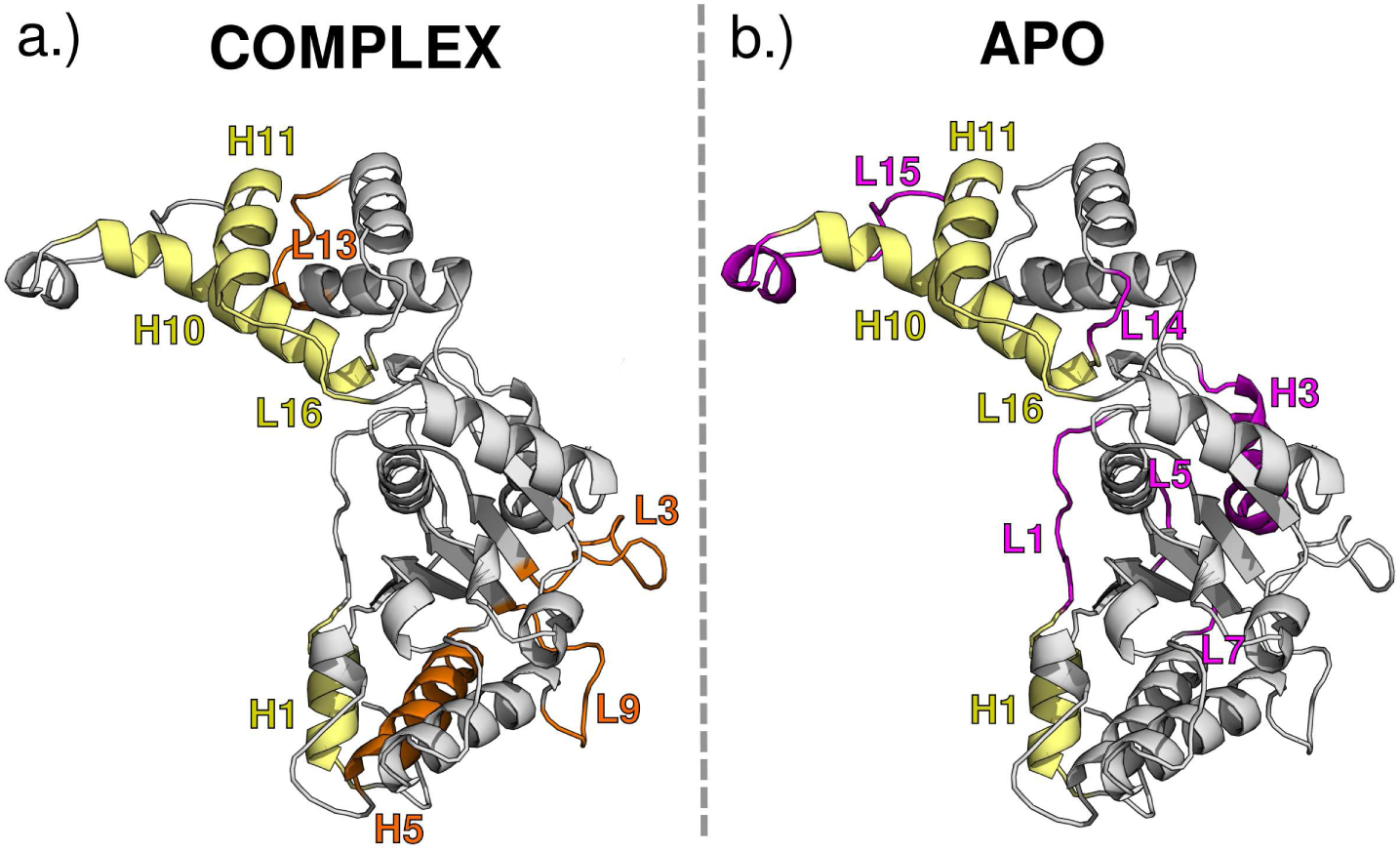
Allosteric regions that were identified as important in classifying the WT and the mutant in at least two descriptors (a) for the COMPLEX state (b) for the APO state. Regions colored in yellow were found to be important in both COMPLEX and APO states. Regions colored in orange and purple were found to be important only in the COMPLEX state and in the APO state, respectively. A detailed description of the descriptors in which these regions were found to be important is in Table S19.

Next, we focused on the allosteric response of the APO state to the mutation. In contrast to the COMPLEX state, no region was allosterically important across all the four descriptors in the APO state due to the mutation. H1, which is located at the concave face, was significant in three descriptors (CONTACTS and the two energetic descriptors) in the APO state. H3, a long helix at the convex interface of the protomer, was another notable allosteric site that was important in three descriptors in the APO state. Other regions that were important in at least two descriptors in the APO state included L1 (located in the concave interface of the protomer), L5, L7 (in the NDB), L14 (in the HBD), and L15 which includes the HBD tip, a region previously identified as important for oligomerization.^17^ Additionally, regions H10, H11, and L16, which were important in the COMPLEX state, remain allosterically significant in the APO state (see Figure 5b).

In summary, our findings reveal that, when comparing the allosterically important regions in the NBD, the affected regions in the COMPLEX state are located mainly in the lower part of the convex interface of the NBD, which plays a key role in forming the NBD-NBD contacts. In the HBD region, similar areas are impacted in both states, but in the APO state, the HBD tip, a region important for oligomerization, is particularly affected by the mutation. This suggests that the mutation allosterically impacts regions important for oligomerization in both states, but in different ways. Specifically, NBD to NBD connections between protomers are more affected in the COMPLEX state, whereas HBD to NBD connections between protomers are more affected in the APO state, especially those involving the HBD tip at the concave interface.

We next aimed to understand how the R591S mutation alters allosteric signaling between different regions upon the removal of both ligands (i.e., in the transition from the COMPLEX state to the APO state), following our previous analysis on WT spastin^16^ (the top 10 features for each descriptor for the R591S mutant system are depicted in Figure S14). Results from our previous study for the WT spastin showed that, in the CONTACTS descriptor, the regions L1, H10, and L16, located at the concave interface of the protomer, become important upon the removal of both ligands, especially in protomers B and E (See Reference^16^ Figure 7(c)). In the R591S mutant we observe a significant change in the signaling pattern, with the critical regions shifting from the concave to the convex interface, where L3 and L11 appear now as important features in the same protomers. Importantly, H10, which plays a crucial role in forming contacts between the HBD of protomer i and the NBD of protomer i+1, is no longer an important allosteric region in the mutant for chains B and E, while becoming important in chain A. In view of the crucial role played by the HBD to NBD inter-protomer contacts in stabilizing the interface between neighboring protomers^17^ these results indicate that the mutation increases the stability of the convex interfaces of chains B and E by strongly favoring the HBD to NBD inter-protomer contacts in the transition from the COMPLEX to the APO state. The signaling within the NBD region changes too, with L1 losing its significance and H1 becoming important in the mutant. Interestingly, L16, located in the HBD region, remains a significant feature in the mutant (See Figure 6).

**Figure 6:**
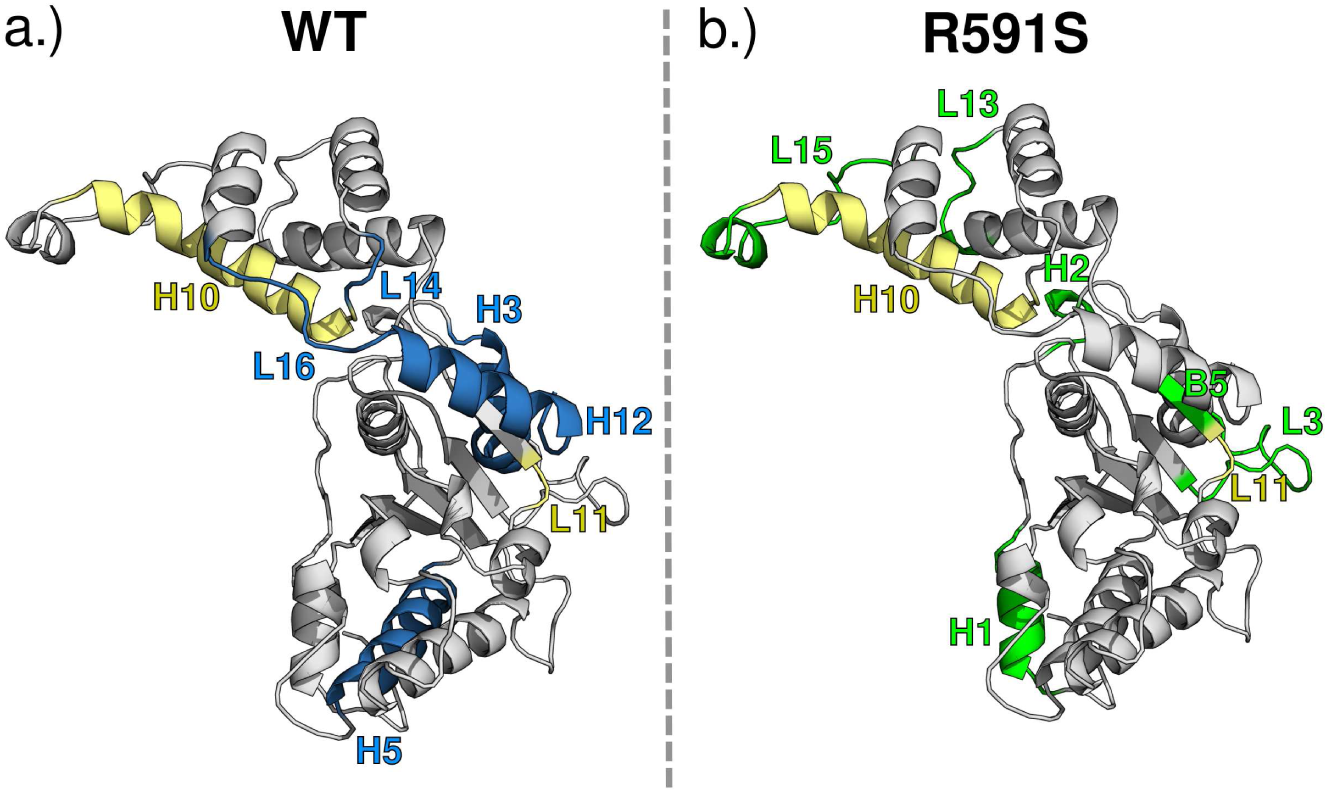
Allosteric regions that were identified as important in classifying the COMPLEX to APO transition in at least two descriptors (a) for the WT system (b) for the R591S mutant system. Regions colored in yellow were found to be important in both WT and the mutant systems. Regions colored in blue and green were found to be important only in the WT and R591S, respectively. A detailed description of the descriptors in which these regions were found to be important is in Table S20.

The RSA descriptor shows that, in the WT spastin, the removal of both ligands makes L3 and L11 from the convex interface important across multiple protomers (See Reference^16^ Figure S41(c)). This finding holds for the mutant, which, coupled with the above results for CONTACTS, indicates that L3 and L11 are important for the allosteric signaling in the mutant both through their inter-protomer contacts and the interactions with the solvent. Furthermore, additional regions become important in the mutant: L13 from the HBD, B5 from the NBD, and H1 from the concave interface in the NBD, which was important for inter-protomer contacts as well (Figure 6). Both energetic descriptors indicate that, in the WT system, L16 and H12 from the HBD become important due to ligand removal (See Reference^16^ Figure S39(c) and S40(c)). As shown in Figure S14 in the mutant system signaling in the HBD shifts to H10, while L16 and H12 are no longer important. Additionally, regions in the NBD, namely H1, B2, L7, B5, and L15, become more prevalent in the mutant system due to the removal of the ligands.

Results for WT spastin suggested that, due to the removal of both ligands, contacts between the BC, CD, and EF interfaces decreased, suggesting that the spiral hexamer of WT spastin tends to separate into dimers AB and DE, while C and F remain as monomers. However, due to the mutation, we found that the AB contacts are particularly enhanced as both H10 in A and L3 in B are increased in the CONTACTS descriptor. This result is further supported by the RSA, which shows that the regions between A and B protomers, which are H1 and H10, decrease in RSA indicating they are more buried, and by the energetic descriptors, which indicate that these interfaces are more stable (see Figure S14).

Overall, this analysis reveals that during the COMPLEX to APO transition L16, from the HBD, which was the most prevalent feature across multiple descriptors in the WT system, shifts to H1 in the NBD region as the most prevalent feature for the mutant system. Additionally, as shown in Figure 6, H12, the CT-Hlx which is crucial for oligomerization, loses its significance due to the mutation, while other regions at the convex interface (B5 and L3) become allosterically relevant during the COMPLEX to APO transition in the mutant. We also observe that L15, which includes the HBD tip, becomes significant in the mutant. This result, combined with the above finding that L15 responds allosterically to the mutation only in the APO state, strongly suggests that the HBD tip is a major allosteric element of the mutant in the absence of the ligands. These findings suggest that during the COMPLEX to APO transition, both regions important for NBD-to-NBD connections, and for HBD-to-NBD connections, are allosterically affected by the mutation.

### 3.6 Mapping Allostery with Graph Network Theory

We mapped the allosteric networks for each system by utilizing dynamic networks derived from pairwise correlations in the DCCM to analyze the impact of the mutation on the tertiary and quaternary assembly of spastin.^16^ As described in the Methods section, we used the betweenness centrality measure to quantify how essential a node is to the overall network. We identified the top 10% of central positions for comparison between the systems to detect allosteric shifts.^53,55–57^ We then paired this analysis of the dynamic network with a detailed path analysis to track how allosteric signal propagation is impacted by the mutation. We specifically looked for positions that appeared in the mutation analysis, which are used to reroute the transmission of an applied perturbation between regions of interest compared to what we found in the WT analysis.

#### Establishing the Allosteric Behavior of the Tertiary Ensemble

We first characterize and compare the allostery of the WT and mutant monomers by examining the influence of the ATP and CTT binding on the allosteric networks via the top 10% *C_B_*.^16^ For the WT, the analysis of the most central positions for each state (Table S22) reveals that both the WA and WB motifs contain highly central residues, while the PLs are never highly central. The overall populations of these most central positions are found in Figure S15, which clearly demonstrate allosteric shifts due to the presence of the ligands. Consistent with our previous findings,^16^ the APO (d) positions are spread across the structure, including on the convex side of the monomer. In contrast, the COMPLEX (a) positions populate primarily the concave side, as well as the hinge region between the NBD and HBD (demonstrated in Figure 7a). By including the NUCLEOTIDE (b) and SUBSTRATE (c) states, we see that the presence of the ATP is the primary driver of the allosteric shift towards the concave side of the monomer. Interestingly, residues from the CT-Hlx are found to “activate” in the presence of the ligands individually (either the ATP or the CTT) but not when they are both bound in the COMPLEX state. In addition, the ATP and CTT both have an “activating” effect on the end of the long helix towards the previously identified HBD tip that is not observed in either the COMPLEX or the APO states.^17^ Because both the CT-Hlx and the HBD tip are involved in the formation of interfaces between protomers in the hexameric structure of spastin, the above findings strongly suggest that the oligomerization of spastin is driven by the formation of contacts at the concave interface of a protomer upon ATP binding and then also at the convex interface upon CTT binding.

**Figure 7:**
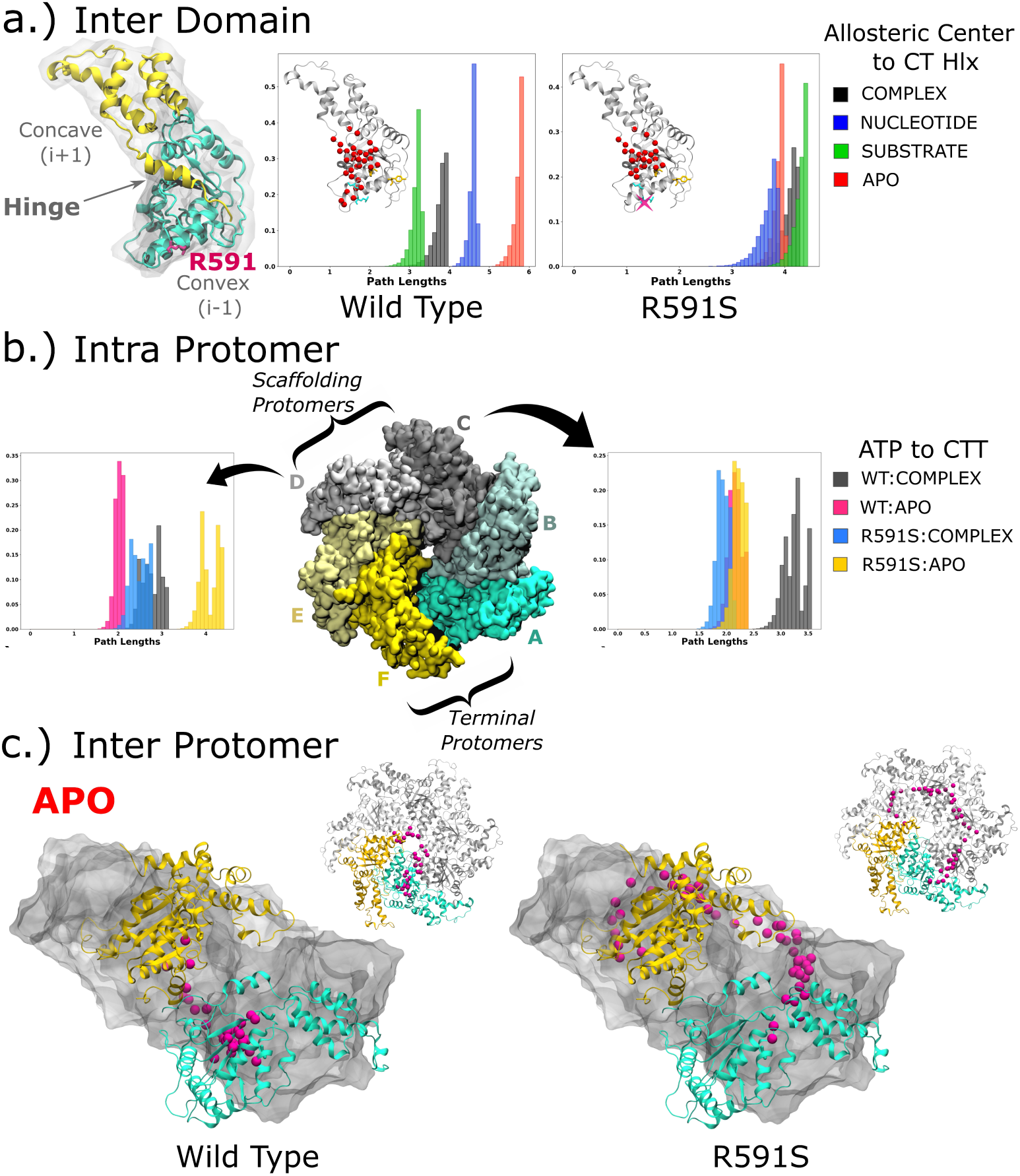
(a.) A monomer is provided to demonstrate the difference between the domains -NBD (teal) and HBD (gold) -as well as regions of interest such as the interfaces, the referenced hinge,^17^ and position R591 that is mutated to S. The distributions of the path lengths taken between domains from the Allosteric Center to the CT-Hlx are provided for the WT and the R591S systems with the highly degenerate positions for the APO system indicated by red beads. (b) The hexamer is shown colored by labeled protomer including indicators for the referenced terminal and scaffolding protomers^16^ The distributions of the intra-protomer path lengths taken for the scaffolding protomers from the ATP binding region to the CTT binding region for each hexameric system. (c.) The highly degenerate positions from the inter-protomer paths taken from the ATP binding region of protomer A to the CTT binding region of protomer F for the APO state.

When the R591S mutation is applied (Table S23), the WA motif remains always central, but the WB is no longer central in the NUCLEOTIDE state. We also found that PL1 becomes central in each state (K555 or E561), except for the COMPLEX, and that PL2 becomes central in the SUBSTRATE state (A598 & R601). These are important changes as they indicate that this single point mutation is impacting the WB motif, which is responsible for the ATP hydrolysis, by abolishing its centrality, and also the PLs, which are responsible for the CTT binding, by activating their centrality, primarily in the SUBSTRATE state. This indicates that the mutation largely disrupts how the ATP interacts with the internal network of the tertiary system, and to a lesser extent the interaction of the substrate with the spastin monomer. We also find that the CT-Hlx is only active in the NUCLEOTIDE state in the R591S system, thus losing the activity in the SUBSTRATE state seen in the WT case. In addition, the *C_B_* signal near the HBD tip is now more active in the COMPLEX and APO states, rather than in the NUCLEOTIDE and SUBSTRATE states seen in the WT, which may indicate a specific disruption of the allosteric networks responsible for the communication between the NBD and the HBD. To directly explore these relationships, we carried out a detailed path analysis from the ATP binding pocket (P525/T530) to the CTT binding region (H596/R601) and from the allosteric center (S589/R591S) to the CT-Hlx (W749/Y753).^16^ The optimal (Dijkstra’s) paths with their path lengths are shown in Tables S24 and S25. This allows us to probe the impact of the mutation on the communication within the NBD, and between the NBD and HBD.

First we extracted 80,000 paths from the ATP binding pocket to the substrate binding PLs to explore the impact of the mutation on ligand binding and to identify how the key co-factor binding sites communicate with each other. We identify those nodes that were found to be highly degenerate in the collection of paths (Figure S17). The mutation influences the paths taken in each state, except for the COMPLEX state. This is further demonstrated in the comparison of the suboptimal path lengths in Figures S16a and S16c.

We summarized the regions important for allosteric signal propagation by calculating the average degeneracy for each alpha helix, beta strand, and loop section, as done in our previous study,^16^ and found that the source and the sink are always degenerate. The allosteric signal in the NUCLEOTIDE state of the WT propagates through regions from the bottom surface (H1/L1) of the motor, whereas in the mutant the same regions are highly degenerate in the APO state. We note that the WB (L8) portion of the ATP binding pocket is degenerate only in the SUBSTRATE state for the R591S mutant, which is in contrast to the WT where it is degenerate in the COMPLEX and APO states. The distribution of the suboptimal path lengths highlights that the presence of the mutation causes a striking shift towards shorter paths in the absence of the binding ligands. This is a reflection of the change in correlations observed in the APO state between the WT and R591S systems where the calculated cutoff changed from 0.5 to 0.6 (Table S21). In addition, we found that the mutation leads to high degeneracy in B1, L5, L7, and B5, which are not found in the WT. This analysis indicates a distinct reorganization of the allosteric signal propagation in the core of the NBD as a result of the R591S mutation.

We then compared the paths taken from the allosteric center to the CT-Hlx from our previous work,^16^ with the ones found in the mutant to describe the inter-domain communication facilitated by the allosteric center. We map the highly degenerate residues in Figures S18 and 7a and provide the distribution of path lengths in Figures S16b and S16d. Similar to the previously described collection of paths, we find that both the source (L8) and the sink (H12) are degenerate. Unlike with the previous set of paths inside the NBD region, we found that the mutation largely impacted the communication between the NBD and HBD both in path length and in rigidity. In the COMPLEX state, the mutation causes a shift in the rigidity of the network from the perfect degeneracy found in the PL3 region (B4 and L10) in the WT to the degeneracy of the last strand in the beta-sheet of the NBD core. This indicates that the binding of the ligands to the monomer, which counteracted the effect of the mutation on the allosteric communications inside the NBD, is unable to restore the communication between the NBD and the HBD. Furthermore, the suboptimal path lengths indicate dramatic shifts in the four states: the NUCLEOTIDE, and the APO path lengths become shorter, while the SUBSTRATE and the COMPLEX paths become longer in the mutant compared to the WT. The changes in path lengths suggest that the HBD reacts differently to the presence of the ligands in the mutant compared to the WT and underlines the important role played by the allosteric center in the communication between the domains. This is particularly important in the COMPLEX state where the mutation leads to a weaker allosteric coupling between the NBD and the HBD regions of the spastin monomer.

Regarding the regions showing high degeneracy in the NBD to HBD paths, we found that, in the WT, the WA motif is degenerate in the NUCLEOTIDE and APO states, while in the mutant it is degenerate in the NUCLEOTIDE and SUBSTRATE states. The PL regions are found to be degenerate in each state of the WT, except for PL1 in the APO state. This trend is carried into the mutant, with the exception of PL2 in the APO state such that neither of the PLs are degenerate in the absence of the ligands (see Figure 7a). We infer that the allosteric signal between the NBD and the HBD in the mutant does not travel through the CTT binding region in the absence of the ligands. In the HBD, the allosteric signal propagates through H10 with perfect degeneracy in the NUCLEOTIDE state for the WT, while for the mutant this occurs in the SUBSTRATE state. The previously identified loop preceding the CT-Hlx (L16) was perfectly degenerate in the SUBSTRATE state of both the WT and the mutant. This indicates that the mutation causes the inter domain communication to become more flexible in the presence of the ATP alone and far more rigid (larger number of regions with high degeneracy) in the presence of the substrate alone.

In summary, we find that the allosteric network of the NBD core of the tertiary network exhibits a shift in both the path and flexibility, especially in the binding of individual co-factors. When binding both cofactors (COMPLEX state) we observe a restoration of the WT behavior, which is significant as we know that severing still occurs in the HSP mutant. In contrast, the propagation of allosteric signals between the NBD and the HBD was substantially disrupted in the mutant, with the allosteric coupling becoming weaker in the COMPLEX state. This result aligns with the above finding from the PCA that the mutation influences the hinge motions between the domains in the spastin monomer.^17^

#### The Allosteric Influence of the Neighboring Protomers on the Intra-Protomer Paths

Following our previous study of WT spastin, we then compared the results of the intra-protomer network analysis in the hexamer to the results from the monomer to identify uniquely quaternary signals. From the Top 10% *C_B_* results in the COMPLEX state, we found that more positions in protomers B and F became central in the mutant (see Table S26), while positions in protomers A, C and E became less central. Most of the observable differences between the mutant and the WT (see Figure S19a) are found at the central pore where more positions in the PL1 in each protomer, but especially in protomers E and F, become more central in the mutant. In the WT, the ARG fingers were highly central in all the protomers with the exception of A, while in the mutant we only find them central in B and F. In addition, the CT-Hlx in protomers E and F are no longer central in the mutant. Because these regions are important for oligomerization, the loss of high centrality in the mutant suggests a disruption to the inter-protomer communication when both ligands are bound to the hexamer. In contrast, in the absence of the ligands (APO; see Table S27), protomers A and D become more central, while protomers B and F become less central. The observed differences are notably in the central pore and in the terminal protomers (see Figure S19b). In the WT, the regions found to be highly central varied from protomer to protomer, but in general corresponded to the ATP binding pocket, PL1, PL3 and the CT-Hlx. We also found that PL2 was central in protomers E and F. In the mutant, we do not find PL3 to be highly central, nor PL2 in any protomer. This points to a disruption in the PL coordination due to the R591S mutation.

We then extracted similar collections of paths as the ones extracted from the tertiary systems to determine how the presence of neighboring protomers influences the communication between the pairs of source and sink sites. The optimal paths for the WT and R591S mutant in the COMPLEX and APO states are provided in Tables S29-S35. For the communication inside the NBD (ATP to CTT), we found that the influence of the neighboring protomers in protomers A through C in the COMPLEX and APO states is similar between the WT and the mutant in that the allosteric signal propagates from the concave interface to the convex interface and from there down to the sink. This often includes propagation through the region around PL3 (H7). The suboptimal path lengths distributions (Figure S20a-c) illustrate differences due to the mutation in the efficiency of passing allosteric signals. While protomer A had only minimal changes to its suboptimal distributions within the NBD, we found that, compared to the WT, the mutant paths are very short for the COMPLEX, particularly in protomer C (Figure 7b). The suboptimal path lengths also indicate that the R591S APO state (shown in gold) is characterized by relatively short paths in every protomer but D (Figure 7b). This suggests the mutation may uniquely affect the scaffolding role of the internal protomers (C and D) in the APO assembly. In protomer F, the paths tend to propagate down the concave interface in every instance, with similar suboptimal path distributions even though the R591S APO paths favor longer lengths (Figure S20f). The tertiary networks indicated that the cooperative presence of the ligands restored the impact of the mutation. In our previous study we found the presence of the neighbor was very influential on the quaternary signal propagation.^16^ Here, we report that the presence of the neighbor does show restorative influence on the allosteric network of the NBD, accounting for the similarity in pathways. We find the most notable changes in the NBD communication due to the presence of the mutation in the internal scaffolding protomers (C and D).

Interestingly, when analyzing the inter domain paths in the quaternary assembly, we found little change in the pathways taken and in the suboptimal distributions, regardless of the protomer (Figure S21). This is dramatically different from the suboptimal paths in the monomers where the NBD-HBD communication was significantly altered in the presence of the mutation. We find this to be consistent with the previously reported finding that the neighboring protomer has a significant impact on the communication between domains.^16^ We do note that in protomer F the R591S APO state favors slightly longer paths, which is reminiscent of the behavior observed in the NBD to NBD allosteric signal propagation paths.

#### The Allosteric Changes to the Scaffolding and Active Terminal Protomers (inter-protomer paths)

Due to the presence of the allosteric center at the i to i-1 interface,^12^ we took the same collection of paths (NBD-NBD and NBD-HBD) between the indicated neighboring protomers to probe the effects of the mutation on each interface. The results reflect the asymmetry previously described of the hexamer where each interface behaves differently from the rest.^16^ This is somewhat captured by slight differences in the regions found to be degenerate, but more easily identified via the changes in the range of the suboptimal paths shown in Figures S22 and S25. Notably, the B to A in S22b and S25b and the F to E suboptimal paths for both inter-protomer path collections in S22f and S25f show that, in the APO state of the R591S system, they are pushed towards longer paths than in the COMPLEX state -a trend reflected in the intra paths described in the previous section. This suggests a weakening of the allosteric communication between protomers in the APO state due to the mutation. The suboptimal path lengths from D to C (Figure S22d and S25d) and C to B (Figure S22c and S25c) were the longest in the assembly. The analysis of the intra protomer paths suggested the mutation disrupted the scaffolding of the motor at protomer D. The inter-protomer paths suggest that the main perturbation of the scaffolding is between protomers D and C, particularly from the NBD of D to the HBD of C. In addition, the distributions when the ligands are bound (APO to COMPLEX) shift from shorter to longer paths in the WT, while in the mutant they go from longer to shorter paths. In general, we identified that the R591S mutant network favors PL1 when communicating between the i and i-1 instead of a larger number of regions used by the WT such as PL2, PL3 and the ATP binding regions. Thus the mutant networks are unable to access these other regions required for network flexibility in the WT thereby suggesting that they are more rigid. Protomer F is the i-1 neighbor for protomer A. In the open spiral conformation presented in this study, A and F are not in contact with each other, which is why they are referred to as the “terminal protomers” - indicated in Figure 7b. As referenced throughout the text, the results from our previous study led us to propose that protomers A and F are the catalytically active protomers in the asymmetric hexameric motor.^16^ We also identified that the allosteric signal from A to F would propagate around the pore primarily via PL1 and occasionally through PL2 (particularly in protomer E). This trend was mostly conserved in the COMPLEX state of the R591S mutant, except in protomer A where it passes via PL3 and the HBD (see Figures S23 and S26). Remarkably, when taking these paths from A to F in the APO state, the mutant cannot engage with the PLs as shown in Figure 7c. Instead, the signal propagates from protomer to protomer via the CT-Hlx and the loop preceding it (L16) with perfect degeneracy (see Figures S24 and S27). The distributions of the suboptimal path lengths indicate this detour causes longer paths in both collections (Figures S22a and S25a). This speaks to a loss of coordinated signaling at the pore of the spiral hexamer in the mutation, primarily in the absence of the binding partners, while in the presence of the ligands the effect of the mutation is more modest.

## 4. CONCLUSIONS

The identities of most of the HSP-related mutations are known. In contrast, an understanding of the molecular factors that lead to the functional changes is currently missing. In our study we focused on a very important HSP-related mutation, which has been proposed to affect spastin’s oligomerization, and it is located at the site of the previously identified allosteric center of spastin: R591S.^12,16^ We found that the mutation reduces dramatically the binding affinity of the spastin monomer for the substrate, in the presence of ATP, resulting in the detachment of the substrate first from PL2, and then from PL1. The order of unbinding of the CTT from the pore loops agrees with the experimental finding that the absence of the substrate results in a large increase in the flexibility of PL2 leading this loop to become intrinsically disordered.^8^ This behavior is in stark contrast with the lack of effect of the mutation on the binding of the ATP to the monomer. This is true even at the level of the hexamer, as the mutation does not induce any changes in the nucleotide binding regions of the various protomers. Intriguingly, our simulations also show that, in the absence of the ligands, the mutation induces rigidity in both the spastin monomer and hexamer, while in the WT spastin the absence of ligands leads to an increase in flexibility. These findings suggest that the binding of the substrate to the spastin monomer is likely a conformational selection process, which is impaired when the monomer is unable to explore a large set of structures, as it happens in the R591S mutant. In contrast, the binding of the ATP to the monomer must be an induced fit process where only localized changes near the binding site are required for high affinity binding.

Our simulations also show that, in the hexameric state, the loss of the ligands results in changes in the orientation of the terminal protomers with respect to the rest of the structure. While this behavior is common between the WT and the mutant, the changes are more pronounced in the mutant. In contrast, the internal structure of the terminal protomers changes less upon the loss of ligands in the mutant than in the WT, as discussed above. Focusing on the functional regions of the hexamer, we found that, in the WT, the CTT binding regions (PL1 and PL2) of protomer F re-orient dramatically with respect to the rest of the hexamer upon the loss of ligands, while in protomers A and B the CTT binding regions stay mostly in place. This behavior is entirely reversed in the mutant, where the CTT binding regions of protomers A and B change substantially with respect to the rest of the hexamer, while in protomer F they stay in place. These differences result in stark changes in the essential dynamics of the mutant versus the WT, with functional implications. Namely, in the WT hexamer, when both ligands are bound, the major motion corresponds to protomers A and F moving axially away from each other resulting in the opening of the gate and the up and down motion corresponding to the pulling and wedging severing action.^2,3,5^ In the mutant this motion is no longer present, being replaced by twisting motions of the A and B protomers, with the rest of the protomers being almost frozen in place. Therefore, the functional hexamer, upon the HSP-related mutation, loses the ability to execute the primary motion associated with the MT severing action. In the absence of the ligands, the WT hexamer populates states corresponding to horizontal motions of protomers A and F moving away from each other. No such states are accessible for the mutant, which instead populates two states where the gate between the terminal protomers A and F is mostly closed. This likely points towards a different hexamer dissociation (and potentially association) pathway in the mutant compared to the WT.

We found that the increased stability (rigidity) in the protomers due to the mutation discussed above can at least in part be attributed to the fact that, when both ligands are bound to the hexamer, in the mutant there are additional salt bridges formed between protomers and intra-protomers in the HBD region. The mutation also leads to increased lifetimes of the intra-protomer salt bridges between positions in both the NBD and the HBD regions that are in common with the WT.

Our analysis of the allosteric response in the spastin hexamer as a result of the mutation shows that the effect is greater (more descriptors show a signal for a given secondary structure element) when both ligands are bound. The regions that are most perturbed are those involved in the formation of the NBD to NBD and the NBD to HBD inter-protomer contacts: secondary structures (loops and helices) from the convex side of the NBD, and, respectively, from the concave side of the HBD. We found a similar result when studying the changes due to the mutation in the allosteric transition from the fully ligand bound hexameric spastin to the APO state. These findings lend support to the proposed role of the R591S mutation in affecting the oligomerization of spastin.^12^

Our graph network analysis of the high centrality regions of the hexamer revealed a disruption in the pore loops PL2 and PL3 coordination due to the R591S mutation: these loops are highly central in the WT, but lose their centrality in the mutant. A similar loss is found in the CT-Hlx from the HBD region of a protomer. Because PL3 and CT-Hlx are important for oligomerization, the loss of high centrality in the mutant suggests a disruption to the inter-protomer communication when both ligands are bound to the hexamer. Further support comes from the finding that the inter-protomer allosteric paths in the mutant are unable to access many of the regions required for network flexibility in the WT. This indicates that the presence of the mutant induces rigidity in the allosteric networks of the ligand-bound state making it more likely to experience loss of function as any applied perturbations (other ligands, further mutations, changes in solvent) would not be easily dissipated by passing through a variety of alternative paths, as required in other proteins.^22,61^ In contrast, the allosteric communication between the terminal protomers, A and F, in the absence of ligands, exhibits a dramatic change due to the mutation indicative of a weakening of the allosteric communication between end protomers, with likely effects on the dissociation of the hexamer. Recent work combining experimental deep scanning mutagenesis and computational modeling of the allostery of TetR proposed that the design of strong allosteric inhibitors should take into account the changes to the free energy landscape (FEL) between the WT and potential mutants.^61^ The idea is that a strong inhibitor will result in dramatic changes of the FEL. Our findings regarding the changes in the allostery of spastin upon the R591S mutation align with this proposal, as this mutation is known to have deleterious effects on the severing function of spastin and it leads to substantial changes in the FEL. In turn, keeping in mind the disease-related roles of the HSP mutations that call for the development of therapeutics, our results suggest that, in addition to the recovery of the WT FEL, additional stringent criteria have to be used for design challenges geared towards the recovery of the proper function of this MT severing machine. Namely, any engineered additional mutations or binding factors should restore (i) the high centrality of the pore loops that are engaging the substrate; (ii) the structural flexibility of the terminal protomers, and the cooperative motions of the protomers in the hexamer, and (iii) the up and down motions of the terminal protomers responsible for the pulling and wedging severing action. The application of our approach to future studies of the effect of other HSP-related mutations on spastin’s dynamics will enable us to discover overarching principles underlying the impaired functionality of this molecular motor in disease.

## Acknowledgement

This research was funded by the National Science Foundation (NSF) MCB-1817948 (to RID). C. StC. was supported through the NSF Research Experience for Undergraduates in Chemistry grant CHE-1950244. This work used the advanced computing systems available through ACCESS based on the allocation BIO210094 (to RID).

## Supporting Information Available

Additional tables and figures providing detailed information for the results discussed in the main text such as RMSD values, list of high centrality positions, list of shortest allosteric paths, representations of PC motions, free energy landscapes with representative structures, and distributions of suboptimal paths.

