## Supplemental tables and figures for "A major disease-related point mutation in spastin alters dramatically the dynamics and allostery of the motor"

**Equation S1.** The equation used to calculate the time dependent correlations between the residue pairs i and j. Here, $r_i(t)$ and $r_j(t)$ are the position vectors of $C_\alpha$ atoms of residues i and j and <·> represents the time ensemble average[5] $\Delta r_i(t) = r_i(t) - < r_i(t) >_t$ and $\Delta r_j(t) = r_j(t) - < r_j(t) >_t$

$$DCCM_{(i,j)} = \frac{<\Delta r_i(t) \bullet \Delta r_j(t)>_t}{\sqrt{(<||\Delta r_i(t)||^2>_t <||\Delta r_j(t)||^2>_t)}}$$

**Equation S2.** The equation used to determine the DCCM convergence where N is the total number of residues and τ = 5ns[5]

$$R(t) = \frac{1}{N} \sum_{(i,j)} (DCCM_{(i,j)}(t) - DCCM_{(i,j)}(t - \tau))^2$$

**Equation S3.** The root-mean-square inner product (RMSIP) used to measure the overlap between the extracted subspaces from the essential dynamics analysis (PCA). Here, the eigenvectors of the two subspaces, A and B are $\eta_i^A$ and $\nu_i^B$. The top 10 eigenvalues in the WT were found to cover around 90% of the overall variance so we set J to 10[5].

$$RMSIP = \left( \frac{1}{J} \sum_{i,j=1}^{J} \left( \eta_i^A \bullet \nu_i^B \right) \right)^{1/2}$$

**Equation S4.** The betweenness centrality equation determines the number of shortest paths between other nodes that pass through the node of interest $(v)$[1,2].

$$c_B(v) = \sum_{(s,t \in V)} \frac{\sigma(s,t|v)}{\sigma(s,t)}$$

**Table S1.** Summary of the MD setups for the R591S mutated spastin hexamer

| State | Nucleotide | Substrate | #Atoms | #Residues | #Water | #Na | #Trajectories | Total Simulation time (ns) |
|---|---|---|---|---|---|---|---|---|
| **COMPLEX** | ATP | E15 | 245340 | 1839 | 75672 | 63 | 5 (300ns) | 1500 |
| **APO** | - | - | 245508 | 1824 | 75878 | 30 | 5 (200ns) | 1000 |

**Table S2.** Summary of the MD setups for the WT spastin monomer. We extended simulations from our previous study to better characterize tertiary allostery[9].

| State | Nucleotide | Substrate | #Atoms | #Residues | #Water | #Na | #Trajectories | Total Simulation Time (ns) |
|---|---|---|---|---|---|---|---|---|
| **COMPLEX** | ATP | E15 | 84034 | 319 | 26944 | 22 | 3 (200ns) | 600 |
| **NUC** | ATP | - | 74836 | 304 | 23934 | 7 | 3 (200ns) | 600 |
| **SUB** | - | E15 | 84029 | 319 | 26958 | 19 | 3 (200ns) | 600 |
| **APO** | - | - | 74843 | 304 | 23952 | 4 | 3 (200ns) | 600 |

**Table S3.** Summary of the MD setups for the R591S mutated spastin monomer

| State | Nucleotide | Substrate | #Atoms | #Residues | #Water | #Na | #Trajectories | Total Simulation Time (ns) |
|---|---|---|---|---|---|---|---|---|
| **COMPLEX** | ATP | E15 | 74885 | 319 | 23897 | 23 | 3 (200ns) | 600 |
| **NUC** | ATP | - | 66512 | 304 | 21162 | 8 | 3 (200ns) | 600 |
| **SUB** | - | E15 | 74829 | 319 | 23894 | 20 | 3 (200ns) | 600 |
| **APO** | - | - | 66552 | 304 | 21191 | 5 | 3 (200ns) | 600 |

**Table S4.** Global Average RMSD values for the WT and the R591S mutant spastin monomer states

| | **WT** | **R591S Mutant** |
|---|---|---|
| | Global Average (Å) | Global Average (Å) |
| **COMPLEX** | $4.25 \pm 0.57$ | $4.03 \pm 0.45$ |
| **NUCLEOTIDE** | $4.14 \pm 0.38$ | $3.99 \pm 0.39$ |
| **SUBSTRATE** | $4.44 \pm 0.66$ | $4.20 \pm 0.52$ |
| **APO** | $5.77 \pm 0.65$ | $3.89 \pm 0.34$ |

**Table S5.** Global Average RMSD values for the WT and R591S mutant spastin hexamer states

| | **WT** | **R591S Mutant** |
|---|---|---|
| | Global Average (Å) | Global Average (Å) |
| **COMPLEX** | $5.776 \pm 0.56$ | $5.98 \pm 0.84$ |
| **APO** | $6.474 \pm 1.00$ | $4.85 \pm 0.53$ |

**Table S6.** Percentage of central structures from the first clustering step covered by the top two clusters in the second clustering step for each state of the WT and the mutant systems across different RMSD cutoff values. This was used to determine the optimal cutoff for the second clustering step in the RMSD-based double clustering analysis. A cutoff of 0.45 (highlighted in bold) was selected, as it covers nearly 70% of the structures in all setups.

| | **WT** | | **R591S Mutant** | |
|---|---|---|---|---|
| RMSD-Cutoff | **COMPLEX** | **APO** | **COMPLEX** | **APO** |
| 0.15 | 1.34% | 0.98% | 0.83% | 0.96% |
| 0.2 | 10.80% | 7.13% | 6.43% | 7.10% |
| 0.25 | 28.24% | 18.79% | 17.35% | 19.76% |
| 0.3 | 45.44% | 33.84% | 31.93% | 28.71% |
| 0.35 | 55.57% | 48.42% | 41.08% | 36.93% |
| 0.4 | 70.19% | 55.87% | 49.61% | 51.21% |
| **0.45** | ***90.89%*** | ***71.42%*** | ***65.13%*** | ***77.82%*** |
| 0.5 | 96.50% | 84.37% | 87.08% | 96.45% |

**Table S7.** The residue numbers associated with the functional binding motifs in spastin's spiral conformation[7,8].

| **Functional Region** | | **Res. No** |
|---|---|---|
| **ATP - Binding** | Walker A (WA) | 523 - 530 |
| | Walker B (WB) | 581 - 585 |
| | Arginine Fingers | 640 - 641 |
| **CTT - Binding** | Pore Loop 1 (PL1) | 552 - 562 |
| | Pore Loop 2 (PL2) | 594 - 601 |

| | | |
|---|---|---|
| **Oligomerization** | Pore Loop 3 (PL3) | 629 - 637 |
| | CT-Helix | 739 - 756 |

**Table S8.** The variance covered (%) in the first two PCs for the **global motions** of the WT system and the R591S mutant system. The corresponding PC motions are shown in Figure 2 and S6.

| Global | WT | R591S Mutant |
|---|---|---|
| *COMPLEX* | | |
| **PC 1** | 49.3 | 43.1 |
| **PC 2** | 26.5 | 16.9 |
| **Variance in top 2 PCs** | **75.8** | **60.0** |
| **Variance in top 10 PCs** | **89.2** | **87.8** |
| *APO* | | |
| **PC 1** | 67.5 | 33.8 |
| **PC 2** | 16.2 | 13.9 |
| **Variance in top 2 PCs** | **83.7** | **47.7** |
| **Variance in top 10 PCs** | **94.0** | **85.1** |

**Table S9.** The variance covered (%) in the first two PCs for the **PL1 motions** of the WT system and the R591S mutant system. The corresponding PC motions are shown in Figures S10 and S11.

| PL1 | WT | R591S Mutant |
|---|---|---|
| *COMPLEX* | | |
| **PC 1** | 49.6 | 31.8 |
| **PC 2** | 17.7 | 21.2 |
| **Variance in top 2 PCs** | **67.4** | **53.0** |
| *APO* | | |
| **PC 1** | 52.6 | 27.6 |

| | | |
|---|---|---|
| **PC 2** | 20.2 | 24.4 |
| **Variance in top 2 PCs** | **72.8** | **52.0** |

**Table S10.** The variance covered (%) in the first two PCs for the **PL2 motions** of the WT system and the R591S mutant system. The corresponding PC motions are shown in Figures S10 and S11.

| PL2 | WT | R591S Mutant |
|---|---|---|
| *COMPLEX* | | |
| **PC 1** | 42.1 | 33.8 |
| **PC 2** | 20.2 | 20.3 |
| **Variance in top 2 PCs** | **62.3** | **54.1** |
| *APO* | | |
| **PC 1** | 59.1 | 47.0 |
| **PC 2** | 17.8 | 15.6 |
| **Variance in top 2 PCs** | **76.9** | **62.6** |

**Table S11.** The **RMSIP values** between the top 10 PCs of the WT and R591S. This indicates moderate overlap between the PC spaces in spite of the observed difference in the PC1/PC2 variance (particularly the global motions) caused by the point mutation for both the COMPLEX and APO states.

| WT-R591S | COMPLEX | APO |
|---|---|---|
| **Global** | 0.634 | 0.644 |
| **PL1** | 0.700 | 0.682 |
| **PL2** | 0.635 | 0.689 |

**Table S12.** The backbone RMSD (Å) values, calculated using VMD, for the representative structure of each FEL minima of the WT and mutant systems, relative to their starting structures in the **COMPLEX state**, shown in Figure S7.

| COMPLEX | WT | R591S |
|---|---|---|
| Minima i | 6.5 | 9.6 |

| | | |
|---|---|---|
| Minima ii | 6.4 | 6.9 |
| Minima iii | 4.5 | 6.6 |
| Minima iv | - | 3.9 |
| Minima v | - | 4.2 |

**Table S13.** The backbone RMSD (Å) values, calculated using VMD, for the representative structure of each FEL minima of the WT and mutant systems, relative to their starting structures in the **APO state**, shown in Figure S8.

| APO | WT | R591S |
|---|---|---|
| Minima i | 10.4 | 7.3 |
| Minima ii | 5.9 | 4.7 |
| Minima iii | 4.7 | 5.2 |
| Minima iv | - | 4.7 |
| Minima v | - | 3.4 |

**Table S14.** The backbone RMSD (Å) values, calculated with VMD, of the representative structure for each FEL minima for the **R591S in the WT space** to the starting structure of the WT hexamer in the COMPLEX and APO states shown in Figure S9.

| R591S in WT | COMPLEX | APO |
|---|---|---|
| Minima i | 7.4 | 5.5 |
| Minima ii | 5.6 | 5.0 |
| Minima iii | 9.3 | - |
| Minima iv | 4.0 | - |
| Minima v | 8.1 | - |

**Table S15.** The list of **inter-protomer salt bridges** within the **pore loop residues (PL1 and PL2)** of the WT and R591S mutated spastin, observed in at least 3 protomers for more than **10 ns** during the MD simulations. The protomers indicated in parentheses denote where the salt bridge is present, and the mentioned time represents the average persistence time of each salt bridge with its standard deviation.

| | WT | | R591S | |
|---|---|---|---|---|
| **Setup** | **SB** | **Time (ns)** | **SB** | **Time (ns)** |
| **COMPLEX** | D559-K555 [BA,CB,ED,FE] | 41.6±21.27 | D559-K555 [BA,CB,DC,ED,FE] | 71.97±21.3 |
| **APO** | D559-K555 [BA,CB,DC,ED,FE] | 37.62±7.8 | D559-K555 [BA,CB,DC,ED,FE] | 52.01±18.3 |

**Table S16.** The list of **inter-protomer salt bridges** within **all other residues (except PL1 and PL2)** of the WT and R591S mutated spastin, observed in at least 3 protomers for more than **10 ns** during the MD simulations.

| | WT | | R591S | |
|---|---|---|---|---|
| **Setup** | **SB** | **Time (ns)** | **SB** | **Time (ns)** |
| **COMPLEX** | D471-K459 [BC,CD,DE] | 32.5±24.5 | D471-K459 [AB,BC,CD,DE,EF] | 29.53±17.1 |
| | E472-K459 [BC,CD,DE,EF] | 21.57±9.34 | E472-K459 [BC,CD,DE,EF] | 45.59±13.2 |
| | E497-R704 [BA,CB,DC,ED,FE] | 105.37±19.9 | E497-R704 [BA,CB,DC,ED,FE] | 153.86±19.5 |
| | D585-R591 [AB,BC,CD] | 11.66±0.45 | | |
| | | | E597-K603 [BA,CB,FE] | 29.36±15.9 |
| | | | D618-K479 [BA,DC,ED] | 22.49±12.6 |
| | D697-K644 [AB,BC,CD,DE,EF] | 37.04±23.0 | D697-K644 [AB,BC,CD,DE,EF] | 41.74±7.9 |
| **APO** | D471-K459 [AB,BC,CD,DE,EF] | 28.66±9.5 | D471-K459 [AB,BC,CD,DE,EF] | 29.97±10.6 |
| | E472-K459 [BC,CD,DE] | 21.3±6.6 | | |

| | | | | |
|---|---|---|---|---|
| | E497-R704 [BA,CB,DC,ED,FE] | 68.09±41.4 | E497-R704 [BA,CB,DC,ED,FE] | 77.42±27.5 |
| | D618-K696 [BA,CB,DC,ED,FE] | 20.04±17.0 | D618-K696 [BA,CB,ED,FE] | 24.78±6.4 |
| | D690-R640 [AB,BC,CD,DE,EF] | 70.57±25.6 | D690-R640 [AB,BC,CD,DE,EF] | 35.34±16.3 |
| | D697-K644 [AB,BC,CD,DE,EF] | 62.51±42.5 | D697-K644 [AB,BC,CD,DE,EF] | 58.01±11.0 |

**Table S17.** The list of **intra-protomer salt bridges** within the **pore loop residues (PL1 and PL2)** of the WT and R591S mutated spastin, observed in at least 3 protomers for more than **10 ns** during the MD simulations.

| | WT | | R591S | |
|---|---|---|---|---|
| **Setup** | **SB** | **Time (ns)** | **SB** | **Time (ns)** |
| **COMPLEX** | | | D559-K562 [A,C,D,E,F] | 24.11±8.3 |
| | | | E561-R601 [A,B,C,D,E,F] | 43.42±26.7 |

**Table S18.** The list of **intra-protomer salt bridges** within **all other residues (except PL1 and PL2)** of the WT and R591S mutated spastin, observed in at least 3 protomers for more than **10 ns** during the MD simulations.

| | WT | | R591S | |
|---|---|---|---|---|
| **Setup** | **SB** | **Time (ns)** | **SB** | **Time (ns)** |
| **COMPLEX** | E462-K464 [B,C,D,E] | 15.13±4.82 | E462-K464 [B,C,D,E,F] | 14.76±2.7 |
| | | | D484-R662 [C,E,F] | 30.3±12.7 |
| | D484-K666 [A,B,C,D,E] | 80.04±24.9 | D484-K666 [A,B,C,D,E,F] | 105.77±28.9 |
| | E497-K644 [A,B,E,F] | 28.42±12.0 | E497-K644 [A,B,C,D,E,F] | 41.88±20.0 |
| | E507-R505 [A,B,C,D,E,F] | 28.94±5.9 | E507-R505 [B,C,D,E,F] | 48.34±3.9 |

| | | | | |
|---|---|---|---|---|
| | E595-K603 [A,B,C,D,E,F] | 128.23±19.5 | E595-K603 [A,B,C,D,E,F] | 123.02±41.6 |
| | E605-R565 [A,B,C,D,E,F] | 63.65±9.9 | E605-R565 [A,B,C,D,E,F] | 135.73±47.5 |
| | E609-R565 [B,C,D,E] | 45.91±12.9 | E609-R565 [B,C,D,E,F] | 54.94±26.1 |
| | E609-R572 [B,C,D,E] | 37.73±6.6 | E609-R572 [B,C,D,E,F] | 74.94±38.5 |
| | D611-K517 [B,C,E,F] | 52.55±20.2 | D611-K517 [A,B,C,D,E,F] | 73.69±22.1 |
| | D620-K517 [B,C,D,E,F] | 75.79±32.8 | D620-K517 [A,B,C,D,E,F] | 70.69±30.9 |
| | D620-R572 [B,C,D,E,F] | 30.6±22.9 | D620-R572 [C,D,E] | 54.80±6.6 |
| | | | E633-R630 [A,B,C] | 22.24±2.4 |
| | | | D635-K603 [A,B,C,D,E,F] | 53.72±27.4 |
| | E653-K681 [A,B,C,D,E,F] | 46.04±5.3 | E653-K681 [A,B,C,D,E,F] | 60.62±14.9 |
| | E657-R677 [A,B,C,D,E,F] | 18.30±3.9 | E657-R677 [A,B,C,D,E,F] | 27.39±5.2 |
| | D697-R704 [A,B,C,D,E] | 76.44±11.9 | D697-R704 [A,B,C,D,E] | 110.82±15.2 |
| | D697-R733 [A,C,E,F] | 14.58±0.9 | D697-R733 [A,B,C,D,E,F] | 19.67±4.7 |
| | | | E701-R704 [A,C,E] | 15.0±3.6 |
| | E701-R733 [A,B,C,D,E] | 22.97±3.4 | E701-R733 [A,B,C,D,E] | 38.23±11.3 |
| | E705-R720 [A,B,C,D] | 17.88±4.9 | E705-R720 [A,B,C,D,E] | 21.72±6.3 |
| | | | E709-K712 [A,B,D,E] | 11.86±0.6 |
| | E724-R678 | 16.11±2.7 | E724-R678 | 21.64±6.1 |

| | | | | |
|---|---|---|---|---|
| | [B,C,D,E] | | [A,B,C,D,E] | |
| | D726-R720 [A,C,D,E] | 21.25±9.3 | D726-R720 [A,B,C,D,E] | 25.33±8.2 |
| | | | D752-K748 [B,C,E,F] | 14.18±4.2 |
| | D755-K644 [B,C,D,E,F] | 83.82±39.1 | D755-K644 [B,C,D,E,F] | 77.97±15.7 |
| | | | D755-R645 [B,C,D,E,F] | 15.92±3.5 |
| **APO** | E462-K562 [A,B,E] | 18.10±7.9 | | |
| | D484-R662 [A,B,C,E,F] | 18.13±7.0 | D484-R662 [A,B,C,D,E,F] | 17.28±4.2 |
| | D484-K666 [A,B,C,D,E,F] | 43.76±11.5 | D484-K666 [A,B,C,D,E,F] | 44.92±18.5 |
| | E497-K644 [A,B,C,D,E,F] | 70.66±49.4 | E497-K644 [A,B,C,D,E,F] | 42.03±22.2 |
| | E507-R505 [B,C,D,E,F] | 29.61±11.0 | E507-R505 [B,C,D,E,F] | 24.09±4.4 |
| | | | D582-K529 [C,D,F] | 29.82±12.6 |
| | E595-K603 [A,B,C,D,E,F] | 145.10±21.5 | E595-K603 [A,B,C,D,E,F] | 112.7±25.6 |
| | E605-R565 [A,B,C,D,E,F] | 68.89±30.3 | E605-R565 [B,C,D,E,F] | 66.43±16.9 |
| | E609-R565 [B,C,D,E] | 38.97±11.9 | E609-R565 [B,C,D,E,F] | 35.66±6.4 |
| | E609-R572 [B,C,D,E] | 59.7±27.8 | E609-R572 [B,C,D,E,F] | 40.30±8.0 |
| | D611-R641 [B,C,E] | 19.71±0.1 | D611-R641 [B,C,E,F] | 19.59±4.1 |
| | D620-K517 [B,C,E] | 92.91±15.4 | D620-K517 [A,B,C,D,E,F] | 36.22±18.1 |
| | D620-R572 [B,D,F] | 45.79±10.6 | D620-R572 [B,C,D,E,F] | 36.90±12.6 |

| | | | | |
|---|---|---|---|---|
| | E633-R630 [A,B,E] | 18.89±1.6 | E633-R630 [A,C,D] | 14.81±3.8 |
| | D635-R591 [A,E,F] | 23.41±8.3 | | |
| | D635-K603 [A,B,D] | 22.71±3.6 | D635-K603 [A,B,E,F] | 25.14±9.7 |
| | E653-K681 [A,B,C,D,E,F] | 41.19±3.7 | E653-K681 [A,B,C,D,E,F] | 40.56±5.0 |
| | E657-R677 [A,B,C,D,E,F] | 16.57±1.8 | E657-R677 [A,B,C,D,E,F] | 17.37±4.8 |
| | D697-R704 [A,B,C,D,E] | 62.59±35.3 | D697-R704 [A,B,C,D,E] | 56.14±13.6 |
| | E701-R704 [A,B,D,E] | 25.07±20.2 | | |
| | E701-R733 [A,B,C,D,E] | 20.10±9.2 | E701-R733 [A,B,C,D,E] | 17.48±3.7 |
| | E705-R720 [A,B,C,E] | 20.60±10.3 | E705-R720 [A,B,C,D,E] | 17.46±5.4 |
| | E724-R678 [A,B,C,E] | 15.15±2.7 | E724-R678 [B,C,D,E] | 13.97±2.3 |
| | D726-R720 [A,B,C,D] | 32.19±14.4 | D726-R720 [A,B,C,D] | 19.19±9.8 |
| | D755-K644 [B,C,D,F] | 48.83±20.6 | D755-K644 [B,C,D,E] | 49.54±18.1 |

**Table S19.** The list of features that were identified as important in classifying WT and the mutant systems for **the COMPLEX state and the APO state**. The features indicated in bold are found to be important in both the states. The locations of these features in the monomeric structure are depicted in Figure 5.

| COMPLEX State | | APO state | |
|---|---|---|---|
| **Feature** | **Descriptors** | **Feature** | **Descriptors** |
| **H1** | COULOMB<br>CONTACTS | **H1** | VDW<br>COULOMB<br>CONTACTS |
| H5 | VDW<br>COULOMB | H3 | RSA<br>VDW |

| | | | |
|---|---|---|---|
| | | | COULOMB |
| **H10** | VDW<br>CONTACTS | **H10** | VDW<br>CONTACTS |
| **H11** | RSA<br>CONTACTS | **H11** | COULOMB<br>CONTACTS |
| L3 | RSA<br>VDW<br>COULOMB | L1 | COULOMB<br>CONTACTS |
| L9 | RSA<br>VDW | L5 | RSA<br>COULOMB |
| L13 | RSA<br>VDW<br>COULOMB<br>CONTACTS | L7 | VDW<br>COULOMB |
| | | L14 | RSA<br>VDW |
| | | L15 | VDW<br>CONTACTS |
| **L16** | VDW<br>COULOMB<br>CONTACTS | **L16** | VDW<br>CONTACTS |

**Table S20.** The list of features that were identified as important in classifying **COMPLEX to APO transition** in the WT and the mutant systems. The features indicated in bold are found to be important in both the systems. The locations of these features in the monomeric structure are depicted in Figure 6.

| **WT** | | **R591S** | |
|---|---|---|---|
| **Feature** | **Descriptors** | **Feature** | **Descriptors** |
| H3 | COULOMB<br>VDW<br>RSA | H1 | COULOMB<br>VDW<br>RSA<br>CONTACTS |
| H5 | COULOMB<br>VDW | H2 | VDW<br>RSA |
| **H10** | COULOMB<br>VDW<br>CONTACTS | **H10** | COULOMB<br>VDW<br>RSA<br>CONTACTS |
| H12 | COULOMB | L3 | VDW |

| | | | |
|---|---|---|---|
| | VDW | | RSA<br>CONTACTS |
| **L11** | COULOMB<br>VDW<br>RSA | **L11** | COULOMB<br>RSA<br>CONTACTS |
| L14 | COULOMB<br>VDW | L13 | VDW<br>RSA<br>CONTACTS |
| L16 | COULOMB<br>VDW<br>RSA<br>CONTACTS | L15 | VDW<br>CONTACTS |
| | | B5 | COULOMB<br>RSA<br>CONTACTS |

**Table S21.** The chosen $c_{ij}$ cutoffs used for the dynamic graph networks of each system.

| | **WT: Monomer** | **R591S: Monomer** | **R591S: Hexamer** |
|---|---|---|---|
| COMPLEX | 0.6 | 0.5 | 0.6 |
| NUC | 0.5 | 0.5 | - |
| SUB | 0.6 | 0.6 | - |
| APO | 0.5 | 0.6 | 0.6 |

**Table S22.** The identified **most central (Top 10%) residues** for the **Monomer** networks of the **WT systems**. The bolded residues have been proposed as functionally/allosterically important, based on the analysis of the cryo-EM structures and on activity assays [10]. These positions are represented in Figure S15.

| **SETUP** | **Top 10% $C_B$<br>Monomer Networks** |
|---|---|
| WT:<br>COMPLEX | **I469**, L495, L521, **N527**, **G528**, L532, A535, V536, E539, S541, A542, T543, I547, S548, L552, **D582**, **Q583**, A627, V648, L650, D652, S687, D690, A693, D697, A699, E701, I734, R736 |
| WT:<br>NUCLEOTIDE | **V466**, I469, L521, **N527**, L531, L532, R534, F544, N546, **E561**, L563, F580, **D585**, A627, V648, L650, P651, R656, S687, G688, D690, T692, K696, L700, R704, L731, V738, **L743**, **Y746** |
| WT:<br>SUBSTRATE | L520, L521, **K529**, L532, S548, A550, L552, T553, L563, **D582**, L602, A627, R645, V648, L650, D652, E653, R656, D690, A693, K696, L700, R704, S737, |

| | |
|---|---|
| | **A739**, **L743**, **Y746**, **S750**, **G754** |
| WT:<br>APO | **I469**, A491, L495, I500, V504, L521, **F522**, **N527**, L532, I547, L563, **D582**, **V584**, L587, L602, E605, V608, D611, A627, T628, V648, P651, R656, L659, S687, D690, A693, D697, L731 |

**Table S23.** The identified **most central (Top 10%) residues** for the **Monomer** networks of the **R591S systems**. The bolded residues have been proposed as functionally/allosterically important, based on the analysis of the cryo-EM structures and on activity assays [10]. These positions are represented in Figure S15.

| SETUP | Top 10% $C_B$<br>Monomer Networks |
|---|---|
| R591S: COMPLEX | **D471**, V474, I500, **N527**, **K529**, **T530**, L532, R534, V536, F544, A549, T553, M574, S577, F580, **D582**, L602, A627, L650, P651, D652, S687, G688, D690, T692, K696, L700, R704, I734 |
| R591S:<br>NUCLEOTIDE | L495, **N527**, **K529**, L531, L532, R534, F544, **E561**, L563, A566, V570, M574, S577, E605, V608, A627, V648, P651, R656, S687, S689, D690, T692, K696, T723, Q725, H728, L731, V738, **L743** |
| R591S:<br>SUBSTRATE | **I469**, I500, L521, **F522**, **N527**, **K529**, L532, V536, C540, I547, A549, T553, **K555**, **D582**, **A598**, **R601**, A627, V648, S649, L650, P651, D652, T655, L659, S687, D690, A693, D697, V738 |
| R591S:<br>APO | **I469**, **E472**, Q488, K492, Q496, L501, **K529**, L532, R534, N546, I547, **E561**, L563, **D582**, A627, V648, L650, D652, T655, S687, D690, A693, K696, L700, I703, R735, S737, V738 |

**Table S24.** The optimal paths (Djikstra's paths) from the **Allosteric Center (S589/591)** to the **CT-Helix (W749/Y753)** for each state of the **WT and R591S monomer**. The residues are indicated as (chain):(amino acid ID)(residue no.). The path lengths are provided under each state name in the order of (WT/R591S). This demonstrates the network connectivity between the indicated regions between the NBD and the HBD as evaluated in our previous publication. It indicates how the propagation from the allosteric center is altered due to its mutation.

| State | Monomer:<br>Allosteric Center to CT-Helix | |
|---|---|---|
| (WT/R591S) | WT | R591S |
| COMPLEX<br>(2.3/2.5) | S589 - **D585** - **Q583** - T628 - **R630** - **S745** - **W749** | S589 - **D635** - A638 - F642 - R645 - **Y753** |
| NUCLEOTIDE<br>(4.1/2.2) | S589 - **D585** - A627 - L521 - V648 - **N527** - S687 - V738 - **L743** - **Y746** - **W749** | S589 - **L634** - **Q632** - **R630** - **Y746** - **W749** |

| | | |
|---|---|---|
| SUBSTRATE (2.3/3.4) | S589 - **D585** - A627 - L520 - R645 - **G754** - **W749** | S589 - **D585** - T628 - **F522** - V648 - L650 - S687 - V738 - **L743** - **Y746** - **W749** |
| APO (5.1/3.3) | S589 - **D585** - T628 - **F522** - V648 - **N527** - S687 - V738 - **740** - **L743** - **Y746** - **W749** | S589 - **D585** - T628 - **F522** - V648 - L650 - S687 - V738 - **L743** - **Y746** - **W749** |

**Table S25.** The optimal paths (Djikstra's paths) from the **ATP binding pocket** (P525/T530) to the **CTT binding region**:PL2 (H596/R601) for each state of the **WT and R591S monomer**. The residues are indicated as (chain):(amino acid ID)(residue no.). The path lengths are provided under each setup name in the order of (WT/R591S). This demonstrates the network connectivity between the indicated regions within the NBD. It indicates how the propagation between the ligand binding regions is altered due to the point mutation of the allosteric center.

| **State** | **Monomer: ATP Pocket to CTT Binding Region** | |
|---|---|---|
| **(WT/R591S)** | **WT** | **R591S** |
| COMPLEX (3.1/2.9) | **T530** - R534 - T538 - S541 - T543 - L545 - **I547** - L552 - L563 - **E561** - E605 - **R601** | **T530** - F580 - **I547** - L552 - S554 - **A598** - **H596** |
| NUCLEOTIDE (2.9/3.3) | **T530** - R534 - N546 - **I469** - L563 - **E561** - **R601** | **T530** - R534 - F544 - S577 - M574 - V570 - A566 - L563 - **G560** - **A598** - **H596** |
| SUBSTRATE (2.4/2.9) | **T530** - R534 - F544 - N546 - S548 - L552 - S554 - **A598** - **R601** | **T530** - L532 - **K529** - L521 - A627 - **D582** - A549 - T553 - **K555** - **A598** - **R601** |
| APO (3.0/2.0) | **P525** - V648 - L521 - A627 - **V584** - L587 - K603 - **R601** | **T530** - R534 - N546 - E472 - **I469** - L563 - **E561** - **R601** |

**Table S26.** The identified **most central (Top 10%) residues** for the **Hexamer** network of the **R591S COMPLEX state**. The bolded residues have been proposed as functionally/allosterically important, based on the analysis of the cryo-EM structures and on activity assays [10]. These positions are represented in Figure S19a.

| **Chain** | **Top 10% $C_B$ Hexamer Networks R591S: COMPLEX** |
|---|---|
| A (20) | L520, **F522**, **P524**, A549, T553, S554, **K555**, **D559**, L563, **D582**, **V584**, S589, L625, V646, V648, L650, D652, R656, S687, I734 |
| B (**42**) | **F522**, **N527**, S548, A549, **K555**, **Y556**, **V557**, **D559**, **G560**, **K562**, L563, A566, A569, R572, **I581**, S586, **A598**, **R600**, **R601**, L625, T628, **E636**, L639, **R640**, F642, R645, V648, L650, P651, R656, Y686, G688, D690, L700, I703, R735, V738, **L743**, **Y746**, |

| | |
|---|---|
| | **W749**, **D752**, **Y753** |
| C<br>(30) | L519, L520, L521, I547, L552, T553, S554, **K555**, **Y556**, **V557**, **E561**, F580, **R601**, K603, E605, L625, R645, Y647, V648, L650, R656, L694, L731, R735, V738, **L743**, **Y746**, **W749**, **Y753** |
| D<br>(36) | F509, R513, P515, G518, L520, L521, **F522**, **P524**, R534, F544, N546, S548, **K555**, **Y556**, **V557**, **K562**, A566, V570, F580, **Q583**, **R601**, K603, A638, F642, R645, Y647, V648, P651, R656, S687, D690, A693, K696, L700, **W749**, **Y753** |
| E<br>(29) | G518, L520, L521, **F522**, **P524**, **G526**, T553, S554, **Y556**, **V557**, **G558**, **D559**, **E561**, **K562**, L563, L567, **R601**, K603, T604, A638, F642, R645, Y647, D690, L691, A693, L694, K696, L700 |
| F<br>(**23**) | L495, L520, L521, **N527**, **T530**, L545, I547, **E561**, **K562**, L567, I579, F580, K603, L625, **A637**, **R640**, V648, L660, G688, L691, L694, A695, A699 |

**Table S27.** The identified **most central (Top 10%) residues** for the **Hexamer** network of the **R591S APO state**. The bolded residues have been proposed as functionally/allosterically important, based on the analysis of the cryo-EM structures and on activity assays [10]. These positions are represented in Figure S19b.

| **Chain** | **Top 10% $C_B$**<br>**Hexamer Networks**<br>**R591S: APO** |
|---|---|
| A<br>(**25**) | I485, Q488, K492, L519, L520, L521, **F522**, **P524**, **N527**, L531, R534, F568, I622, V624, L625, K644, V646, V648, L650, P651, T683, Y686, S730, I734, R735 |
| B<br>(27) | L519, L520, L521, **K529**, I547, A549, A571, F580, V623, L625, **R640**, F642, K644, R645, V648, L650, Y686, H728, L731, R735, V738, **L743**, **Y746**, **E747**, **W749**, **S750**, **Y753** |
| C<br>(28) | Q488, L519, L521, A550, L552, S554, **Y556**, **G560**, F580, E605, A626, A638, F642, R645, V646, V648, L650, P651, R656, Y686, R733, R735, S737, **A739**, **S742**, **S745**, **W749**, **Y753** |
| D<br>(**44**) | E472, V474, L495, L519, L520, **P524**, **G526**, **G528**, L532, C540, A542, F544, N546, S548, T553, **K555**, **V557**, F580, S586, E605, L625, A626, T628, R645, V646, V648, L650, Y686, D690, L694, L731, R735, V738, **P740**, **S742**, **L743**, **N744**, **S745**, **Y746**, **K748**, **W749**, **S750**, **Y753**, **G754** |
| E<br>(38) | L519, L520, L521, N546, I547, S548, S551, **K562**, R565, F568, A569, A571, I579, F580, E605, L607, V624, L625, A638, F642, R645, V646, Y647, V648, S649, L650, P651, R656, Y686, S687, D690, L694, A698, R735, V738, **L743**, **K748**, **Y753** |
| F<br>(18) | A491, L495, L521, **K529**, **K562**, E605, L625, A627, Y647, L650, R656, S730, I734, R736, V738, **L743**, **E747**, **Q751** |

**Table S28.** The optimal paths (Djikstra's paths) from the **allosteric center (S589/591)** to the **CT-Hlx (W749/Y753)** in the Hexamer WT (previously reported[10]) and R591S **COMPLEX state**. The residues are indicated as (chain):(amino acid ID)(residue no.). The path lengths are provided under each setup name in the order of (WT/R591S). This demonstrates the network connectivity between the indicated regions between the NBD and the HBD as evaluated in our previous publication.

| **Chain (WT/R591S)** | **Hexamer COMPLEX: Intra Protomer Allosteric Center to CT Hlx** | |
|---|---|---|
| | **WT** | **R591S** |
| A (0.4/0.4) | A:S589 - A:**P631** - A:**W749** | A:S589 - A:**L634** - A:**Q632** - A:**W749** |
| B (0.6/0.5) | B:S589 - B:**D635** - B:L639 - B:**Y753** | B:S589 - B:**D635 -** B:L639 - B:**Y753** |
| C (0.7/0.6) | C:S589 - C:**Q632** - C:**W749** | C:S589 - C:**Q632** - C:**W749** |
| D (0.8/1.0) | D:S589 - D:**Q632** - D:**W749** | D:S589 - D:**D585** - D:**R630** - D:**W749** |
| E (0.8/1.1) | E:S589 - E:**Q632** - E:**W749** | E:S589 - E:**L634** - E:**Q632** - E:**W749** |
| F (0.5/0.6) | F:S589 - F:**Q632** - F:**W749** | F:S589 - F:**R630** - F:**Y746** - F:**W749** |

**Table S29.** The optimal paths (Djikstra's paths) from the **allosteric center (S589/591)** to the **CT-Hlx (W749/Y753)** in the Hexamer WT (previously reported[10]) and R591S **APO state**. The residues are indicated as (chain):(amino acid ID)(residue no.). The path lengths are provided under each setup name in the order of (WT/R591S). This demonstrates the network connectivity between the indicated regions between the NBD and the HBD as evaluated in our previous publication.

| **Chain (WT/R591S)** | **Hexamer APO: Intra Protomer Allosteric Center to CT Hlx** | |
|---|---|---|
| | **WT** | **R591S** |
| A (0.6/0.8) | A:S589 - A:**D585** - A:T628 - A:**F522** - A:**Y746** - A:**W749** | A:S589 - A:**L634** - A:**Q632** - A:**W749** |
| B (0.4/0.6) | B:S589 - B:**D635** - B:L639 - B:**Y753** | B:S589 - B:**P631** - B:**W749** |
| C | C:S589 - C:**Q632** - C:**W749** | C:S589 - C:**Q632** - C:**W749** |

| | | |
|---|---|---|
| (0.8/1.0) | | |
| D<br>(0.7/1.4) | D:S589 - D:**Q632** - D:**W749** | D:S589 - D:**D585** - D:**R630** - D:**W749** |
| E<br>(0.9/0.9) | E:S589 - E:**D635** - E:L639 - E:**Y753** | E:S589 - E:**D635** - E:L639 - E:**Y753** |
| F<br>(0.3/0.7) | F:S589 - F:**P631** - F:**W749** | F:S589 - F:**Q632** - F:**W749** |

**Table S30.** The optimal paths (Djikstra's paths) from the **allosteric center (S589/591)** to the **CT-Hlx (W749/Y753)** in the Hexamer WT (previously reported[10]) and R591S **COMPLEX state**. The residues are indicated as (chain):(amino acid ID)(residue no.). The path lengths are provided under each setup name in the order of (WT/R591S). This demonstrates the network connectivity between the domains of neighboring protomers as evaluated in our previous publication.

| **Chain (WT/R591S)** | **Hexamer COMPLEX: Inter Protomer Allosteric Center to CT Hlx** | |
|---|---|---|
| | **WT** | **R591S** |
| B to A<br>(0.7/0.8) | B:S589 - B:**E636** - A:**P524** - A:**Y746** - A:**W749** | B:S589 - B:**E636** - A:**P524** - A:**Y746** - A:**W749** |
| C to B<br>(1.3/1.7) | C:S589 - C:**E636** - B:**P524** - B:**Y746** - B:**W749** | C:S589 - C:**D635** - C:L639 - C:**Y753** - B:R735 - B:V738 - B:**L743** - B:**Y746** - B:**W749** |
| D to C<br>(2.3/2.5) | D:S589 - D:A638 - D:**R640** - C:**G526** - C:**P524** - C:Y647 - C:**W749** | D:S589 - D:**V584** - D:L520 - D:R645 - D:**Y753** - C:R735 - C:V738 - C:**L743** - C:**Y746** - C:**W749** |
| E to D<br>(2.3/2.5) | E:S589 - E:A638 - E:**R640** - D:**G526** - D:G523 - D:**Y746** - D:**W749** | E:S589 - E:**L634** - E:A638 - E:T604 - D:**Q583** - D:**R630** - D:**W749** |
| F to E<br>(1.2/1.2) | F:S589 - F:**D635** - F:**A637** - E:**P524** - E:**Y746** - E:**W749** | F:S589 - F:**E636** - E:**P524** - E:**Y746** - E:**W749** |
| A to F<br>(5.6/4.0) | A:S589 - A:L587 - A:L552 - A:S554 - B:**V557** - B:**K555** - C:**V557** - C:**K555** - D:**Y556** - D:T553 - E:**R601** - E:K603 - E:L588 - E:**L634** - E:**Q632** - E:**Y746** - E:**L743** - E:V738 - E:R736 - F:**Y753** | A:S589 - B:**R600** - B:**A598** - B:**Y556** - C:**V557** - C:**K555** - D:**Y556** - E:**V557** - E:S554 - F:**K562** - F:L567 - F:I579 - F:L625 - F:L521 - F:Y647 - F:**W749** |

**Table S31.** The optimal paths (Djikstra's paths) from the **allosteric center (S589/591)** to the **CT Hlx (W749/Y753)** in the Hexamer WT (previously reported[10]) and R591S **APO state**. The residues are indicated as (chain):(amino acid ID)(residue no.). The path lengths are provided under each setup name in the order of (WT/R591S). This demonstrates the network connectivity between the domains of neighboring protomers as evaluated in our previous publication.

| **Chain (WT/R591S)** | **Hexamer APO: Inter Protomer Allosteric Center to CT Hlx** | |
|---|---|---|
| | **WT** | **R591S** |
| B to A (0.7/1.2) | B:S589 - B:**E636** - A:**P524** - A:**Y746** - A:**W749** | B:S589 - B:**E636** - A:**P524** - A:**Y746** - A:**W749** |
| C to B (1.2/2.2) | C:S589 - C:**E636** - B:**P524** - B:F522 - B:R645 - B:**Y753** | C:S589 - C:K603 - C:E605 - B:L549 - B:I547 - B:F580 - B:L625 - B:L520 - B:R645 - B:**Y753** |
| D to C (1.8/2.9) | D:S589 - D:**Q632** - D:**Y753** - C:S737 - C:**A739** - C:**S742** - C:**S745** - C:**W749** | D:S589 - D:**D585** - D:A626 - D:L520 - D:R645 - D:**G754** - C:R735 - C:S737 - C:**A739** - C:**S742** - C:**S745** - C:**W749** |
| E to D (1.2/2.2) | E:S589 - E:**E636** - D:**N629** - D:**Y746** - D:**W749** | E:S589 - E:**D635** - E:L639 - E:**Y753** - D:V738 - D:**P740** - D:**N744** - D:**W749** |
| F to E (1.6/2.1) | F:S589 - F:**P631** - F:**W749** - F:**D752** - E:S737 - E:**A739** - E:**S742** - E:**S745** - E:**W749** | F:S589 - F:**R630** - F:**L746** - F:**Q751** - E:R735 - E:V738 - E:**L743** - E:**Y746** - E:**W749** |
| A to F (3.7/7.1) | A:S589 - B:**R600** - B:L602 - B:T553 - C:**V557** - C:**K555** - D:**Y556** - E:**V557** - E:T553 - F:**R601** - F:K603 - F:L587 - F:**D585** - F:**P631** - F:**W749** | A:S589 - A:**D635** - A:L639 - A:L520 - A:V646 - A:V648 - A:L650 - A:Y686 - A:R735 - B:**S750** - B:**E747** - B:**L743** - B:V738 - B:R735 - C:**Y753** - C:**W749** - C:**S745** - C:**S742** - C:**A739** - C:S737 - C:R735 - D:**Y753** - D:**K748** - D:**N744** - D:**P740** - D:V738 - E:**Y753** - E:**K748** - E:**L743** - E:V738 - E:R735 - F:**Y753** |

**Table S32.** The optimal paths (Djikstra's paths) from the **ATP binding pocke**t (P525/T530) to **CTT binding region**:PL2 (H596/R601) in the **Hexamer** WT and R591S **COMPLEX state**. The residues are indicated as (chain):(amino acid ID)(residue no.). The path lengths are provided under each setup name in the order of (WT/R591S). This demonstrates the network connectivity between the indicated regions within the NBD. It indicates how the propagation between major binding regions is altered due to the point mutation of the allosteric center.

| **Chain (WT/R591S)** | **Hexamer COMPLEX: Intra Protomer ATP Pocket to CTT Binding Region** | |
|---|---|---|
| | **WT** | **R591S** |
| A | A:**P525** - A:**F522** - A:L520 - A:F642 - | A:**P525** - A:**F522** - A:L520 - A:F642 - |

| | | |
|---|---|---|
| (0.6/0.7) | A:A638 - A:K603 - A:**R601** | A:A638 - A:K603 - A:**R601** |
| B (1.3/1.1) | B:**P525** - B:**F522** - B:L520 - B:F642 - B:A638 - B:T604 - B:**R601** | B:**P525** - B:**G523** - B:Y647 - B:R645 - B:F642 - B:A638 - B:T604 - B:**R601** |
| C (2.4/1.4) | C:**P525** - C:**G523** - C:L521 - C:F642 - C:A638 - C:L607 - C:E605 - C:**R601** | C:**P525** - C:**F522** - C:L520 - C:F642 - C:A638 - C:K603 - C:**R601** |
| D (2.1/1.8) | D:**T530** - D:R534 - D:F544 - D:A571 - D:A566 - D:**K562** - D:**R601** | D:**P525** - D:V648 - D:L521 - D:F642 - D:A638 - D:K603 - D:**R601** |
| E (1.9/1.6) | E:**T530** - E:L521 - E:T628 - E:**D585** - E:L588 - E:K603 - E:**R601** | E:**P525** - E:**F522** - E:L520 - E:F642 - E:A638 - E:K603 - E:**R601** |
| F (0.8/0.9) | F:**P525** - F:T628 - F:**V584** - F:L587 - F:K603 - F:**R601** | F:**T530** - F:A627 - F:**V584** - F:L587 - F:K603 - F:**R601** |

**Table S33.** The optimal paths (Djikstra's paths) from the **ATP binding pocke**t (P525/T530) to **CTT binding region**:PL2 (H596/R601) in the **Hexamer** WT and R591S **APO state**. The residues are indicated as (chain):(amino acid ID)(residue no.). The path lengths are provided under each setup name in the order of (WT/R591S). This demonstrates the network connectivity between the indicated regions within the NBD. It indicates how the propagation between major binding regions is altered due to the point mutation of the allosteric center.

| **Chain (WT/R591S)** | **Hexamer APO: Intra Protomer ATP Pocket to CTT Binding Region** | |
|---|---|---|
| | **WT** | **R591S** |
| A (0.7/1.0) | A:**T530** - A:L521 - A:K644 - A:F642 - A:A638 - A:K603 - A:**R601** | A:**T530** - A:L521 - A:K644 - A:F642 - A:A638 - A:K603 - A:**R601** |
| B (0.9/1.1) | B:**P525** - B:T628 - B:**V584** - B:L588 - B:K603 - B:**R601** | B:**T530** - B:L521 - B:F642 - B:A638 - B:K603 - B:**R601** |
| C (1.5/1.7) | C:**P525** - C:T628 - C:**L634** - C:A638 - C:T604 - C:**R601** | C:**P525** - C:**F522** - C:L520 - C:F642 - C:A638 - C:K603 - C:**R601** |
| D (1.4/3.1) | D:**P525** - D:T628 - D:**D585** - D:S589 - D:K603 - D:**R601** | D:**P525** - D:**G523** - D:**Y746** - D:**Q632** - D:**D635** - D:E590 - D:S592 - D:**S594** - D:**H596** |
| E (1.4/1.8) | E:**T530** - E:A533 - E:I579 - E:F568 - E:F606 - E:**R601** | E:**T530** - E:A533 - E:I578 - E:A571 - E:F568 - E:R565 - E:E605 - E:**R601** |

| | | |
|---|---|---|
| F<br>(0.6/1.1) | F:**T530** - F:**D582** - F:**V584** - F:L587 - F:K603 - F:**R601** | F:**T530** - F:A627 - F:**V584** - F:L588 - F:K603 - F:**R601** |

**Table S34.** The optimal paths (Djikstra's paths) from the **ATP binding pocket** (P525/T530) in ***protomer i*** to **CTT binding region**:PL2 (H596/R601) in ***protomer i-1*** for the **Hexamer** WT and R591S **COMPLEX state**. The residues are indicated as (chain):(amino acid ID)(residue no.). The path lengths are provided under each setup name in the order of (WT/R591S). This demonstrates the network connectivity between the NBD of i with the i-1 protomers.

| **Chain (WT/R591S)** | **Hexamer COMPLEX: Inter Protomer ATP Pocket to CTT Binding Region** | |
|---|---|---|
| | **WT** | **R591S** |
| B to A<br>(1.5/1.5) | B:**P525** - B:F522 - B:L520 - B:F642 - B:**R640** - A:**P524** - A:F522 - A:L520 - A:F642 - A:A638 - A:K603 - A:**R601** | B:**P525** - B:**G523** - B:Y647 - B:R645 - B:F642 - B:**R640** - A:**P524** - A:F522 - A:L520 - A:F642 - A:A638 - A:K603 - A:**R601** |
| C to B<br>(3.2/2.7) | C:**P525** - C:**G523** - C:L521 - C:F642 - C:**R640** - B:**P525** - B:F522 - B:L520 - B:F642 - B:A638 - B:T604 - B:**R601** | C:**P525** - C:**G523** - C:Y647 - C:**Y753** - B:R735 - B:V738 - B:**L743** - B:**Y746** - B:**W749** - B:**Y753** - B:L639 - B:**A637** - B:T604 - B:**R601** |
| D to C<br>(3.8/3.1) | D:**T530** - D:A533 - D:I579 - D:L567 - D:L563 - D:**G560** - D:**Y556** - C:**S554** - C:**D559** - C:**E561** - C:**R601** | D:**T530** - D:R534 - D:F544 - D:S577 - D:H573 - D:V570 - D:A566 - D:**K562** - C:S554 - C:**V557** - C:**R601** |
| E to D<br>(3.5/2.7) | E:**T530** - E:L521 - E:T628 - E:**D585** - E:L588 - E:K603 - E:**R601** - D:T553 - D:**G560** - D:E605 - D:**R601** | E:**P525** - F:A638 - F:F606 - F:**E561** - E:S554 - E:**V557** - D:**Y556** - D:**A598** - D:**R601** |
| F to E<br>(2.8/1.7) | F:**P525** - F:F522 - F:L520 - F:**R641** - E:**G526** - E:**P524** - E:F522 - E:T628 - E:**D585** - E:L588 - E:K603 - E:**R601** | F:**T530** - F:F580 - F:L545 - F:L567 - F:**K562** - E:T553 - E:**E561** - E:**R601** |
| A to F<br>(5.7/3.4) | A:**T530** - A:**I581** - A:S548 - A:T553 - B:**V557** - B:**K555** - C:**V557** - C:**K555** - D:**Y556** - D:T553 - E:**R601** - E:K603 - E:L588 - E:**D585** - E:T628 - E:F522 - E:**P524** - F:**A637** - F:K603 - F:**R601** | A:**P525** - A:T628 - A:**V584** - B:**R601** - B:**A598** - B:**Y556** - C:**V557** - C:**K555** - D:**Y556** - E:**V557** - E:**K555** - E:T553 - F:**R601** |

**Table S35.** The optimal paths (Djikstra's paths) from the **ATP binding pocket** (P525/T530) ***protomer i*** to **CTT binding region**:PL2 (H596/R601) in ***protomer i-1*** for the **Hexamer** WT and R591S **APO state**. The residues are indicated as (chain):(amino acid ID)(residue no.). The path lengths are provided under each setup name in the order of (WT/R591S). This demonstrates the network connectivity between the NBD of i with the i-1 protomers.

| **Chain (WT/R591S)** | **Hexamer APO: Inter Protomer ATP Pocket to CTT Binding Region** | |
|---|---|---|
| | **WT** | **R591S** |
| B to A (1.4/1.9) | B:**P525** - B:T628 - B:**V584** - B:L588 - B:K603 - B:**R601** - B:**E597** - A:**H596** | B:**T530** - B:L521 - B:R645 - B:**Y753** - A:R735 - A:Y686 - A:L650 - A:V648 - A:V646 - A:K644 - A:F642 - A:A638 - A:K603 - A:**R601** |
| C to B (2.0/3.2) | C:**P525** - C:T628 - C:**L634** - C:**E636** - B:**N629** - B:**P631** - B:**L634** - B:L588 - B:K603 - B:**R601** | C:**P525** - C:**G523** - C:**Y746** - C:**W749** - C:**Y753** - B:R735 - B:Y686 - B:L650 - B:V648 - B:L520 - B:F642 - B:A638 - B:K603 - B:**R601** |
| D to C (2.3/3.8) | D:**T530** - D:A533 - D:I579 - D:L567 - D:**K562** - C:T553 - C:**A598** - C:**R601** | D:**P525** - D:**G523** - D:**S750** - D:**Y753** - C:R735 - C:Y686 - C:L650 - C:V648 - C:R645 - C:F642 - C:A638 - C:K603 - C:**R601** |
| E to D (2.3/3.2) | E:**T530** - E:T533 - E:I579 - E:F568 - E:R565 - E:**K562** - D:A550 - D:**G560** - D:**R601** | E:**T530** - E:A533 - E:I578 - E:A571 - E:A569 - E:A566 - E:**K562** - D:T553 - D:**K555** - D:**V557** - D:**A598** - D:**R601** |
| F to E (1.0/3.2) | F:**T530** - F:**D582** - F:**V584** - F:L587 - F:L602 - F:**A598** - E:**H596** | F:**P525** - F:**G523** - F:**E747** - F:**Q751** - E:R735 - E:Y686 - E:L650 - E:Y647 - E:R645 - E:F642 - E:A638 - E:L607 - E:T604 - E:**R601** |
| A to F (3.1/6.9) | A:**T530** - B:V608 - B:E605 - B:L602 - B:T553 - C:**V557** - C:**K555** - D:**Y556** - E:**V557** - E:T553 - F:**R601** | A:**P525** - A:S737 - B:**Y753** - B:**W749** - B:**Y746** - B:**L743** - B:V738 - B:R735 - C:**Y753** - C:**W749** - C:**S745** - C:**S742** - C:**A739** - C:S737 - C:R735 - D:**Y753** - D:**K748** - D:**N744** - D:**P740** - D:V738 - E:**Y753** - E:R645 - E:L520 - E:L625 - E:**F580** - E:N546 - E:S548 - F:**R601** |

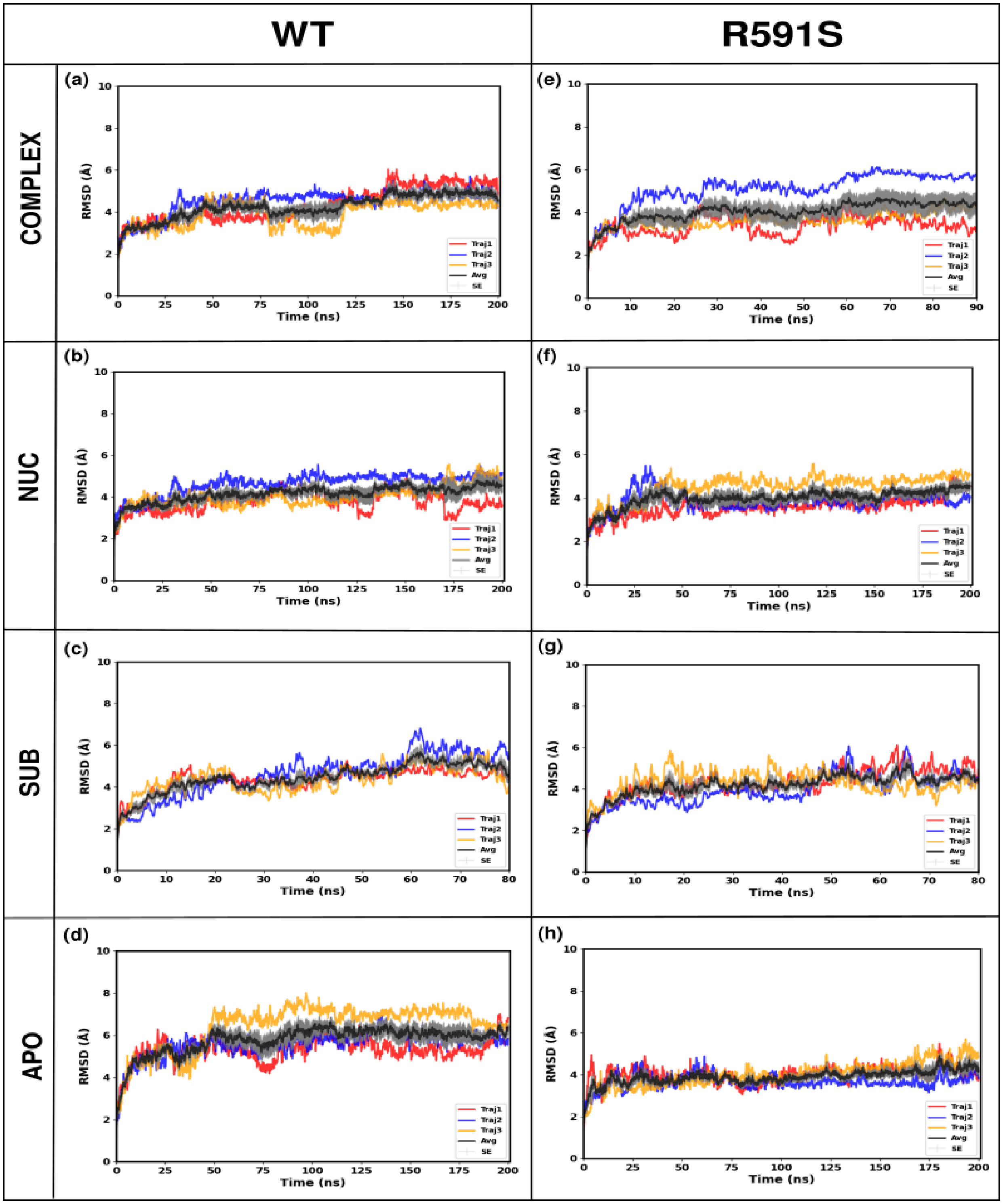


**Figure S1.** The **RMSD vs Time plots** for the 200 ns trajectories of the **Monomer systems**. a-d) The RMSD plots of the WT monomer states. (In the SUB state, the glutamate peptide dissociates after 80 ns. Therefore, we only use the first 80 ns of the simulation to represent the WT SUB state, and the RMSD values only up to 80 ns are in Fig. (c) ); e-h) The RMSD plots of the R591S monomer states. (In the mutant SUB state, the glutamate peptide dissociates after

80 ns, same as the WT system. So we have only shown the RMSDs up to 80 ns in Fig. (g); similarly, in the mutant COMPLEX state, the substrate dissociates after 90 ns, and we show the RMSD values only up to 90 ns in Fig. (e) ).

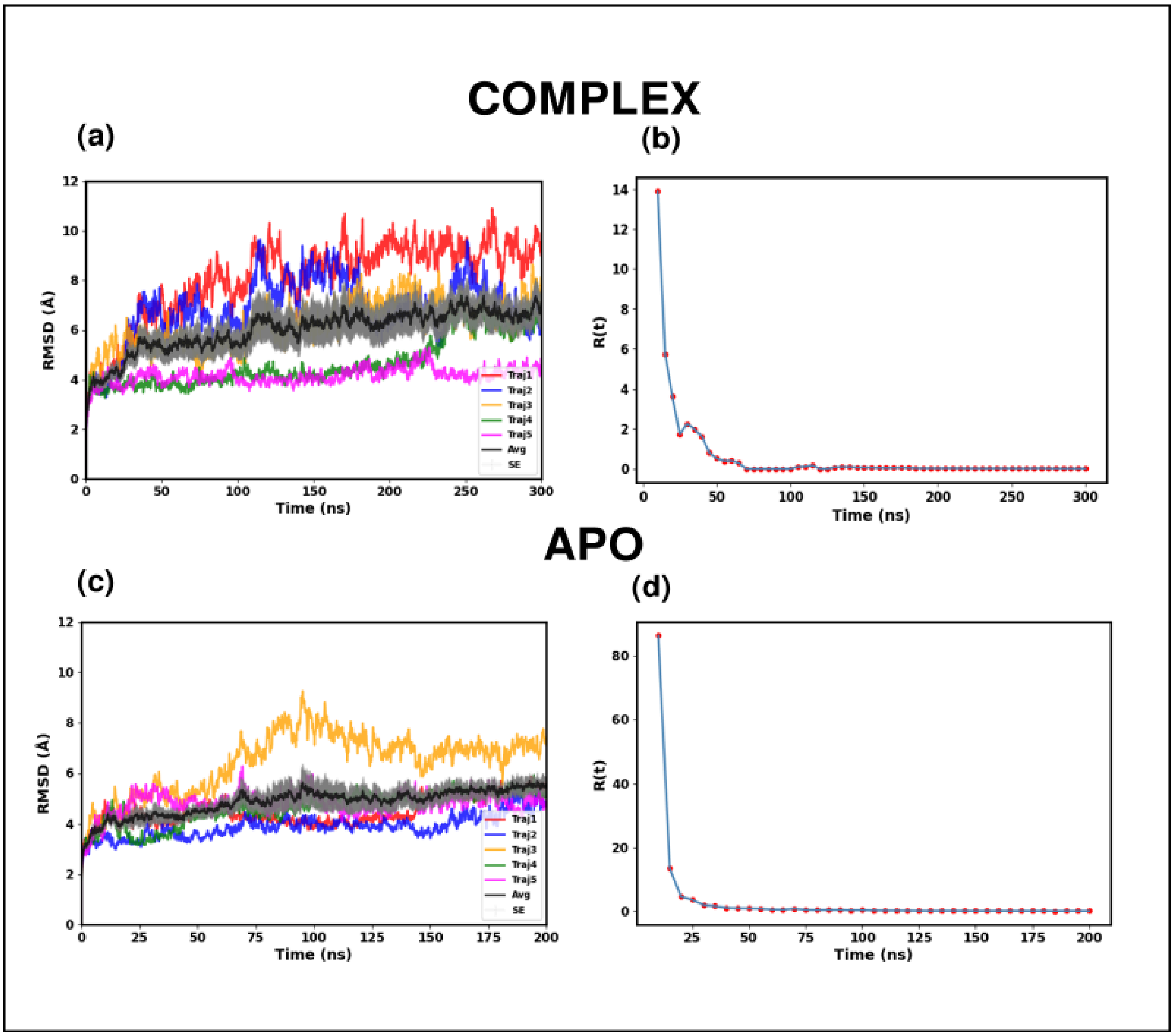


**Figure S2.** The **RMSD vs Time plots** and the **DCCM convergence plots** of **R591S mutated hexamer systems**. (a) RMSD plot of R591S mutated COMPLEX system (b) DCCM convergence plot of R591S mutated COMPLEX system (c) RMSD plot of R591S mutated APO system (d) DCCM convergence plot of R591S mutated APO system. In the COMPLEX state, the DCCM matrix converges on a timescale of 70 ns, while, in the APO state, the matrix converges on a timescale of 40 ns.

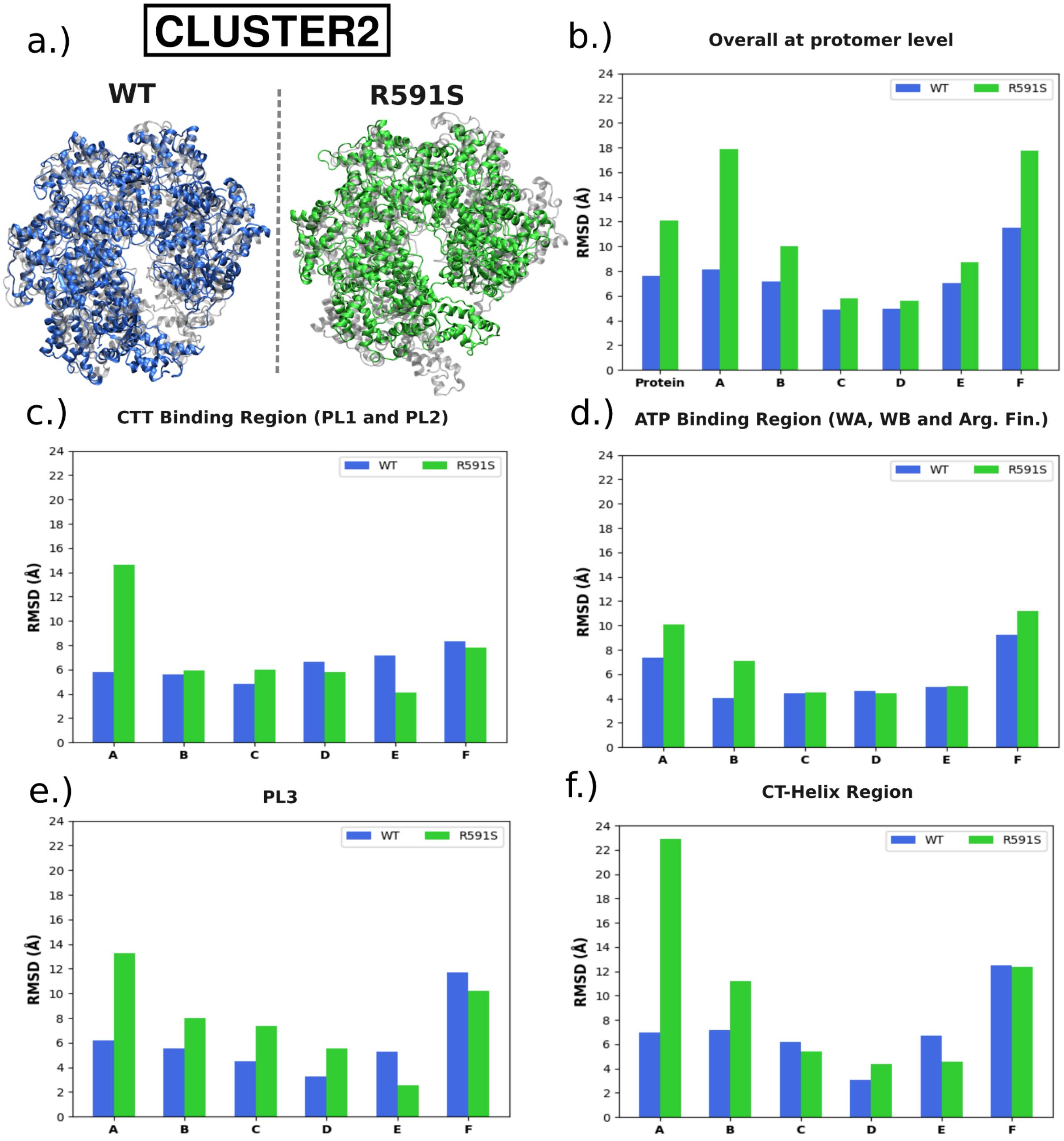


**Figure S3.** The RMSD values of the central structure of the APO state, aligned with the central structure of the COMPLEX state for **cluster 2**. (These values were obtained **by aligning the whole protein**) (a) Central structures of cluster2 obtained from the second clustering step for the WT and the mutant systems.(Grey indicates the COMPLEX structure, while blue and green indicate the APO structure of the WT and mutant systems, respectively) (b) Comparison of the entire protein and each protomer (c) Comparison of the CTT binding region (including both PL1 and PL2) (d) Comparison of the ATP binding region (including WA, WB and arg.fin) (e) Comparison of the PL3 region (f) Comparison of the CT-helix region.

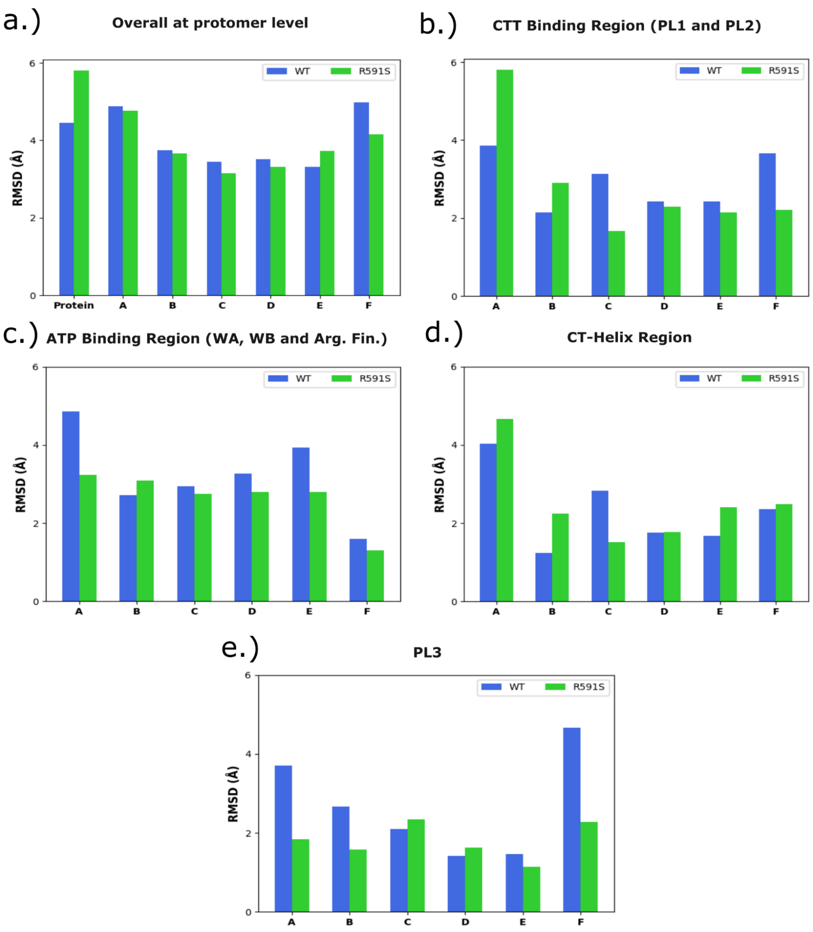


**Figure S4.** The RMSD values of the central structure of the APO state, aligned with the central structure of the COMPLEX state for **cluster 1**. (These values were obtained **by aligning the specific functional regions separately**) (a) Comparison of the entire protein and each protomer (b) Comparison of the CTT binding region (including both PL1 and PL2) (c) Comparison of the ATP binding region (including WA, WB and arg.fin) (d) Comparison of the CT-Helix region (e) Comparison of the PL3 region.

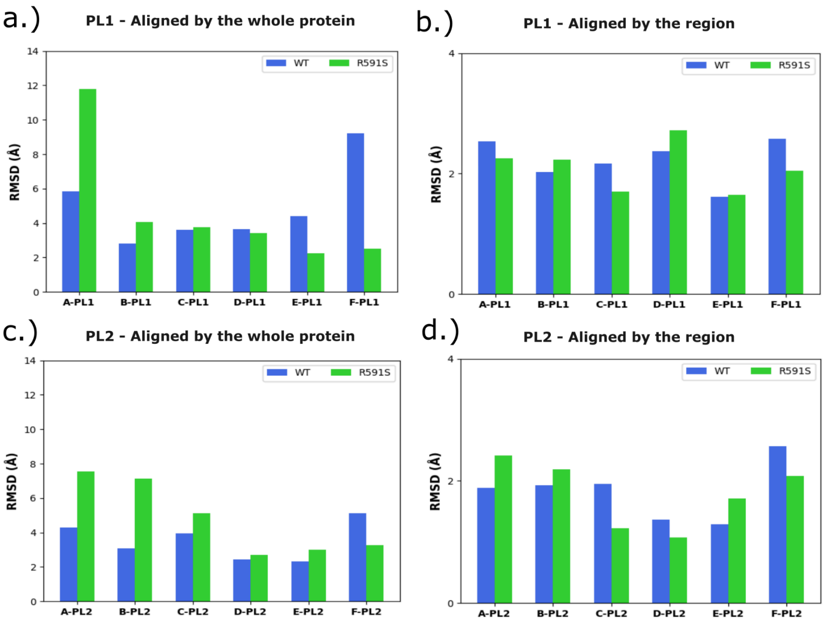


**Figure S5.** The RMSD values of the central structure of the APO state, aligned with the central structure of the COMPLEX state of **cluster 1** for **PL1 and PL2 regions**. (a) PL1 region aligned by the whole protein (b) PL1 region aligned by the region (c) PL2 region aligned by the whole protein (d) PL2 region aligned by the region

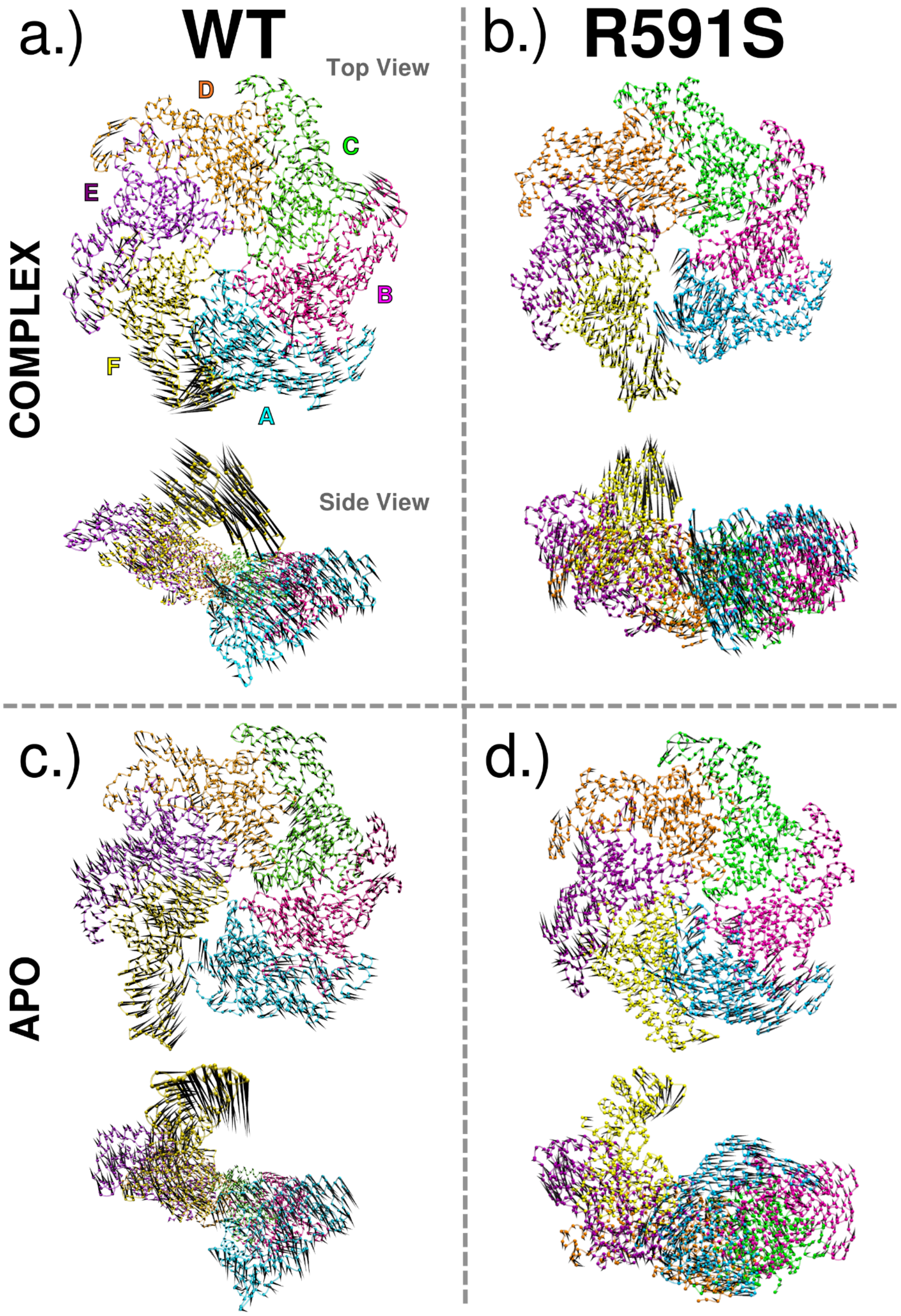


**Figure S6.** The porcupine plots illustrate the **global motions** corresponding to **PC2** (a) global motions of WT COMPLEX state (b) global motions of R591S COMPLEX state (c) global motions of WT APO state (d) global motions of R591S APO state.The protomer labels, as indicated in (a), are used consistently across the entire figure. A side view of the protein is also included to clearly display the motions of the terminal protomers, A and F. The variance covered by PC2 for each of these systems is provided in Table S8.

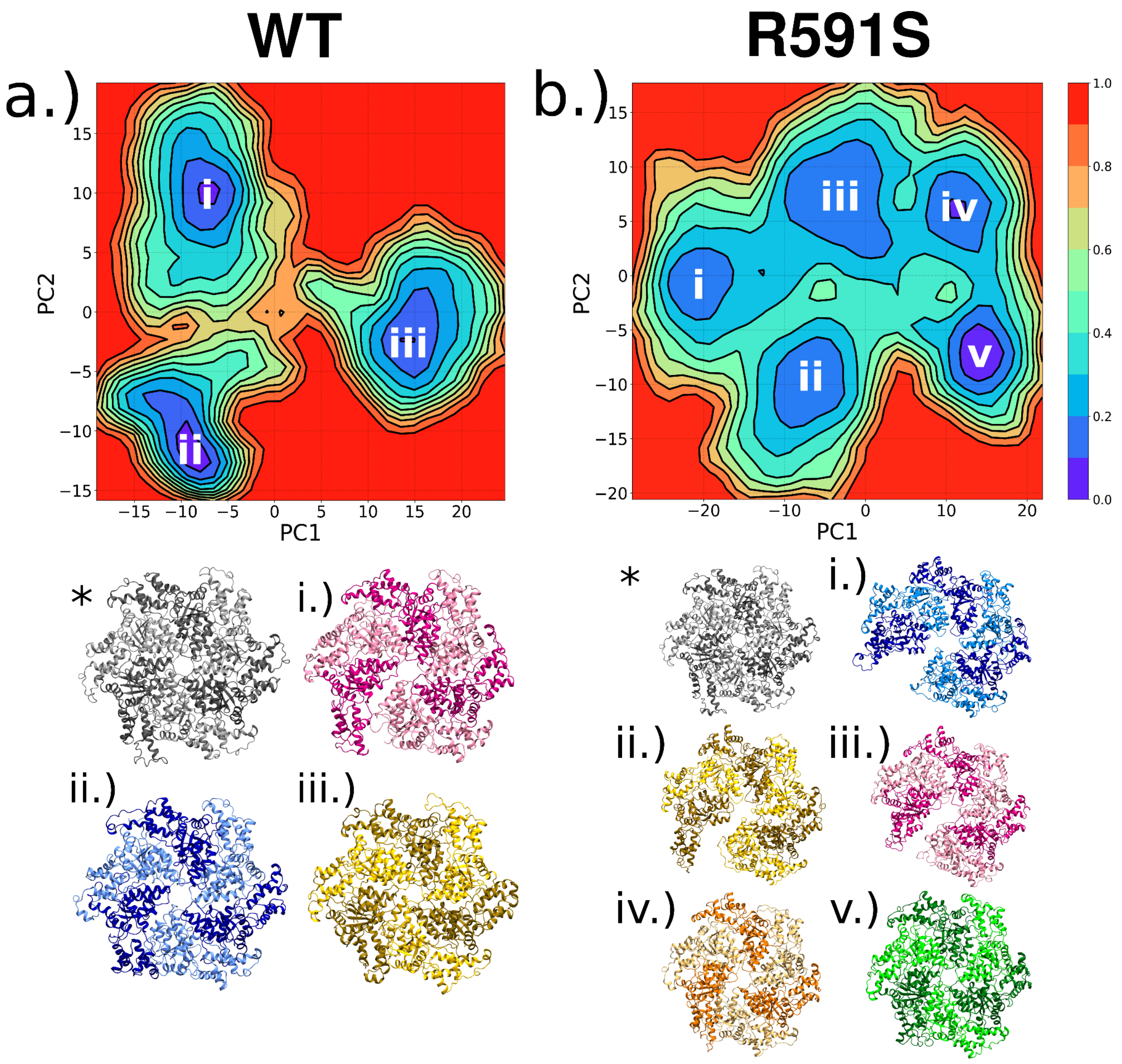


**Figure S7.** The free energy landscape in the PC1/PC2 subspace for the **COMPLEX state**. (a) For the WT system, we identified three minima and extracted the corresponding lowest-energy representative structures, labeled as i - iii (b) For the R591S system, we identified five minima and extracted the corresponding lowest-energy representative structures, labeled as i - v. The starting structure (*) is shown in gray. The accompanying backbone-RMSD between the minima structures and the starting structure are provided in Table S12. Note that the actual range of free energy values for the WT COMPLEX system is 0–3.7 kcal/mol, and for the mutant system, it is 0–4.02 kcal/mol. For visualization purposes, we have selected an arbitrary range of 0–1.

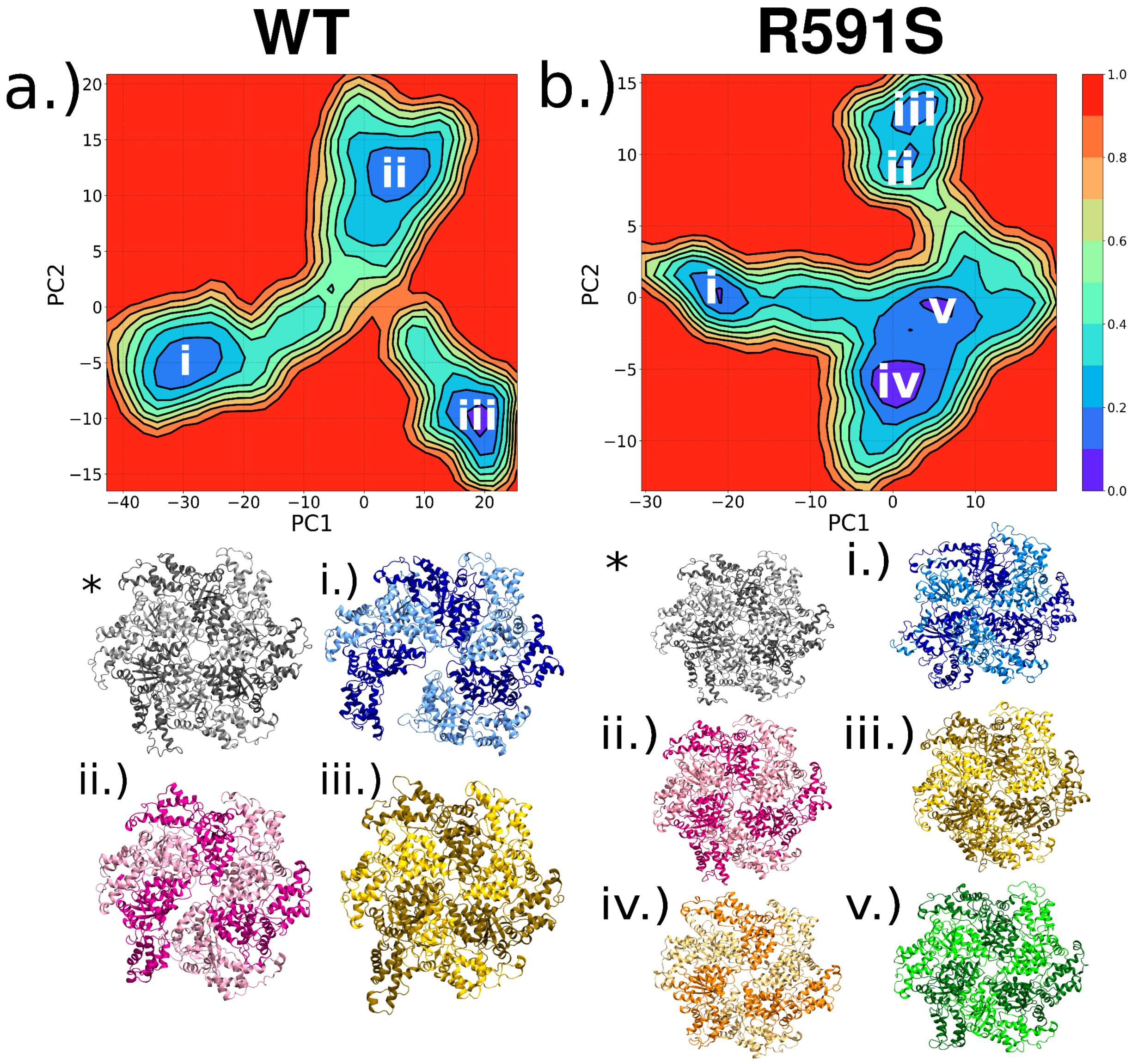


**Figure S8.** (a) The free energy landscape in the PC1/PC2 subspace for the **APO state**. (a) For the WT system, we identified three minima and extracted the corresponding lowest-energy representative structures, labeled as i - iii (b) For the R591S system, we identified five minima and extracted the corresponding lowest-energy representative structures, labeled as i - v. The starting structure (*) is shown in gray. The accompanying backbone-RMSD between the minima structures and the starting structure are provided in Table S13. Note that the actual range of free energy values for the WT APO system is 0–4.02 kcal/mol, and for the mutant system, it is 0–4.09 kcal/mol. For visualization purposes, we have selected an arbitrary range of 0–1.

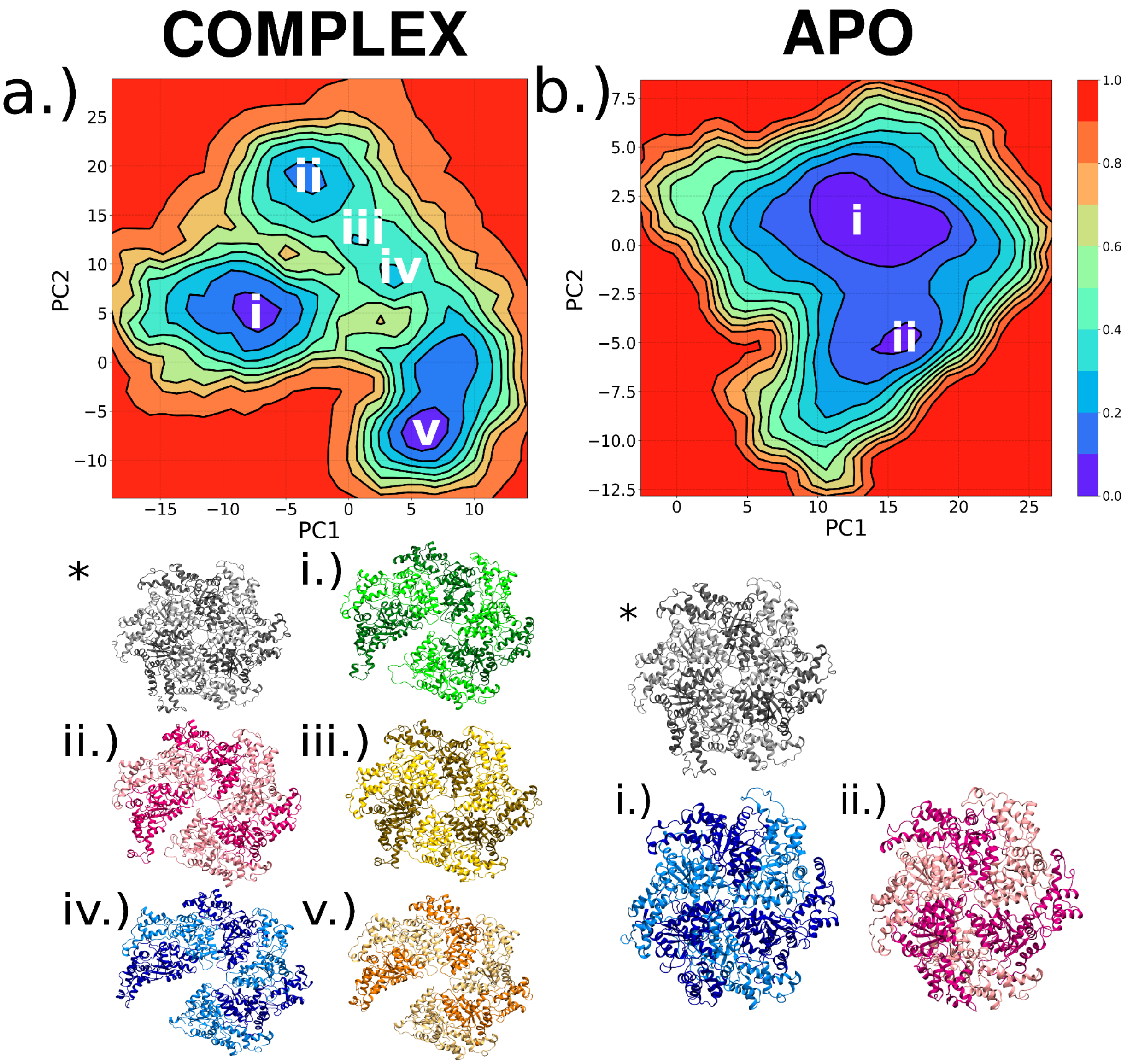


**Figure S9.** The free energy landscape in the PC1/PC2 subspace for the **R591S in WT space**. (a) For the COMPLEX state, we identified five minima and extracted the corresponding lowest-energy representative structures, labeled as i - v (b) For the APO state, we identified two minima and extracted the corresponding lowest-energy representative structures, labeled as i and ii. The starting structure (*) is shown in gray. The accompanying backbone-RMSD between the minima structures and the starting structure are provided in Table S14. Note that the actual range of free energy values for the COMPLEX state of R591S on WT space is 0–2.65 kcal/mol, and for the APO state of R591S on WT space, it is 0–3.49 kcal/mol. For visualization purposes, we have selected an arbitrary range of 0–1.

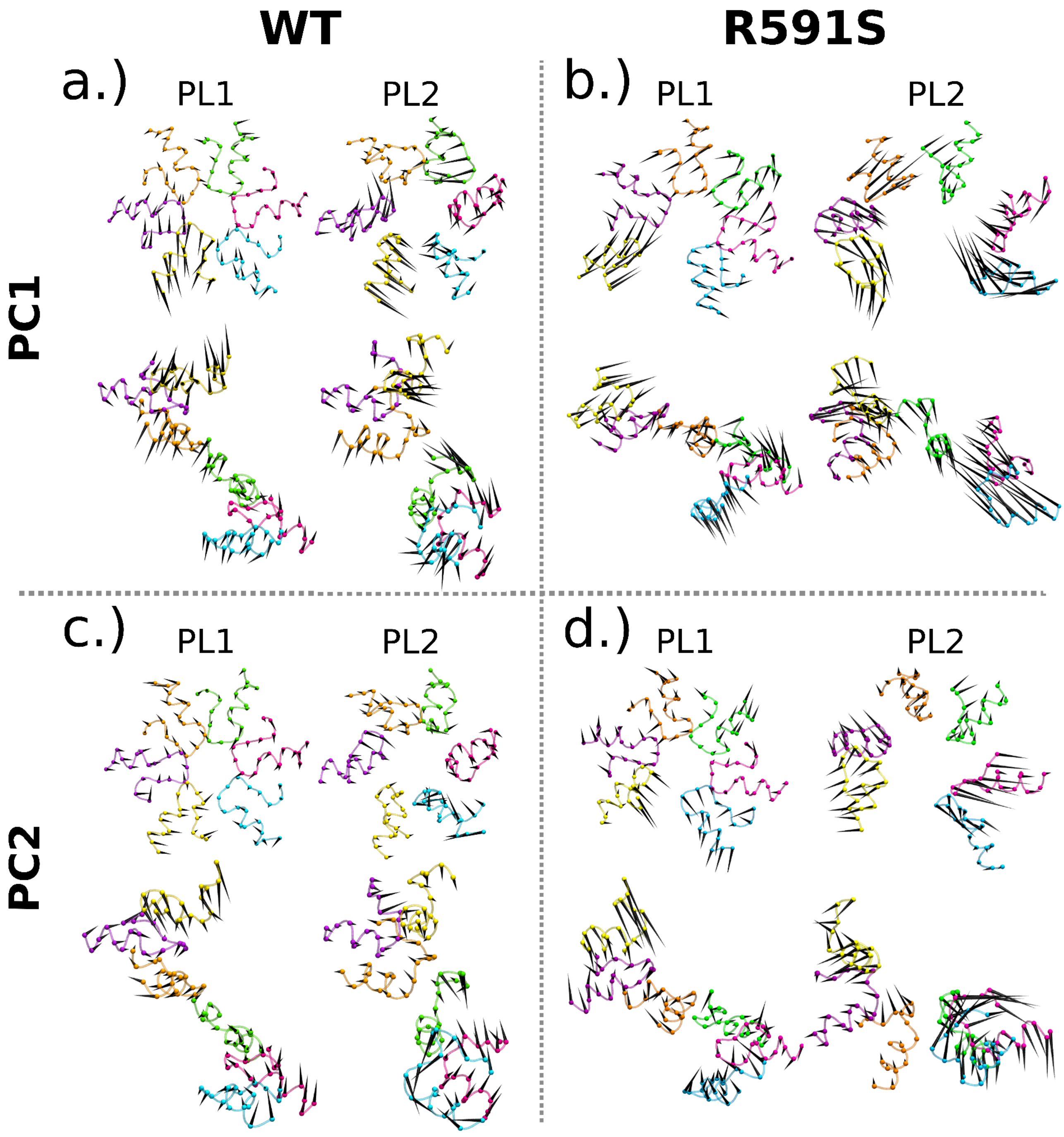


**Figure S10.** The porcupine plots for the extracted **PL motions** of PC1 and PC2 for the WT and R591S hexameric systems in the **COMPLEX** state. The variance covered in each system is provided in Table S9-S10.

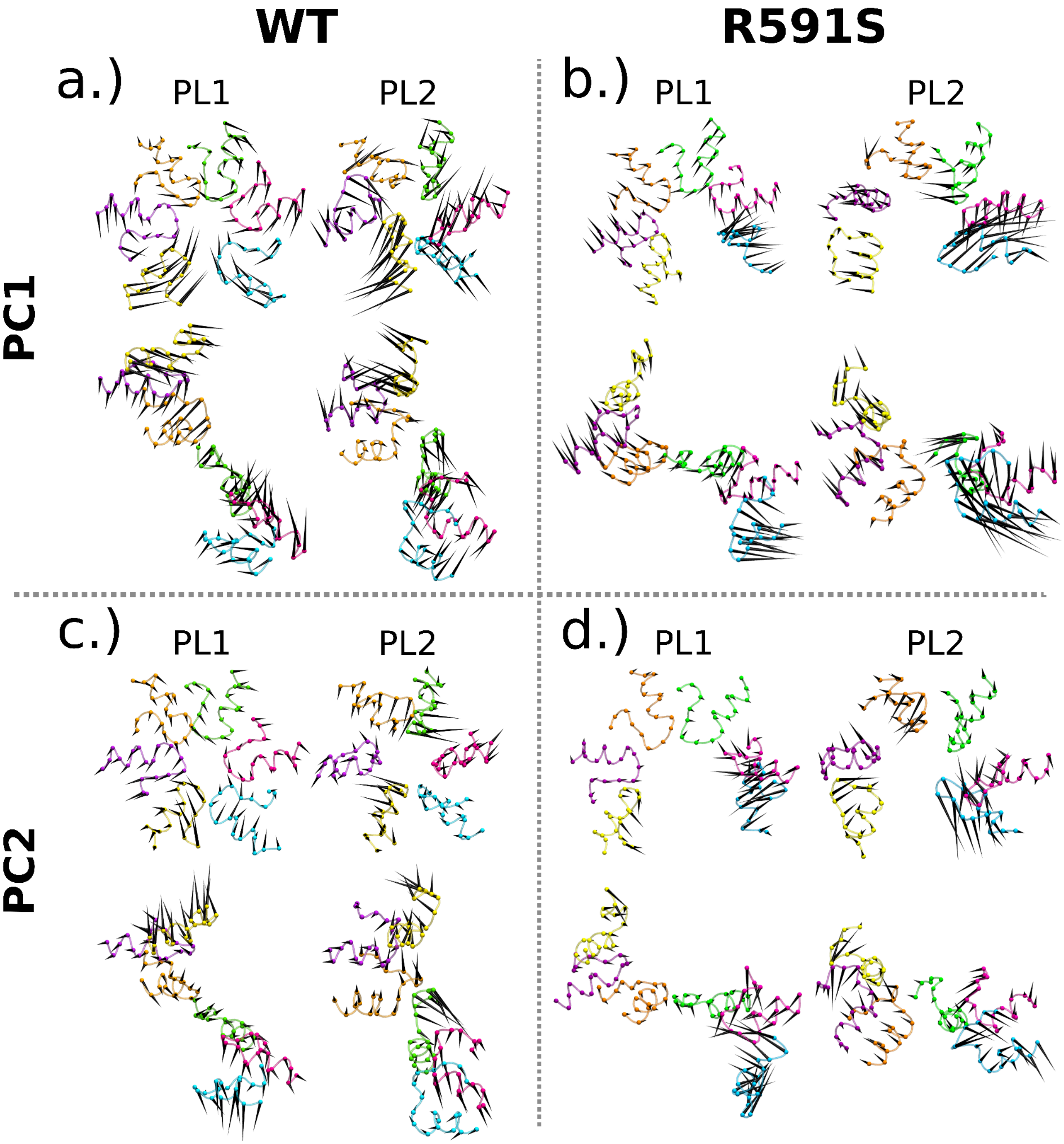


**Figure S11.** The porcupine plots for the extracted **PL motions** of PC1 and PC2 for the WT and R591S hexameric systems in the **APO** state. The variance covered in each system is provided in Table S9-S10.

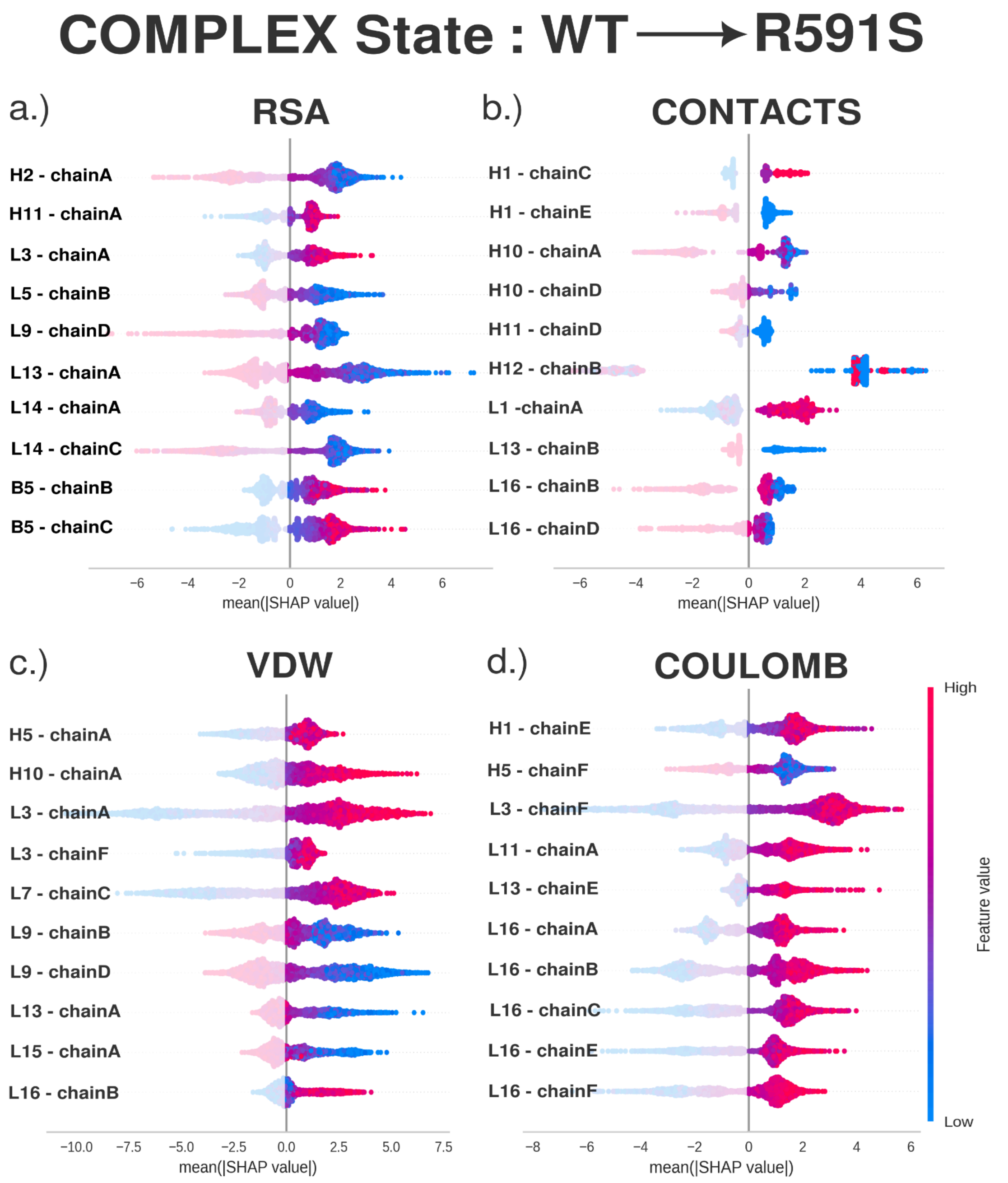


**Figure S12.** The SHAP beeswarm plots correspond to the **COMPLEX** state. Plots (a) - (d) shows the top 10 features that contributed to the model's classification as either WT or the R591S mutant for the four descriptors. Each point is colored based on how its descriptor value compares to the average, with magenta indicating an increase and blue indicating a decrease. The right side of each plot shows the observations with positive SHAP values that helped the model to classify the mutant system. Features found in at least two descriptors are listed in Table S19 and are depicted in Figure 5a.

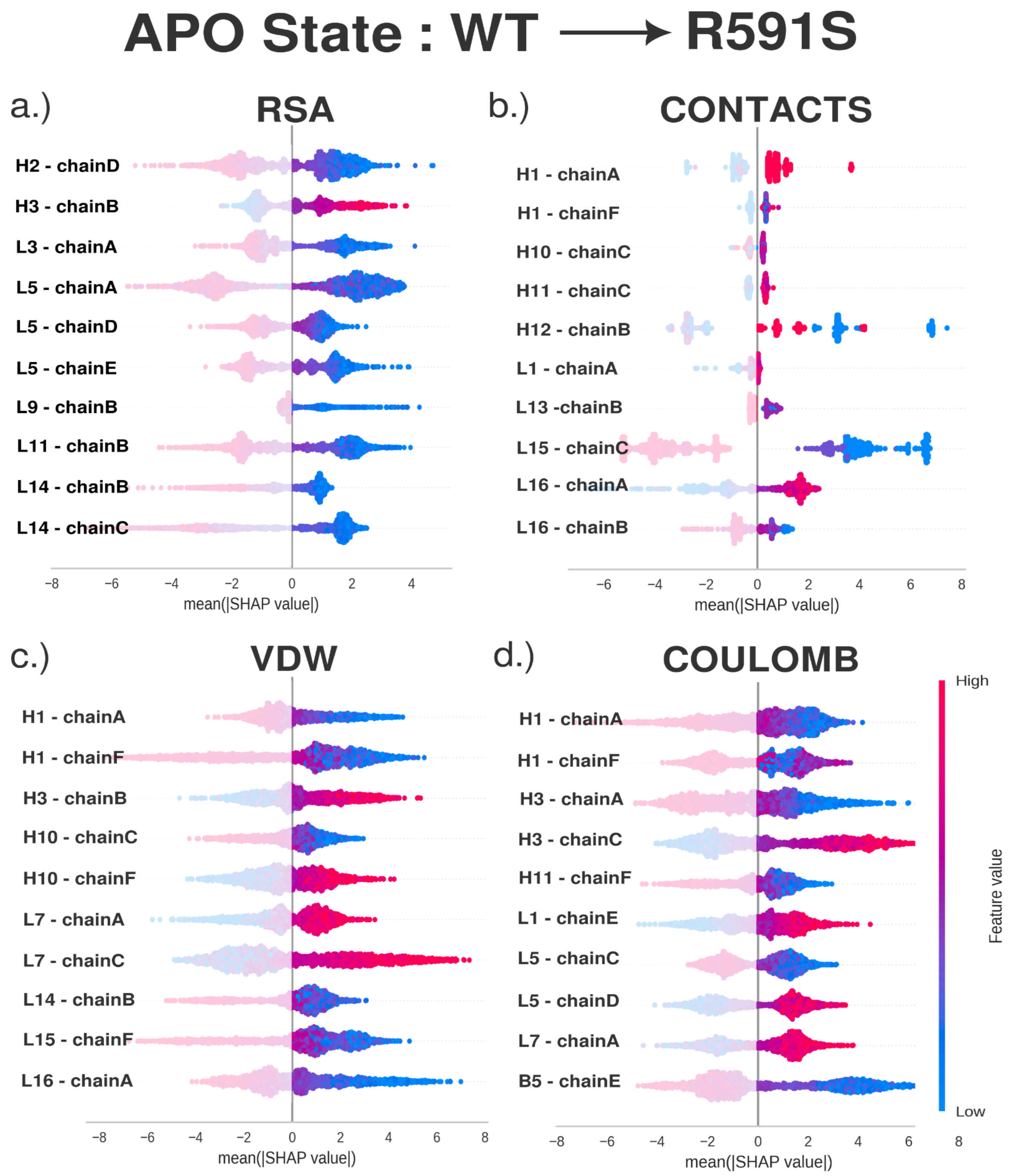


**Figure S13.** The SHAP beeswarm plots correspond to the **APO** state. Plots (a) - (d) shows the top 10 features that contributed to the model's classification as either WT or the R591S mutant for the four descriptors. Each point is colored based on how its descriptor value compares to the average, with magenta indicating an increase and blue indicating a decrease. The right side of each plot shows the observations with positive SHAP values that helped the model to classify the mutant system. Features found in at least two descriptors are listed in Table S19 and are depicted in Figure 5b.

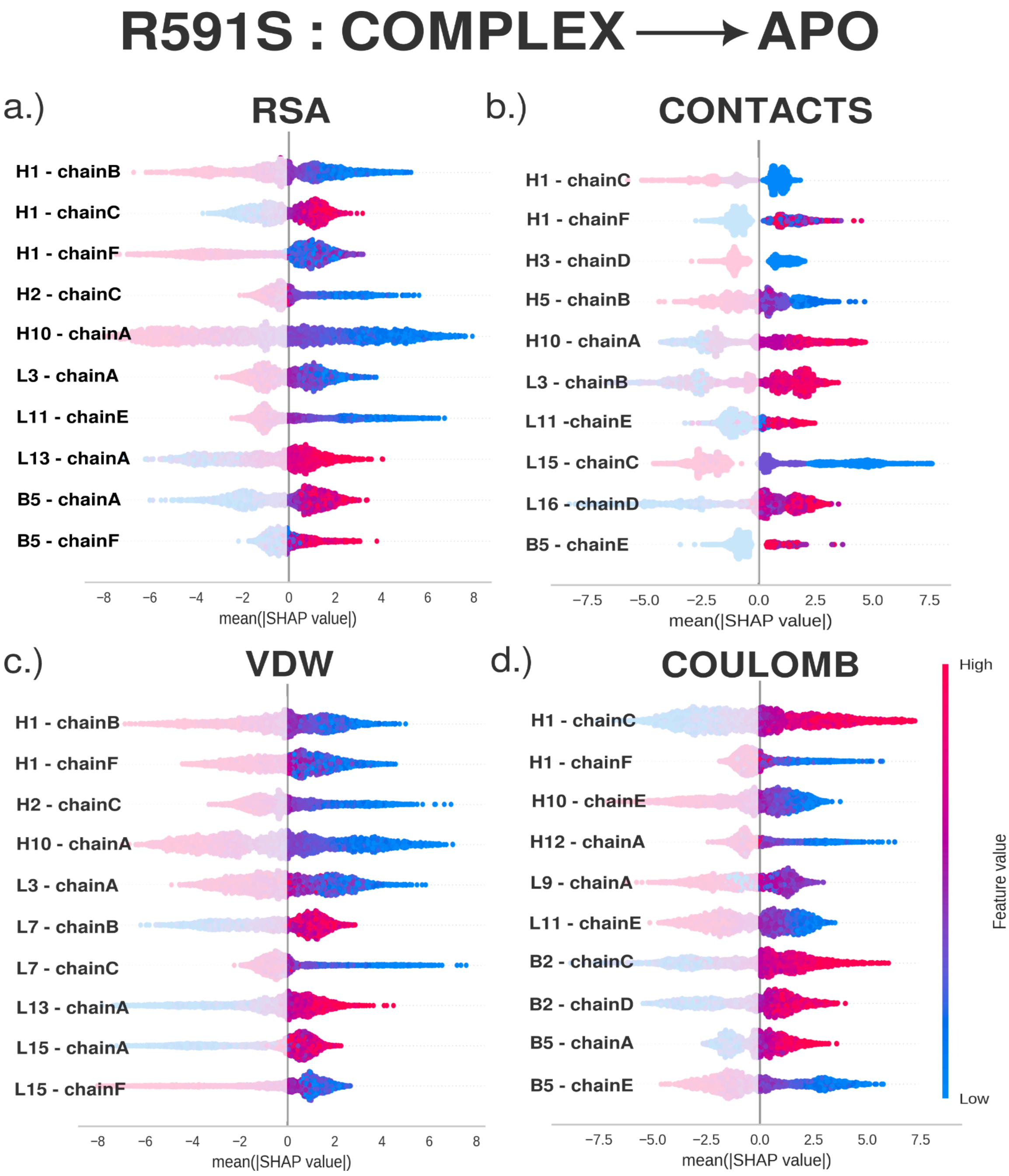


**Figure S14.** The SHAP beeswarm plots correspond to the **COMPLEX to APO transition of the R591S mutant system**. Plots (a) - (d) shows the top 10 features that contributed to the model's classification as either COMPLEX or APO for the four descriptors. Each point is colored based on how its descriptor value compares to the average, with magenta indicating an increase and blue indicating a decrease. The right side of each plot shows the observations with positive SHAP values that helped the model to classify the APO state. Features found in at least two descriptors are listed in Table S20 and are depicted in Figure 6b.

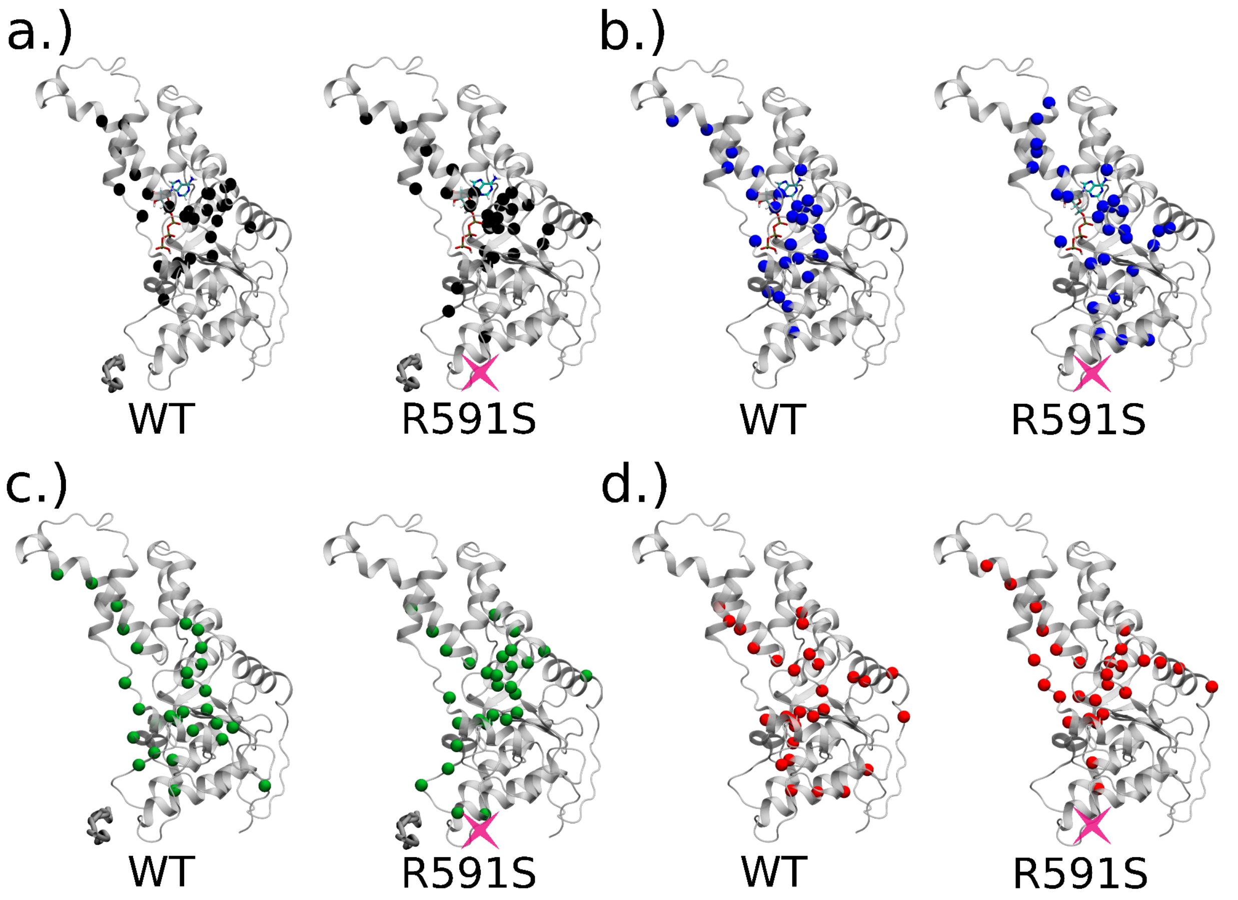


**Figure S15.** The positions of top 10% betweenness centrality residues from Tables S22 and S23 of the **WT** and the **R591S** mutant **monomeric** systems for the (a) COMPLEX, (b) NUCLEOTIDE, (c) SUBSTRATE, and (d) APO states. The position of the R591S mutation is indicated with a magenta “X”.

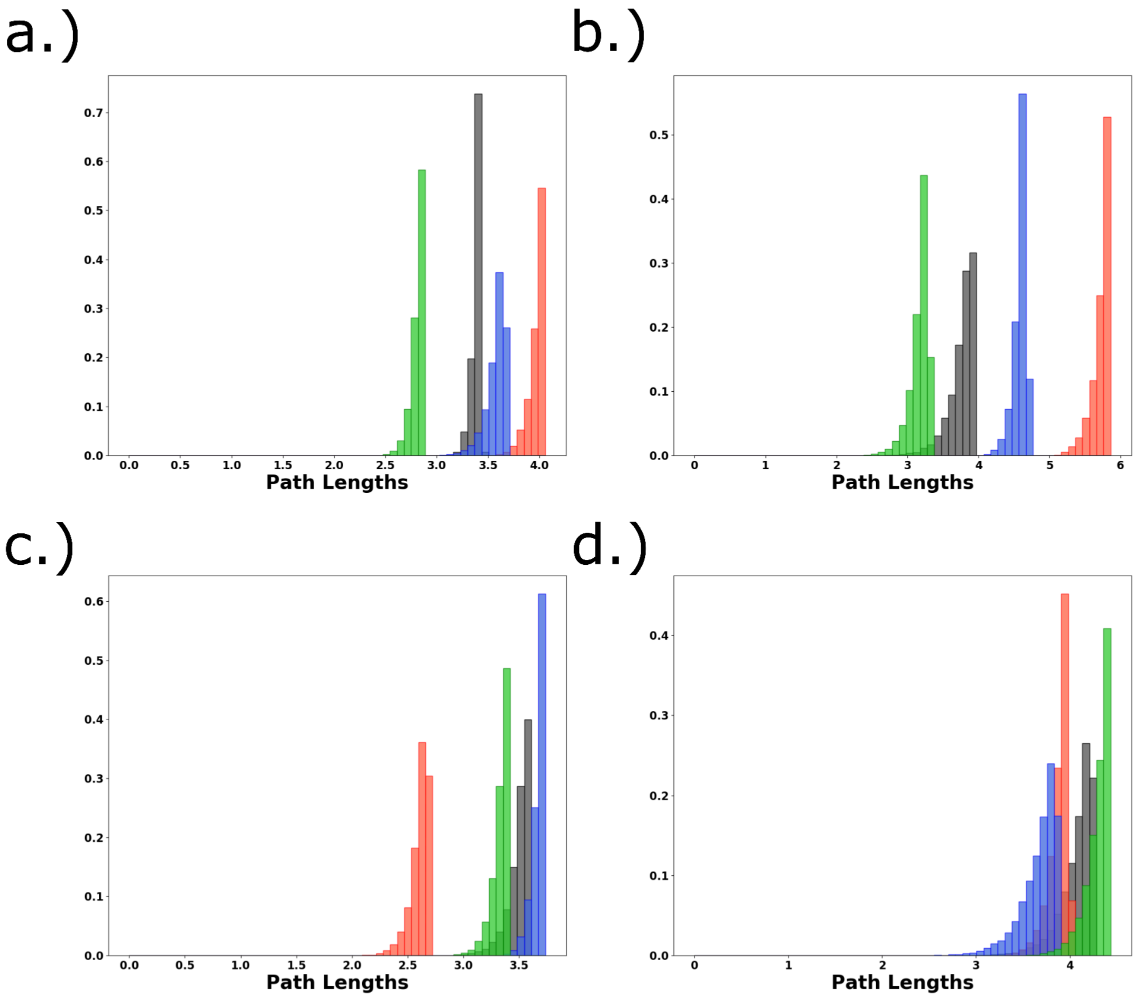


**Figure S16.** The suboptimal path lengths of the 80,000 paths collected for the shortest path pair for the monomeric systems between the **ATP binding region** and the **CTT binding region** in (a.) the **WT** and (c.) the **R591S** mutation systems; as well as between the **Allosteric Center** and the **CT Hlx** in the (b.) in **WT** and (d.) the **R591S** systems. This allows us to track the allosteric propagation through the NBD and between the NBD and HBD. The COMPLEX setup is shown in gray, the NUCLEOTIDE setup in blue, the SUBSTRATE setup in green, and the APO setup in red.

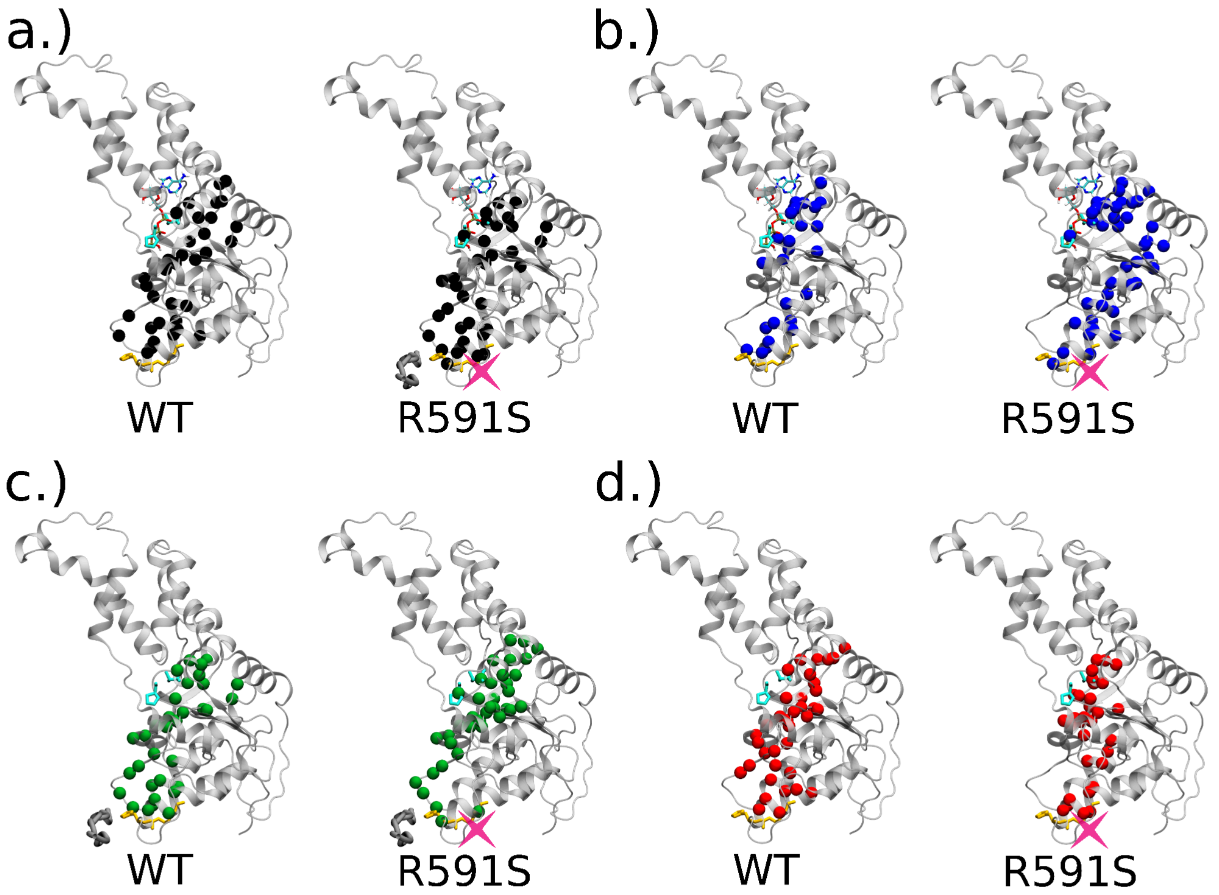


**Figure S17.** The positions found to be the most degenerate in 80,000 paths collected between the **ATP binding region** (teal) and the **CTT binding region** (gold) between the **WT** and **R591S** mutation **monomeric** systems for the (a) COMPLEX, (b) NUCLEOTIDE, (c) SUBSTRATE, and (d) APO states. The position of the R591S mutation is indicated with a magenta “X”. We find the largest differences due to the mutation of the allosteric center in the NUCLEOTIDE state. This describes the communication between the two main ligand binding regions within the NBD.

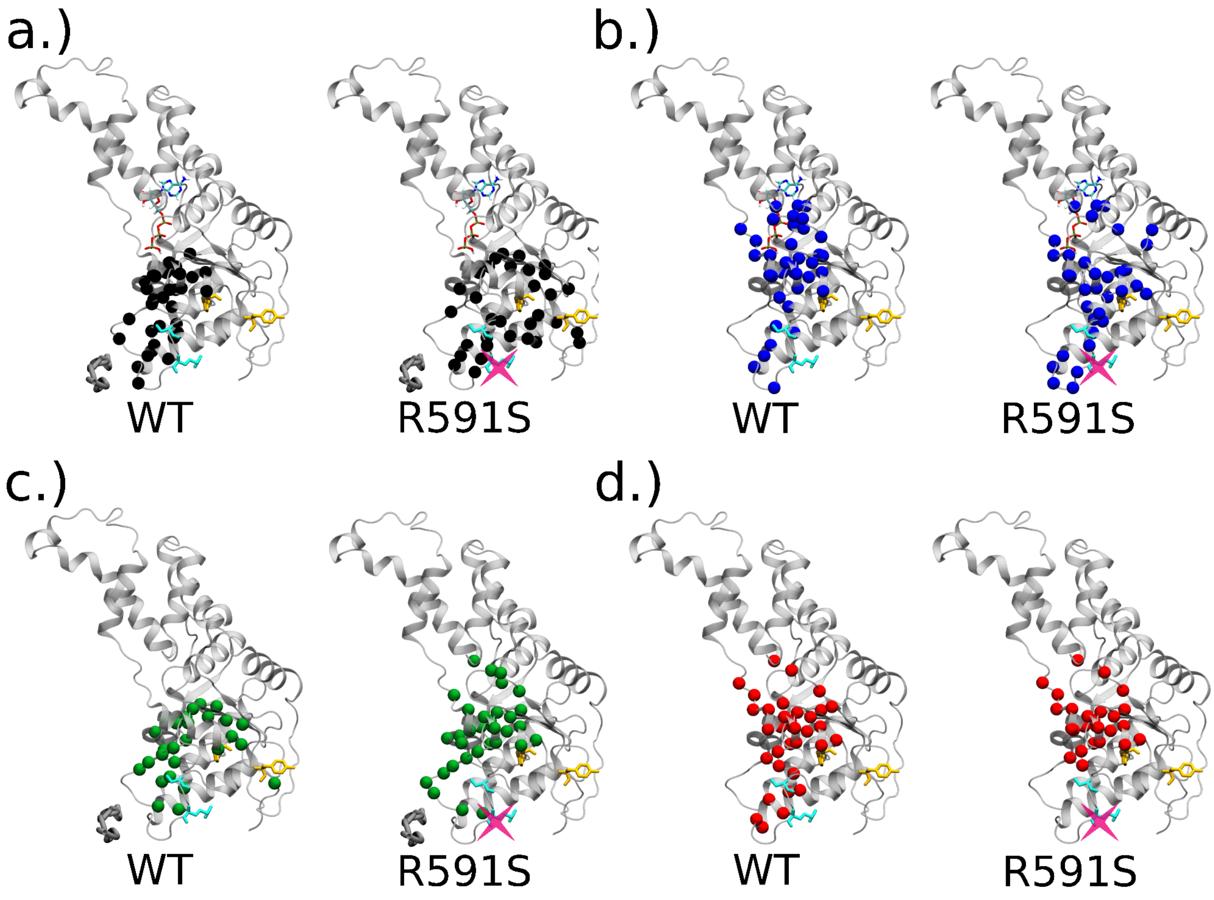


**Figure S18.** The positions found to be the most degenerate in 80,000 paths collected between the **Allosteric Center** (teal) and the **CT-Helix** (gold) between the **WT** and **R591S** mutation **monomeric** systems for the (a) COMPLEX, (b) NUCLEOTIDE, (c) SUBSTRATE, and (d) APO setups. The position of the R591S mutation is indicated with a magenta "X". This describes the communication between the NBD and the HBD which is observed to experience changes when binding one or both of the binding ligands.

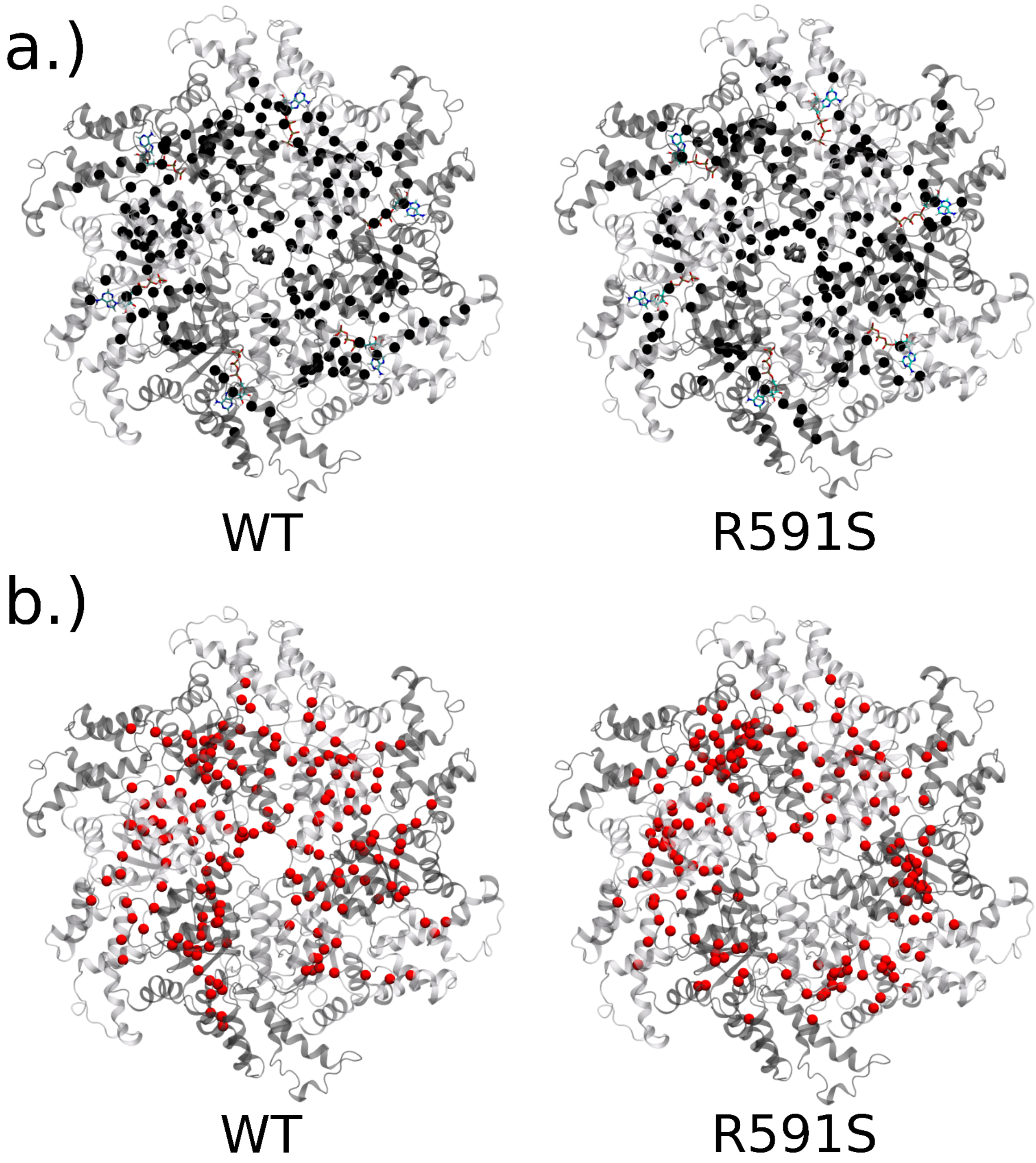


**Figure S19.** The positions of top 10% betweenness centrality residues for the **WT** from our previous publication[10] and the **R591S** mutation, as listed in Table S26 and S27, for the hexameric systems in the (a) COMPLEX and (b) APO states. We note the most significant changes are observed around the pore as well as in the terminal protomers (A & F) and their neighbors (B & E).

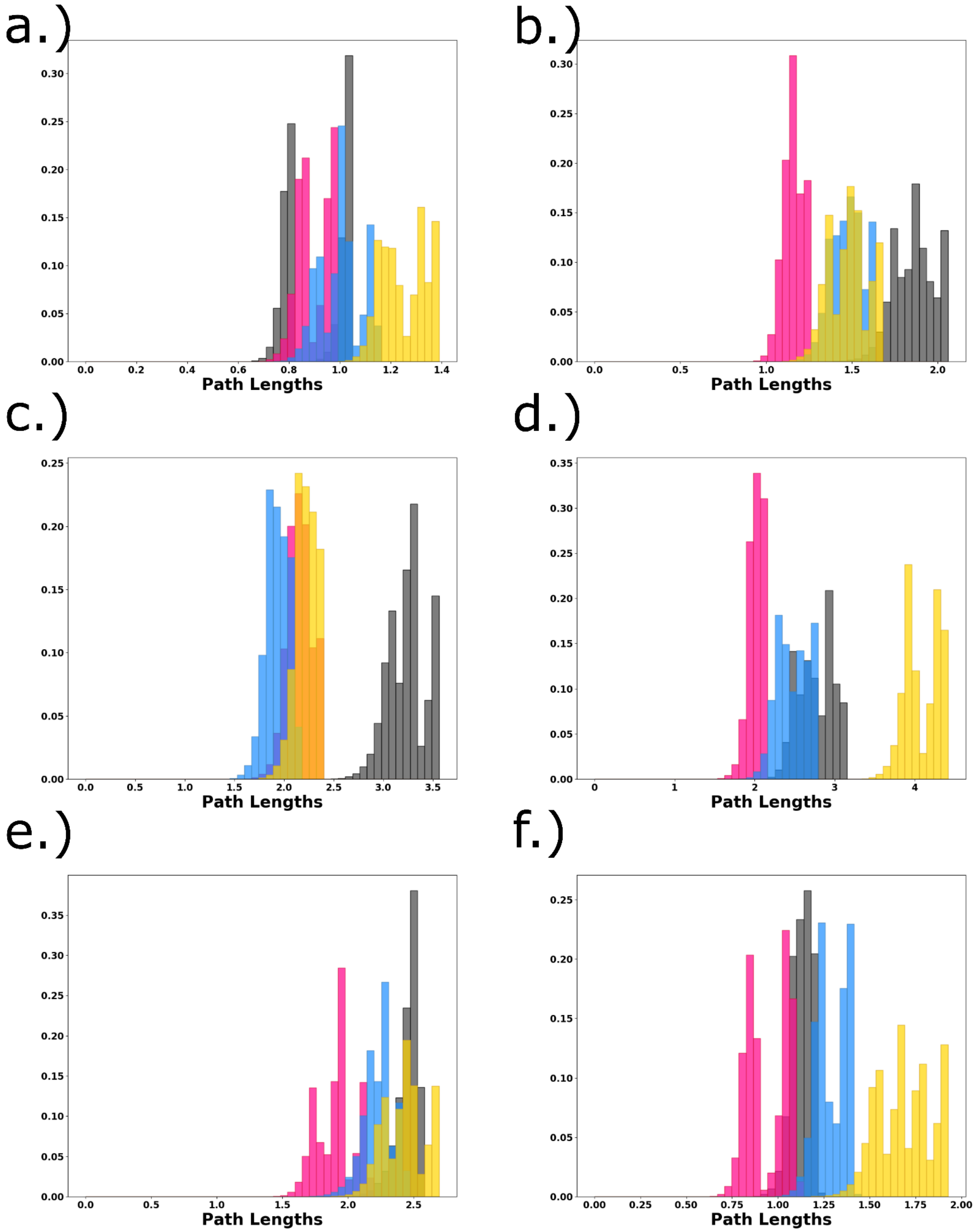


**Figure S20.** The suboptimal path lengths of the 80,000 **intra-protomer** paths collected between the ATP binding region and the CTT binding region (**intra-NBD**) between **WT** and **R591S** mutation **hexameric** systems. The COMPLEX setup is shown in gray for the WT and blue for R591S while the APO setup is shown in magenta for the WT and gold for R591S. Each protomer is labeled by the indicated letter.

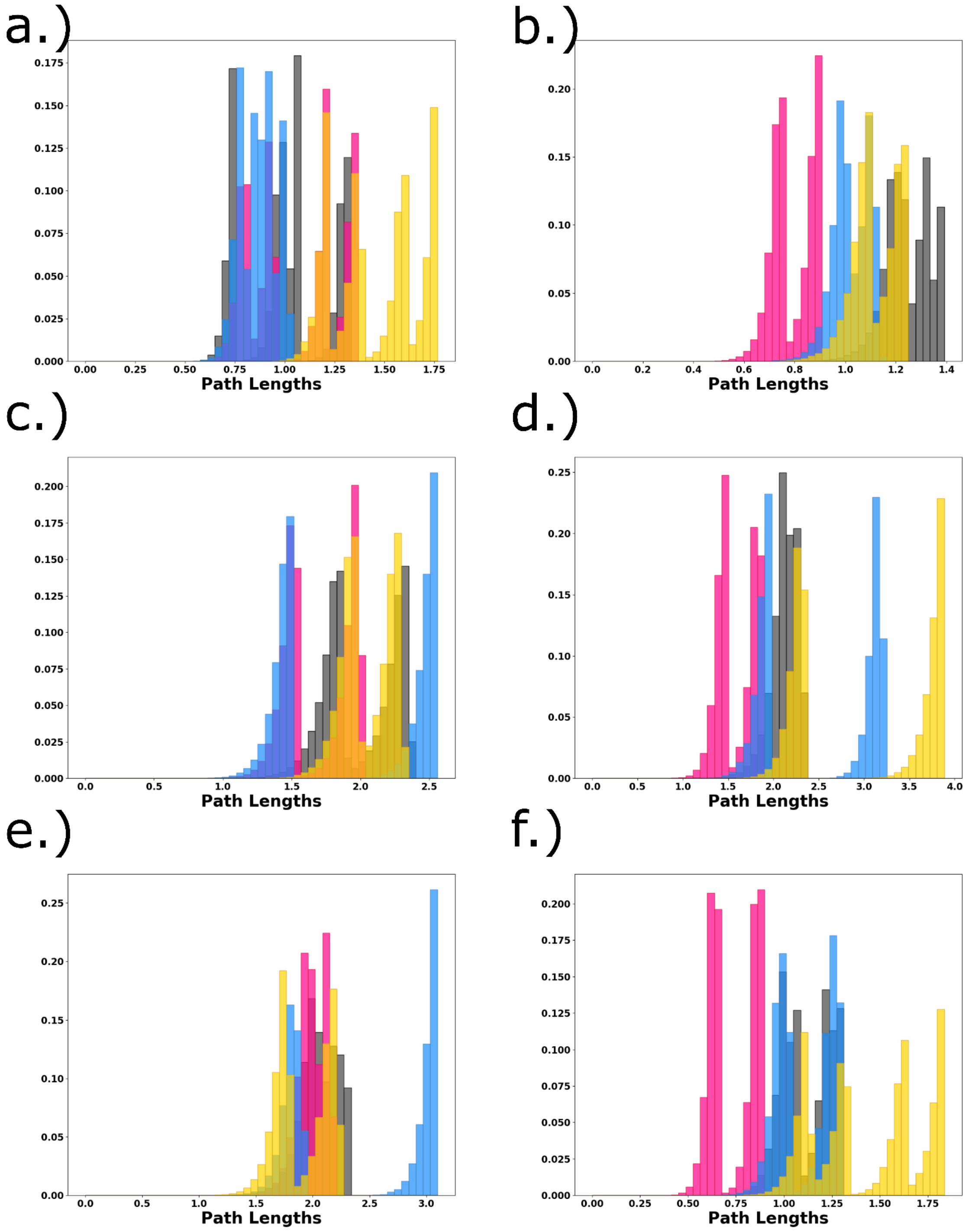


**Figure S21.** The suboptimal path lengths of the 80,000 **intra-protomer** paths collected between the Allosteric Center and the CT Hlx (**NBD-HBD**) between **WT** and **R591S** mutation **hexameric** systems. The COMPLEX setup is shown in gray for the WT and blue for R591S while the APO setup is shown in magenta for the WT and gold for R591S. Each protomer is labeled by the indicated letter.

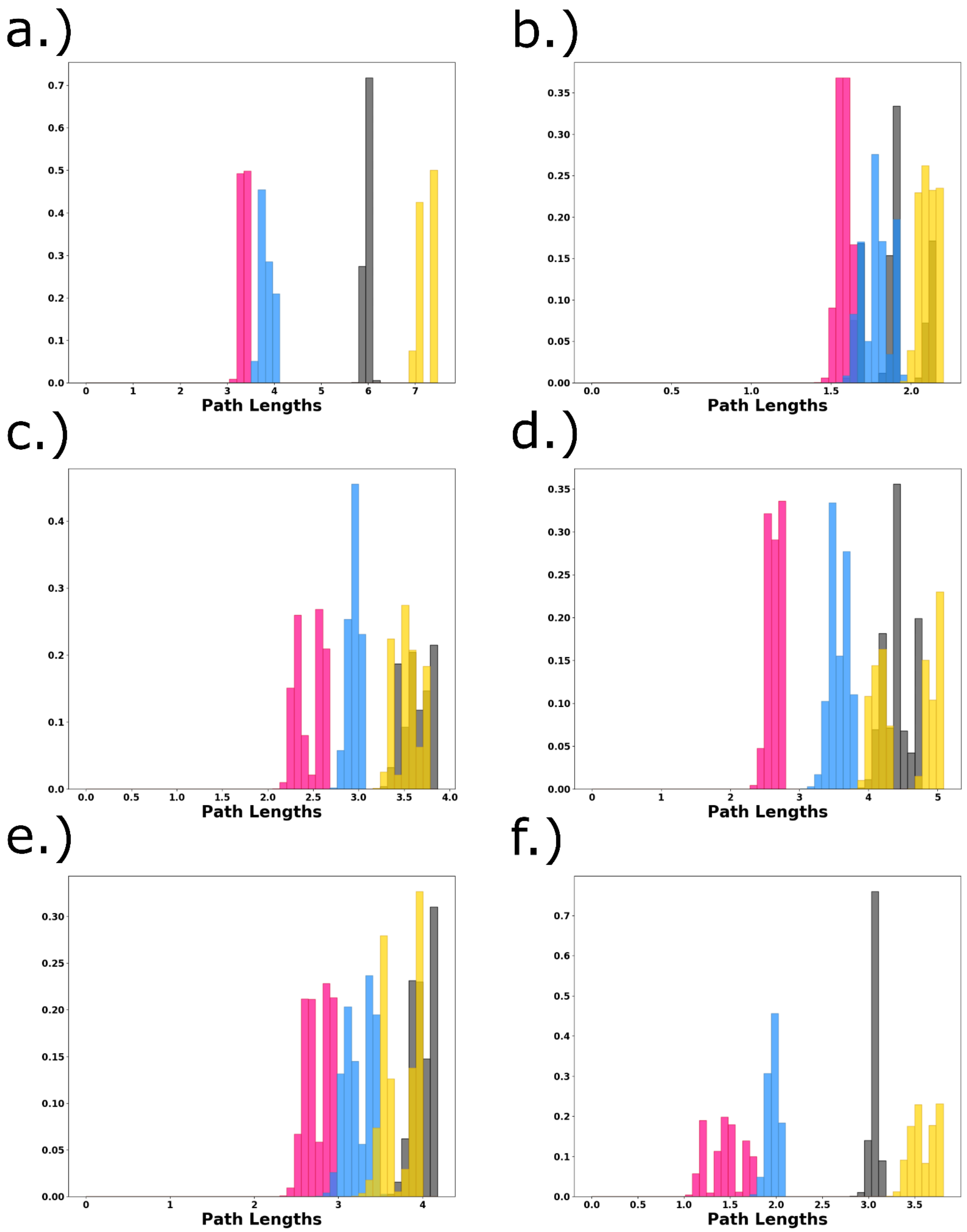


**Figure S22.** The suboptimal path lengths of the 80,000 **inter-protomer** paths collected between the ATP binding region in protomer i and the CTT binding region (**inter-protomer NBD**) in protomer i-1 for the **WT** and the **R591S** mutant **hexameric** systems. The COMPLEX setup is shown in gray for the WT and blue for R591S while the APO setup is shown in magenta for the WT and gold for R591S. a.) source in protomer A to sink in protomer F, b.) B to A, c.) C to B, d.) D to C, e.) E to D and f.) F to E.

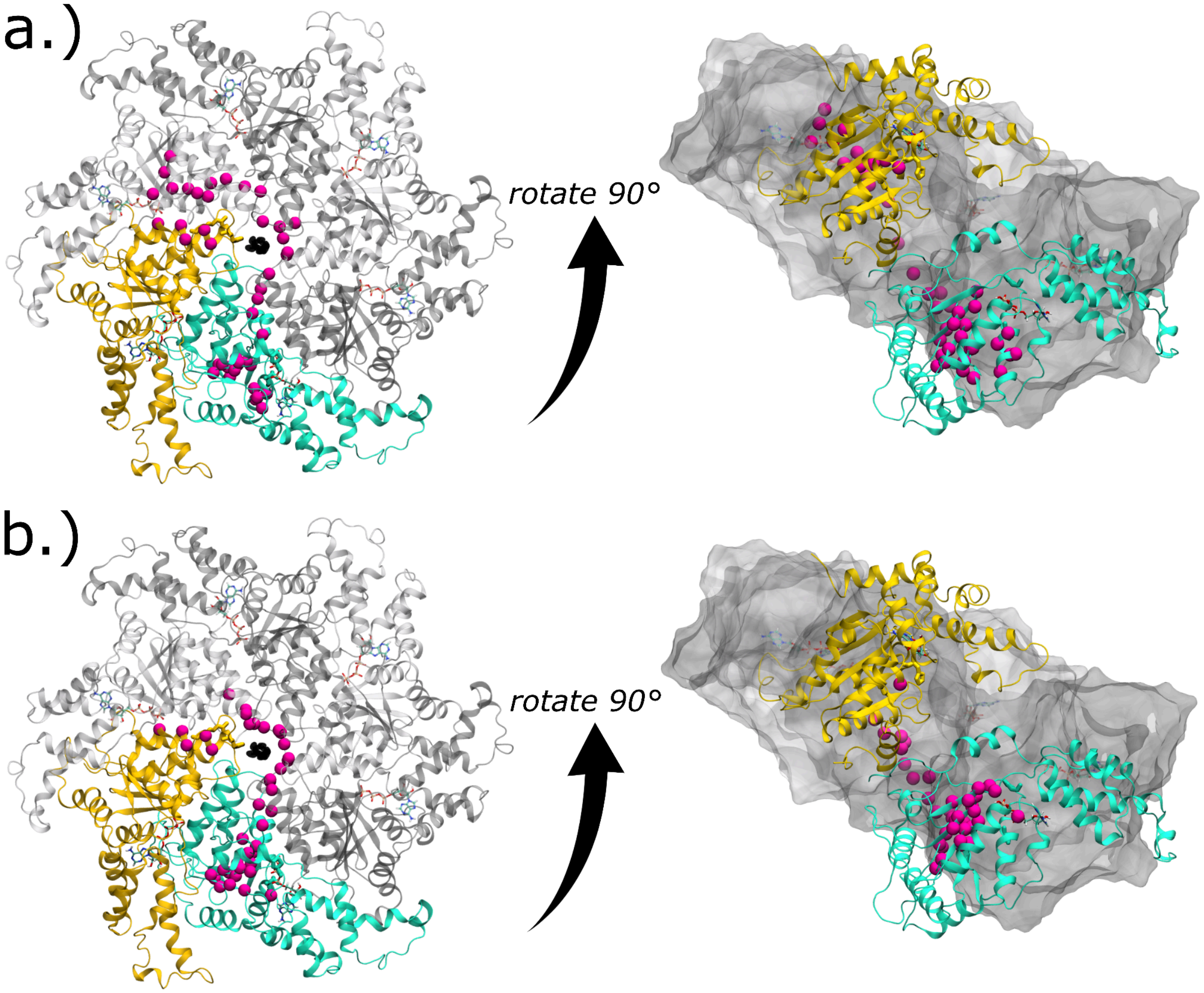


**Figure S23.** The positions found to be the most degenerate in the 80,000 **A to F** paths collected between the ATP binding region (teal) and the CTT binding region (gold) (**inter-protomer NBD**) between the (a) **WT** and (b) **R591S** mutation **hexameric** systems for the **COMPLEX** setups. This describes the communication between the terminal protomers which is observed to experience dramatic changes due to the mutation.

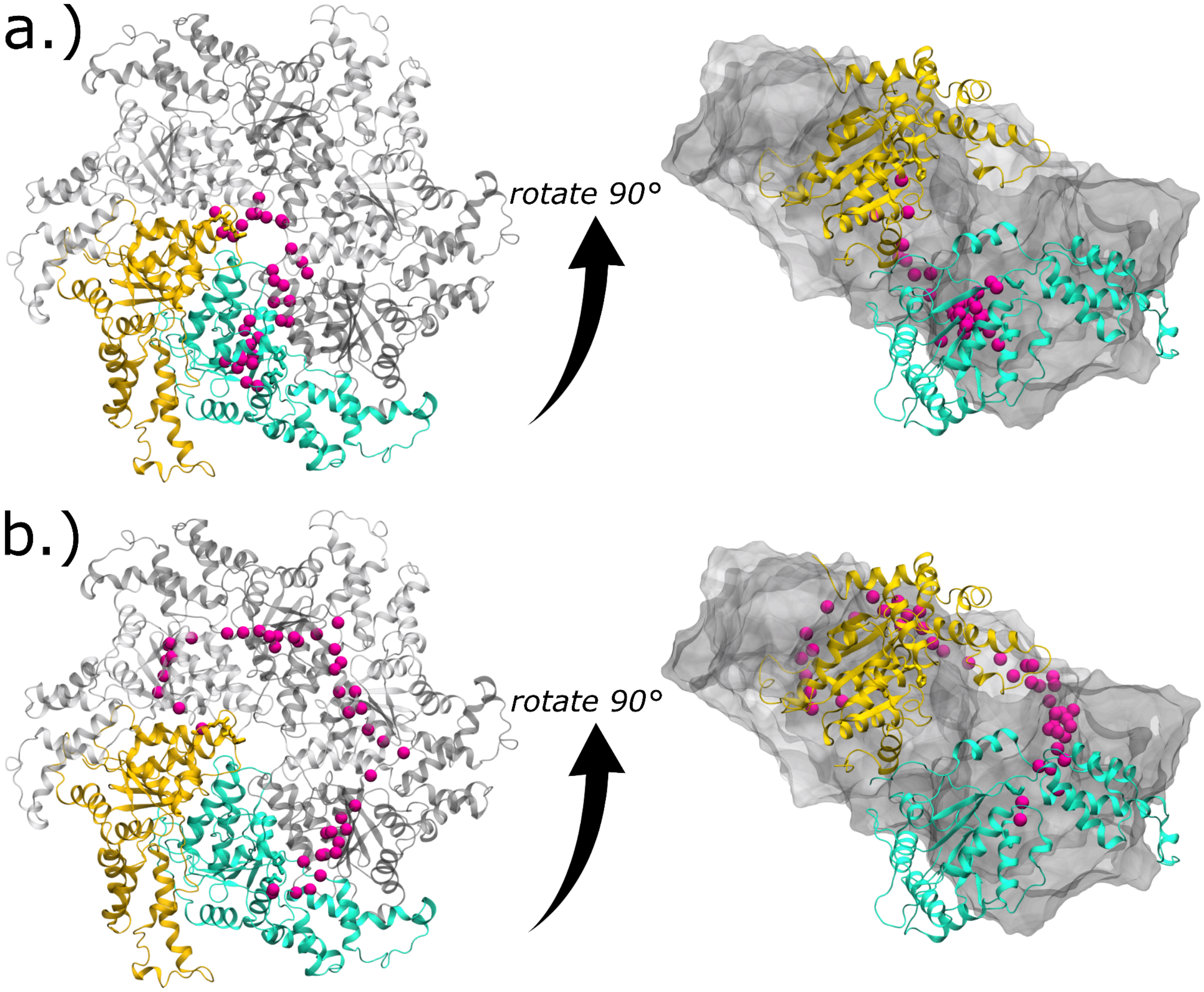


**Figure S24.** The positions found to be the most degenerate in the 80,000 **A to F** paths collected between the ATP binding region (teal) and the CTT binding region (gold) (**inter-protomer NBD**) between the (a) **WT** and (b) **R591S** mutation **hexameric** systems for the **APO** setups. This describes the communication between the terminal protomers which is observed to experience dramatic changes due to the mutation.

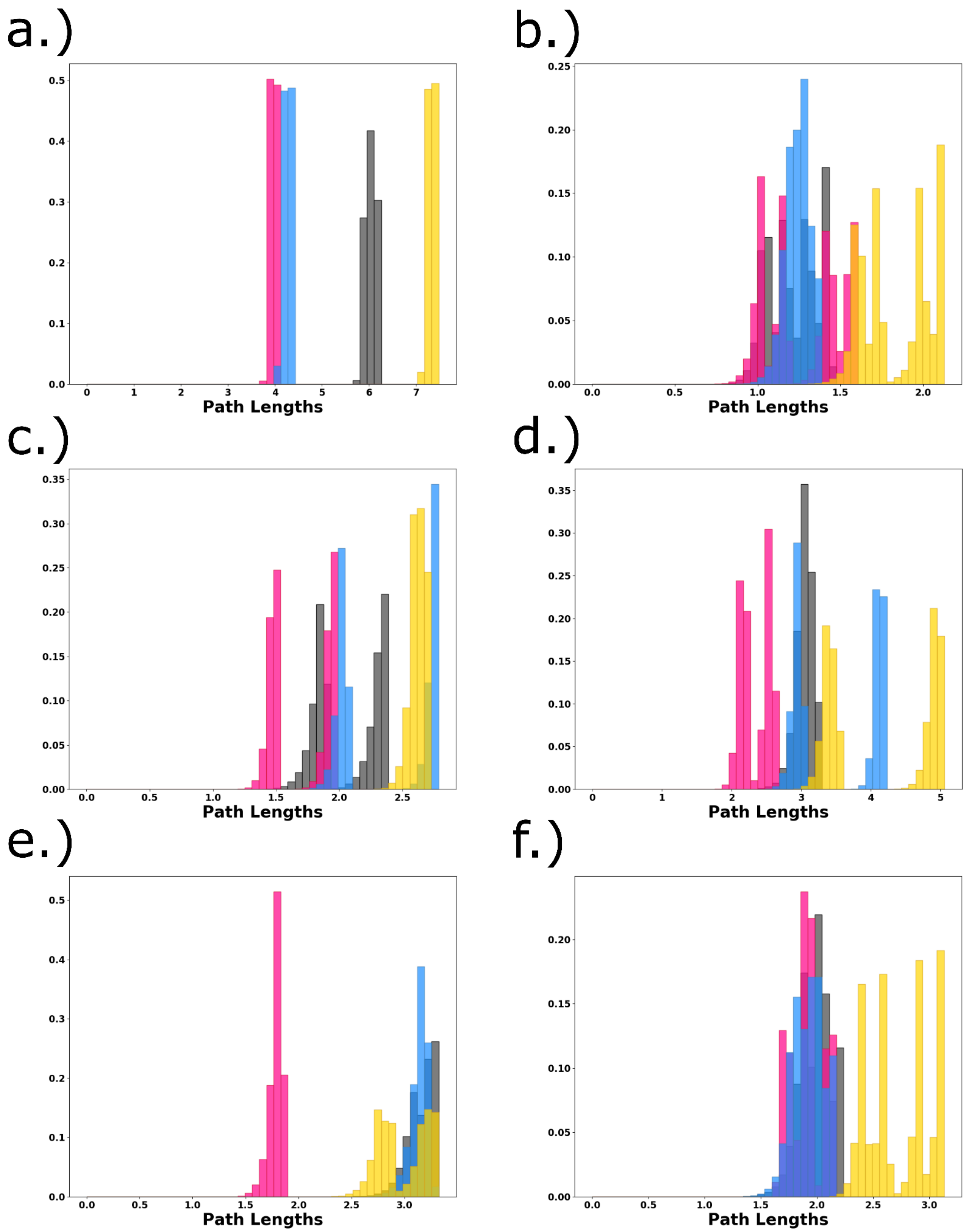


**Figure S25.** The suboptimal path lengths of the 80,000 **inter-protomer** paths collected between the Allosteric Center in protomer i and the CT Hlx (**NBD i - HBD i-1**) in protomer i-1 between **WT** and **R591S** mutation **hexameric** systems. The COMPLEX setup is shown in gray for the WT and blue for R591S while the APO setup is shown in magenta for the WT and gold for R591S. a.) source in protomer A to sink in protomer F, b.) B to A, c.) C to B, d.) D to C, e.) E to D and f.) F to E.

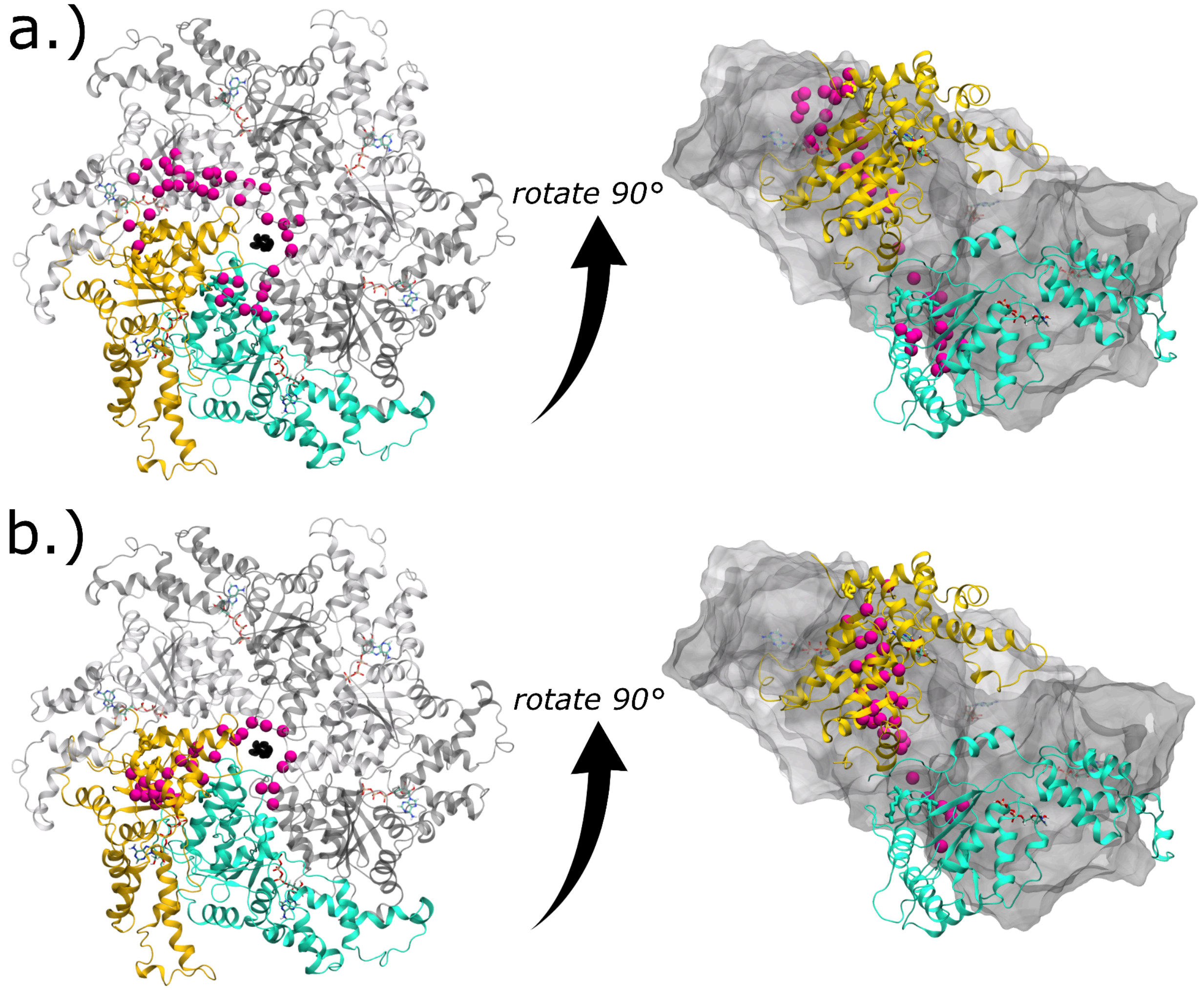


**Figure S26.** The positions found to be the most degenerate in the 80,000 A to F paths collected between the Allosteric Center (teal) and the CT Hlx (gold) (**NBD i - HBD i-1**) between the (a) **WT** and (b) **R591S** mutation **hexameric** systems for the **COMPLEX** setups. This describes the communication between the terminal protomers which is observed to experience dramatic changes due to the mutation.

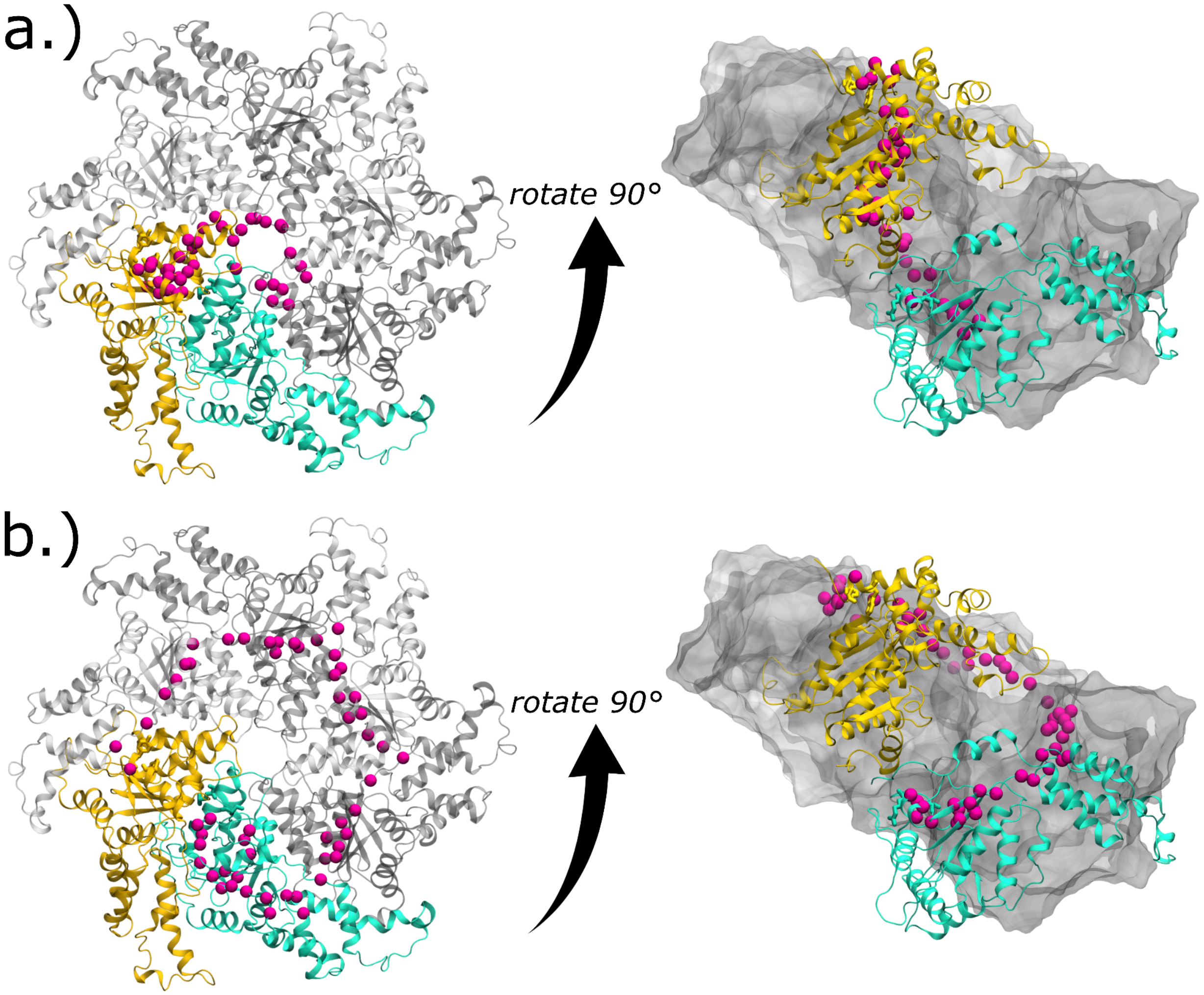


**Figure S27.** The positions found to be the most degenerate in the 80,000 **A to F** paths collected between the Allosteric Center (teal) and the CT Hlx (gold) (**NBD i - HBD i-1**) between the (a) **WT** and (b) **R591S** mutation **hexameric** systems for the **APO**setups. This describes the communication between the terminal protomers which is observed to experience dramatic changes due to the mutation.
